# Transitions in dynamical regime and neural mode underlie perceptual decision-making

**DOI:** 10.1101/2023.10.15.562427

**Authors:** Thomas Zhihao Luo, Timothy Doyeon Kim, Diksha Gupta, Adrian G. Bondy, Charles D. Kopec, Verity A. Elliot, Brian DePasquale, Carlos D. Brody

**Author notes:** Correspondence should be addressed to: T.Z.L., T.D.K., or C.D.B. equal contribution (Thomas Zhihao Luo, Timothy Doyeon Kim).

## Abstract

Perceptual decision-making is the process by which an animal uses sensory stimuli to choose an action or mental proposition. This process is thought to be mediated by neurons organized as attractor networks^1,2^. However, whether attractor dynamics underlie decision behavior and the complex neuronal responses remains unclear. Here we use simultaneous recordings from hundreds of neurons, together with an unsupervised, deep learning-based method, to discover decision-related neural dynamics in frontal cortex and striatum of rats while the subjects accumulate pulsatile auditory evidence. We found that trajectories evolved along two sequential regimes, the first dominated by sensory inputs, and the second dominated by the autonomous dynamics, with flow in a direction (i.e., “neural mode”) largely orthogonal to that in the first regime. We propose that the transition to the second regime corresponds to the moment of decision commitment. We developed a simplified model that approximates the coupled transition in dynamics and neural mode and allows precise inference, from each trial’s large-scale neural population activity, of a putative neurally-inferred time of commitment (“nTc”) on that trial. The simplified model captures diverse and complex single-neuron temporal profiles, such as ramping and stepping^3–5^, as well as trial-averaged curved trajectories^6–8^, and reveals distinctions between brain regions. The estimated nTc times were not time-locked to stimulus onset or offset, or to response onset, but were instead broadly distributed across trials. If nTc marks the moment of decision commitment, sensory evidence before nTc should affect the decision, while evidence afterward should not. Behavioral analysis of trials aligned to their estimated nTc times confirmed this prediction. Our results show that the formation of a perceptual choice involves a rapid, coordinated transition in both the dynamical regime and the neural mode of the decision process that corresponds to commitment to a decision, and suggest this moment as a useful entry point for dissecting mechanisms underlying rapid changes in internal state.

---

Theories of attractor dynamics have been successful at capturing multiple brain functions^9^, including motor planning^10^ and neural representations of space^11,12^. Attractors are a set of states toward which a system tends to evolve, from a variety of starting positions. In these theories, the computations of a brain function are carried out by the temporal evolution, or the dynamics, of the system. Experimental findings support the idea that the brain uses systems with attractor states for computations underlying working memory^10^ and navigation^11^. These theories often focus on the low-dimensional nature of neural population activity^6,7,13,14^, and account for the responses across a large number of neurons using a dynamical system model whose variable has only a few dimensions^15,16,11,17^.

Attractor network models have also been hypothesized to underlie perceptual decision-making: the process through which noisy sensory stimuli are categorized to select an action or mental proposition. In these hypotheses, the network dynamics carry out the computations needed in decision formation^1,6,18–26^, such as accumulating sensory evidence and committing to a choice. While some experimental evidence favors a role of attractors in perceptual decisions^10,27,28^, the actual population-level dynamics underlying decision-making have not been directly estimated. Knowledge of these dynamics would directly test the current prevailing attractor hypotheses, provide fundamental constraints on neural circuit models, and account for the often complex temporal profiles of neural activities.

A separate line of work involves tools, sometimes based on deep learning, for discovering, in a data-driven manner, the low-dimensional component of neural activity^14,29,30^. In this approach, the spike trains of many simultaneously recorded neurons are modeled as being a function of a few “latent” variables that are shared across neurons.

To combine both lines of work, we used a novel method^31^ that estimates, from the spike trains of simultaneously recorded neurons, the dynamics of a low-dimensional variable ***z***, given by:

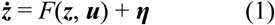

where ***u*** are external inputs, ***η*** is noise, and, when applied to perceptual decisions, ***z*** represents the dynamical state of the brain’s decision process at a given time (**Fig**. 1a-c). The instantaneous change of the decision variable, or its dynamics, is given by ***ż***, which depends on ***z*** itself, as well as ***u*** and ***η***. This approach aims to estimate the function *F*, and through it, capture the presence and nature of attractors in the neural dynamics.

**Figure 1.**
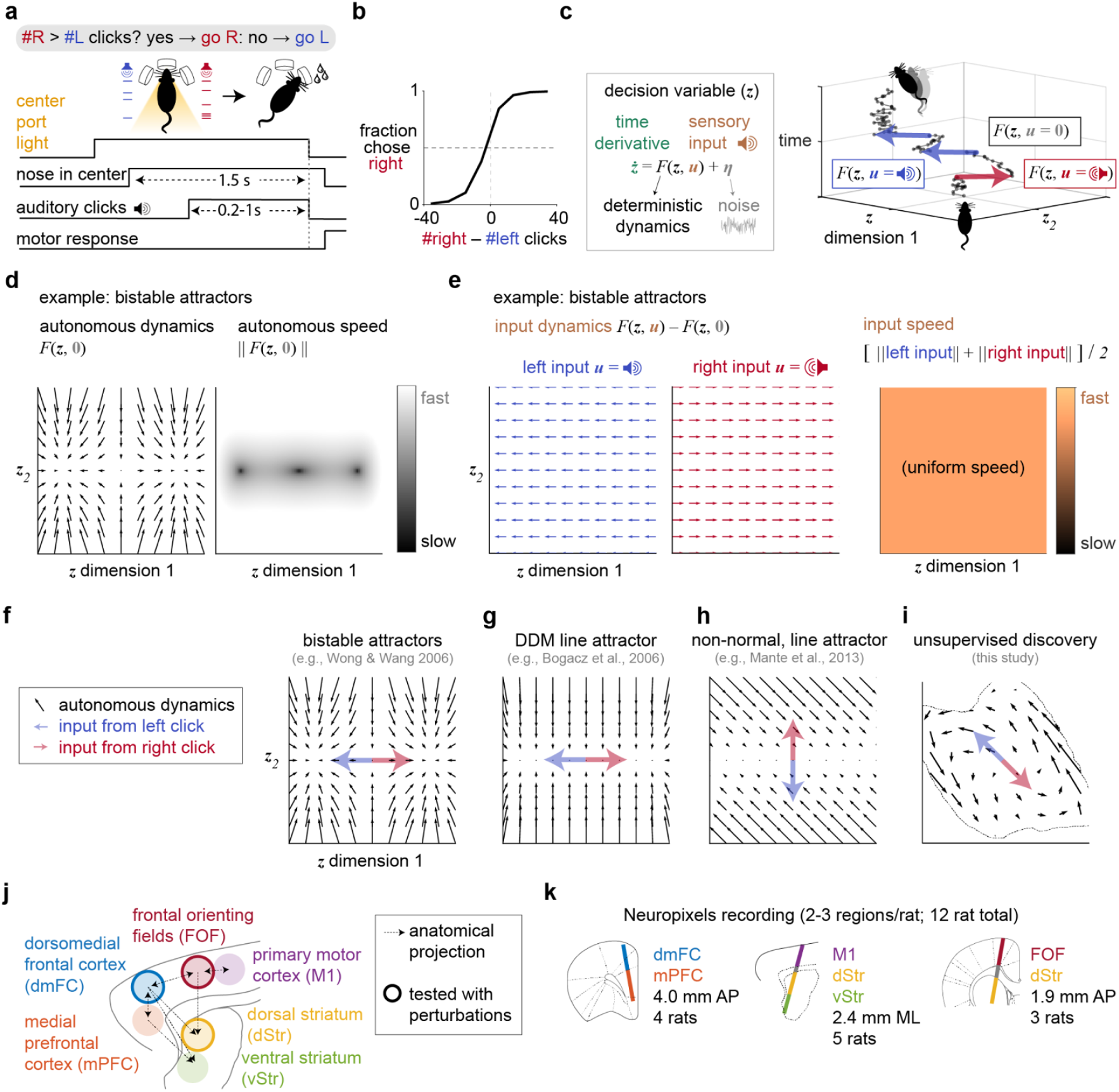
Attractor models of decision-making were tested by recording from rat frontal cortex and striatum. **a**, Rats were trained to accumulate auditory pulsatile evidence over time. While keeping its head stationary, the rat listened to randomly timed clicks played from loudspeakers on its left and right. At the end of the stimulus, the rat receives a water reward for turning to the side with more clicks. The earliest time when a rat can respond is fixed to be 1.5 s relative to the moment of inserting its nose in the center port (i.e., not a reaction time paradigm). **b**, Behavioral performance in an example recording session. **c**, The decision process is modeled as a dynamical system. Right: the blue, red, and black lines represent the change in the decision variable in the presence of left, right, or no click, respectively. **d**, Autonomous dynamics illustrated using the bistable attractors hypothesis. In the velocity vector field (i.e., flow field; left), the arrow at each value of the decision variable ***z*** indicates how the instantaneous change depends on ***z*** itself. The arrow’s orientation represents the direction of the change, and its size the speed, which is also quantified using a heat map (right). **e**, Changes in ***z*** driven solely by the external sensory inputs. **f**, Bistable attractors hypothesis of decision-making, with the directions of the input dynamics. **g**, A hypothesis supposing a line attractor in the autonomous dynamics, based on the drift-diffusion model (DDM) of decision behavior. **h**, Recurrent neural networks (RNN) can be trained to make perceptual decisions using a line attractor that is not aligned to the input dynamics. **i**, Unsupervised discovery of dynamics that have not been previously hypothesized. **j**, Six interconnected frontal cortical and striatal regions are examined here. **k**, Neuropixels recordings (318±147 neurons/session/probe, mean±STD) from twelve rats.

## Autonomous and input dynamics differentiate hypotheses

The deterministic dynamics *F* (that is, in the absence of ***η***) are useful for distinguishing among attractor hypotheses of decision-making. *F* can be dissected into two components: the autonomous dynamics and the input-driven dynamics. Autonomous dynamics are the dynamics in the absence of sensory inputs ***u*** (i.e., *F*(***z*, 0**); **Fig**. 1d; Extended Data Fig. 1a-b). The input dynamics are the changes in ***z*** driven by sensory inputs ***u***, which can be distinguished from the autonomous dynamics as *F*(***z, u***) − *F*(***z*, 0**). The input dynamics can depend on ***z*** (**Fig**. 1e; Extended Data Fig. 1c-e).

Many of the currently prevailing neural attractor hypotheses of decision-making have been inspired by a classic and successful behavioral level model, the drift-diffusion model (DDM)^32–34^. In the behavioral DDM, a scalar (i.e., one-dimensional) decision variable *a* is driven by sensory evidence inputs (Extended Data Fig. 8a-b). For example, for decisions between Go-Right versus Go-Left, momentary evidence for Right (Left) might drive *a* in a positive (negative) direction. Through these inputs, the momentary evidence is accumulated over time in *a* until the value of *a* reaches an absorbing bound, a moment thought to correspond to decision commitment, and after which inputs no longer affect *a*. Different bounds correspond to different choice options: a positive (negative) bound would correspond to the decision to Go-Right (Go-Left). A straightforward implementation of the DDM in neural population dynamics, which we will refer to as the “DDM line attractor,” would posit a line attractor in neural space, with position of the neural state ***z*** along that line representing the value of *a*, and two point attractors at the ends of the line to represent the decision commitment bounds (**Fig**. 1g)^35^. Another hypothesis approximates the DDM process using bistable attractors^1^, with each of the two attractors representing each of the decision bounds, and in between the two attractors, a one-dimensional stable manifold of slow autonomous dynamics that corresponds to the evidence accumulation regime (**Fig**. 1f). In both the DDM line attractor and bistable attractor hypotheses, differential evidence inputs are aligned with the slow dynamics manifold and the attractors at its endpoints. A third hypothesis, inspired by trained recurrent neural networks (RNNs), also posits a line attractor (**Fig**. 1h), but allows for evidence inputs that are not aligned with the line attractor and that accumulate over time through non-normal autonomous dynamics^6^. In all three hypotheses, the one-dimensional line attractor/slow manifold is stable, meaning that the autonomous dynamics flow towards it (**Fig**. 1f-h). Because these three hypotheses were each designed to explain a particular set of the phenomena observed in decision-making experiments, a broader range of experimental observations could suggest dynamics that have not been previously hypothesized. As but one example, the autonomous dynamics may contain discrete attractors that do not lie at the endpoints of a one-dimensional slow dynamics manifold; many other arrangements are possible. In the data-driven approach we describe below, *F* is estimated purely from the spiking data and the timing of the sensory input pulses, without incorporating any assumptions from the behavioral DDM or other existing hypotheses.

Dissociating between autonomous and input dynamics requires neural recordings during a decision unfolding over a time period that includes intervals both with and without momentary evidence inputs. We trained rats to perform a task in which they listened to randomly timed auditory pulses played from their left and right and reported the side where more pulses were played^36^ (**Fig**. 1a). The stochastic pulse trains allow us to sample neural responses time-locked to pulses, which are useful for inferring the input-driven dynamics, and also the neural activity in the intervals between pulses, which is useful for inferring the autonomous dynamics. Expert rats are highly sensitive to small differences in the auditory pulse number (**Fig**. 1b; Extended Data Fig. 2a), and the behavioral strategy of rats in this task is typically well captured by gradual accumulation of evidence, which is at the core of the DDM^36–38^.

While the rats performed this task, we recorded six frontal cortical and striatal regions with chronically implanted Neuropixels probes (**Fig**. 1j-k; Extended Data Fig. 2b). The frontal orienting fields (FOF) and anterior dorsal striatum (dStr) are known to be causally necessary for this task and are interconnected^39–41^. The dorsomedial frontal cortex (dmFC) is a major anatomical input to dStr^42^, as confirmed by our retrograde tracing (Extended Data Fig. 2c) and is also causally necessary for the task (Extended Data Fig. 1d). The dmFC is interconnected with the medial prefrontal cortex (mPFC), and less densely, the FOF, the primary motor cortex (M1)^43^, and the anterior ventral striatum (vStr)^42^.

## Unsupervised discovery reveals transitions in dynamical regime and neural mode

To test the current attractor hypotheses and to allow discovery of dynamics that were not previously hypothesized, a flexible yet interpretable method is needed. We used a novel deep learning method (Flow-field Inference from Neural Data using deep Recurrent networks; FINDR^31^) that infers the low-dimensional stochastic dynamics that best account for population spiking data. The low-dimensionality of the description is critical for interpretability. Prominent alternative deep learning-based tools for inferring neural latent dynamics involve models in which these latent dynamics have hundreds of dimensions and are deterministic^29,44,45^. In contrast, FINDR infers latent dynamics that are low-dimensional and stochastic. The stochasticity in the latent dynamics accounts for noise in the decision process that contributes to errors. FINDR uses artificial neural networks as function approximators to infer the function *F* (Eq. 1; **Fig**. 1c)^46,47^. Specifically, the decision-related dynamics *F* are approximated with gated neural stochastic differential equations^48^ (**Fig**. 2a). Each of the many neurons’ firing rates at each timepoint is modeled as a weighted sum of the ***z*** variables, followed by a softplus nonlinearity, which can be thought of as approximating neuronal current-frequency curves^10^ (**Fig**. 2b). The weighting for each neuron (vector ***w***_*n*_ for neuron *n*, comprising the *n*-th row of a weight matrix ***W***; **Fig**. 2b) is fit to the data, and to aid the interpretability of ***z***, we transform ***W*** post-training such that its columns are orthonormal and it therefore acts as a rotation. As a result, angles and distances in ***z*** are preserved in ***Wz*** (neural space before softplus nonlinearity). Prior to applying FINDR, we separately account for the decision-independent, deterministic but time-varying baseline firing rate for each neuron (“baseline” in **Fig**. 2b), so that FINDR can focus directly on the choice formation process.

**Figure 2.**
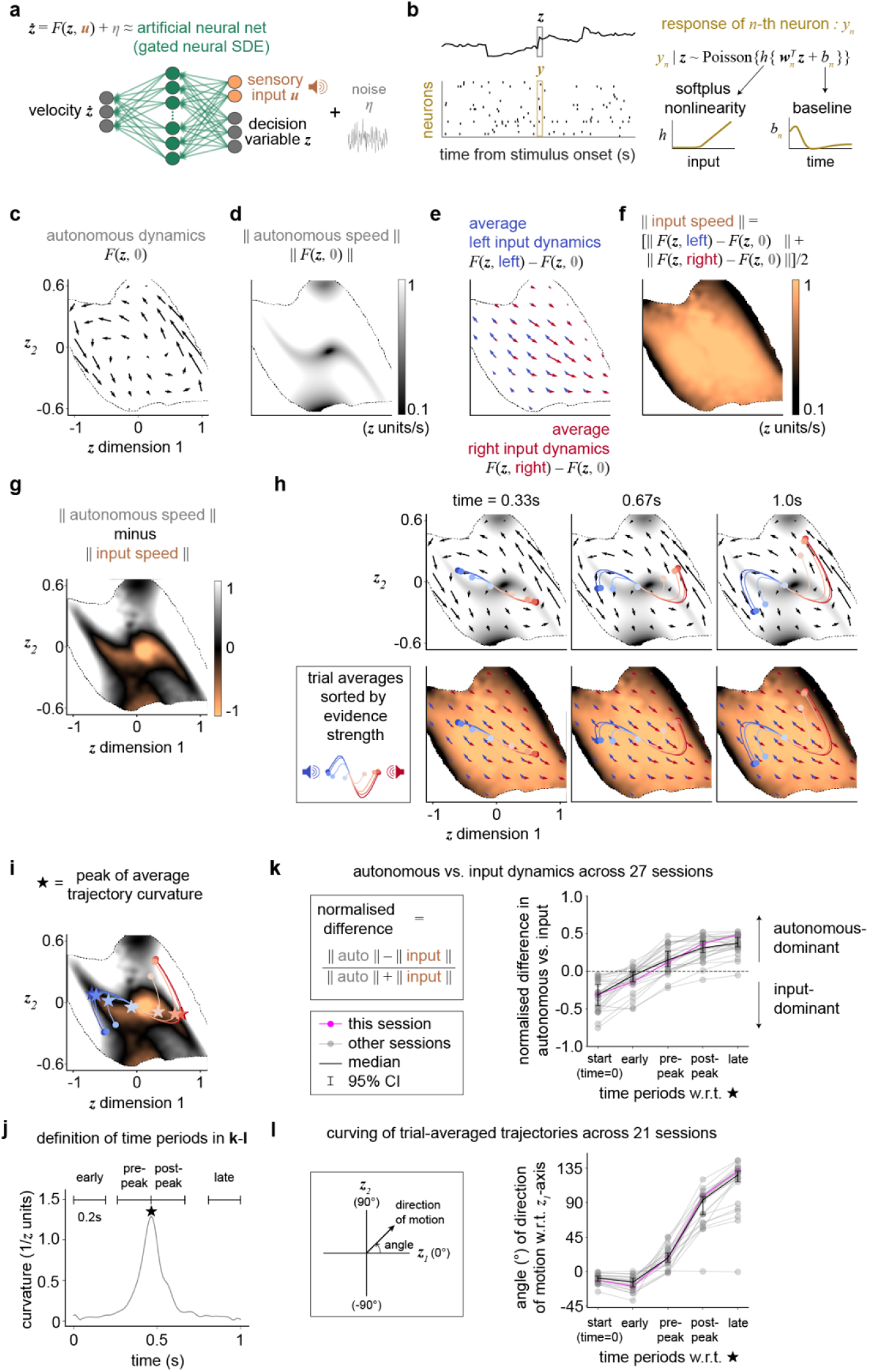
Unsupervised discovery reveals a transition in both the dynamical regime and neural representation that underlie the shift from evidence accumulation to decision commitment. **a**, The decision-related dynamics are approximated with an artificial neural network (gated neural stochastic differential equations; gnSDE) and inferred using the method “Flow-field Inference from Neural Data using deep Recurrent networks” (FINDR)^31^. The deterministic component *F* is approximated using a gated feedforward network, and stochasticity ***η*** is modeled as a Gaussian with diagonal covariance (not constrained to be isotropic). **b**, The parameters of the gnSDE are fit so that the value of the decision variable ***z*** can best capture the spike trains of simultaneously recorded neurons. Conditioned on the decision variable at each time step, the spiking response of each neuron at that time is modeled as a Poisson random variable. A softplus nonlinearity is used to approximate the threshold-linear frequency-current curve observed in cortical neurons during awake behavior. A baseline temporal function is learned for each neuron to account for the decision-irrelevant component of the neuron’s response. **c-h**, Vector field inferred from a representative recording session from 96 choice-selective neurons in dmFC and mPFC. At each time point, we show only the well-sampled subregion of the state space, which is the portion occupied by at least 50 of 5000 simulated single-trial latent trajectories of 1s in duration. **c**, Autonomous dynamics. **d**, Speed of the autonomous dynamics. **e**, Input dynamics of either a left or right click. Unlike sensory stimuli that are continuous over time, such as the random dot motion patches^49^, the stimulus ***u*** in our task is pulsatile. If ***u*=**[1;0] indicates the input to be a single left click, *F*(***z***, [1;0]) − *F*(***z*, 0**) gives the input dynamics *given* that the input was a single left click. However, the average left input dynamics depend on the frequency of left clicks, given by *p*(***u*=**[1;0]|***z***). Therefore, we compute the average left input dynamics *F*(***z***, left) − *F*(***z*, 0**) as *p*(***u*=**[1;0]|***z***)[*F*(***z***, [1;0]) − *F*(***z*, 0**)]. We similarly compute the single right click input dynamics *F*(***z***, [0;1]) − *F*(***z*, 0**), with ***u*=**[0;1] indicating that input was a single right click, and the average right input dynamics *F*(***z***, right) − *F*(***z*, 0**) as *p*(***u*=**[0;1]|***z***)[*F*(***z***, [0;1]) − *F*(***z*, 0**)]. **f**, Speed of the input dynamics. **g**, The strength of the input-driven and autonomous dynamics differ across the decision state space: the input-driven dynamics are far stronger near the origin, whereas the autonomous dynamics are far stronger in the periphery. These differences indicate the decision state-space to be approximately partitioned into two separate regimes: one in which the inputs drive the accumulation of evidence and another in which the autonomous dynamics mediate the commitment to a choice. Difference in the speed between autonomous and input dynamics. **h**, Initially, the trajectories representing the evolutions of the decision variable are strongly driven by inputs and develop along an axis parallel to the direction of the input dynamics, which we term the “evidence accumulation axis.” At a later time, the trajectories become largely insensitive to the inputs, and are instead driven by the autonomous dynamics to evolve along an axis defined by the direction of the autonomous dynamics, which we term the “decision commitment axis.” The sample zone is the same across time points. **i**, To quantify how the difference in the speed between autonomous and input dynamics changes over the course of a trial, we identify the time point at which the latent trajectories curve (indicated as stars), and compute the difference in **g** before and after this time point. **j**, To define when the trajectories curve, we compute the curvature of the trial-averaged trajectories and define the “peak” to be the time point when the curvature is maximum. We then define 200ms time periods with respect to the peak (“pre-peak” and “post-peak”), and with respect to the start and end of the trial (“early” and “late”). **k**, Difference in the speed between autonomous and input dynamics for five different time periods (“start (time=0s)” “early”, “pre-peak”, “post-peak”, and “late”), computed across vector fields inferred from recording sessions that had more than 30 neurons and 400 trials, and sessions where the animal performed with greater than 80% accuracy (*n*=27). The difference is normalized to lie between −1 and 1. **l**, For sessions where the FINDR model with two-dimensional decision variable ***z*** fit significantly better than the FINDR model with one-dimensional ***z*** (*n*=21 out of 27; Extended Data Fig. 5), the direction of motion of the trial-averaged trajectories is computed, and its angle with respect to the ***z***_*1*_-axis for different time periods is shown.

We first confirmed that FINDR can indeed capture population dynamics on a trial-by-trial basis. FINDR provided a good fit to individual neurons’ heterogeneous single-trial firing rates, as well as to the complex dynamics in their peristimulus time histograms (PSTH) conditioned on the sign of the evidence (Extended Data Fig. 4). We found that two latent dimensions suffice to well capture our data (Extended Data Fig. 5). For models with latent dimensions greater than two, the latent dynamics are still mostly confined to two dimensions, and this two-dimensional manifold is approximately an attractor (Extended Data Fig. 5d-g). Focusing on the low-dimensional velocity vector fields (i.e., flow fields) with which FINDR describes the multi-neuron population dynamics, we confirmed that in synthetic data, FINDR-inferred vector fields can be used to distinguish between dynamical systems hypotheses of perceptual decision-making (Extended Data Fig. 3). After these validations, we turned to examining the vector fields estimated by FINDR from the recorded spiking data.

**Fig**. 2c-h illustrates a representative recording session from dmFC and mPFC. We found that generally, the two-dimensional input-driven and autonomous dynamics inferred by FINDR were not well described by the existing hypotheses: in all three hypotheses illustrated in **Fig**. 1d-h, there is a one-dimensional stable manifold that either is, or approximates, a line attractor. In contrast, even though over the first 330 milliseconds the average trajectories evolve along an approximately straight line (**Fig**. 2h), the line is not a one-dimensional attractor, and individual trials will diverge from it. Furthermore, in all three hypotheses of **Fig**. 1d-h, and indeed all other hypotheses we are aware of, the autonomous dynamics play an important role throughout the entire decision-making process. For example, the autonomous dynamics are what enforce the stability of the one-dimensional slow manifolds in **Fig**. 1d-h. In contrast, at least within the space of the latent variable ***z***, the FINDR-inferred dynamics suggest that initially, motion in neural space is dominated and driven by the inputs to decision-making regions (i.e., by the input-dependent dynamics), not the autonomous dynamics, which are slow in both dimensions (**Fig**. 2c-h), not only one. Later in the decision-making process, the balance between autonomous versus input-driven dynamics inverts, and it is the autonomous dynamics that become dominant. **Fig**. 2g plots the difference in magnitude between the autonomous minus input-driven dynamics (indicated with the color scale), on the plane of the latent dimensions ***z***. The initial dominance of the input-driven dynamics can be seen in the zone near the (0,0) origin of the latent dimensions, at the negative end of the color scale. The later dominance of the autonomous dynamics can be seen in the right and left edges of the sampled region, which are reached later in the decision-making process, and which are at the opposite end of the color scale. Moreover, the direction of instantaneous change driven by the inputs (slightly clockwise from the horizontal in **Fig**. 2e) is not aligned with the direction of the strongest autonomous dynamics in the left and right edges of the sampled region (slightly anti-clockwise from vertical in **Fig**. 2c). The curved trial-averaged trajectories of ***z*** emerge from the change in the overall direction of motion due to this non-alignment in the input direction and the autonomous direction later on in the decision-making process. The change from input-dominated to an autonomous-dominated dynamical regime, as well as the sharp turn in the direction of the neural trajectories that we have described for the example session in **Fig**. 2c-h were observed consistently across rats and behavioral sessions (**Fig**. 2i-l). These observations were also robust to multiple different initializations of the deep neural networks in FINDR, the order in which the mini-batches of our datasets were supplied to FINDR during training, and how we split our datasets into training and test sets (Extended Data Fig. 6). They are therefore a consistent finding of the analysis.

To perform a head-to-head comparison with the three hypotheses of **Fig**. 1d-h, we constructed a variant of FINDR in which the gated neural network that parametrizes *F()* was replaced by a parametrization of the dynamics that was constrained to describe the three pre-existing hypotheses (Extended Data Fig. 7). If the data were well-described by one of these three hypotheses, we would expect this variant (which we refer to as cFINDR, for constrained FINDR) to fit the data well, and in particular, to fit out-of-sample data sets better than FINDR, since it has far fewer parameters than FINDR. However, unconstrained FINDR consistently fit the data better than cFINDR, confirming that previous hypotheses do not adequately capture the data. While one of the previous hypotheses (**Fig**. 1h, suggesting non-normal dynamics with a line attractor) can generate curved trial-averaged trajectories apparently similar to those we see in the data (Extended Data Fig. 7g), there is a key difference, which is that in this particular previous hypothesis, the turn from the initial flow direction induced by the inputs happens early, for the autonomous dynamics causing it are strong the moment the latent state departs from the line attractor. However, our data suggests that there is a more prolonged initial phase of flow along the input directions before the turn, with the stronger autonomous dynamics happening much later in the decision-making process. We believe this underlies the much better fits to the data for FINDR than with cFINDR.

Unsupervised inference of dynamics underlying decision-making, based only on spiking activity and sensory evidence inputs, thus suggests that the process unfolds in two separate sequential regimes. In the initial regime, the dynamics are largely determined by the inputs, with autonomous dynamics playing a minor role. The inputs from sensory evidence (right and left clicks) drive the decision variable to evolve along an axis, parallel to the directions of the input dynamics, that we will term the “evidence accumulation axis.” In a second, later regime, these characteristics reverse, the trajectories representing the evolution of the decision variable become largely independent of the inputs, and are instead mostly determined by the autonomous dynamics. We will term the straight line along the direction of the autonomous dynamics in the later regime the “decision commitment axis.” Of note, the evidence accumulation axis and the decision commitment axis are not aligned to each other. During the regime transition, the trajectories in ***z*** veer from evolving along the evidence accumulation axis to developing along the decision commitment axis. In neural space, this will equate to a transition from evolving along one “mode” (i.e., a direction in neural space) corresponding to evidence accumulation, to another mode, that as we explain below, we believe may correspond to decision commitment.

Although derived entirely from unsupervised analysis of neural spiking activity and auditory click times, these two regimes are reminiscent of the two regimes of the behavioral DDM: namely, an initial regime in which momentary sensory inputs drive changes in the state of a scalar decision variable *z*, and a later regime, after *z* reaches a bound, in which the state becomes independent of the sensory inputs (Extended Data Fig. 9a-b). The correspondence between the two regimes inferred from spiking activity and the behavioral DDM suggests that the transition between regimes may correspond to decision commitment. It further suggests that a modified neural implementation of the DDM, focusing on key aspects of the two regimes, could be a simple model that captures many aspects of the neural data, while having far fewer parameters than FINDR and thus greater statistical power. We next develop this model, and show that it can be used to precisely infer the regime transition time on each trial, and test the proposal that this transition corresponds to decision commitment.

## Simplified model of coordinated transitions in dynamical regime and neural mode

The FINDR-inferred vector fields suggest a rapid transition from a strongly input-driven regime to an autonomous-dominant regime that is conceptually similar to the transition from evidence accumulation to decision commitment regimes of the behavioral DDM (**Fig**. 3a-b). The DDM has been shown to describe behavior in a wide range of decision-making tasks, including tasks in which the stimulus duration is determined by the environment^36,40,50,51^ as used here. This raises the possibility that the FINDR-inferred dynamics may be approximated by a simplified model in which the decision variable behaves exactly the same as in the behavioral DDM.

**Figure 3.**
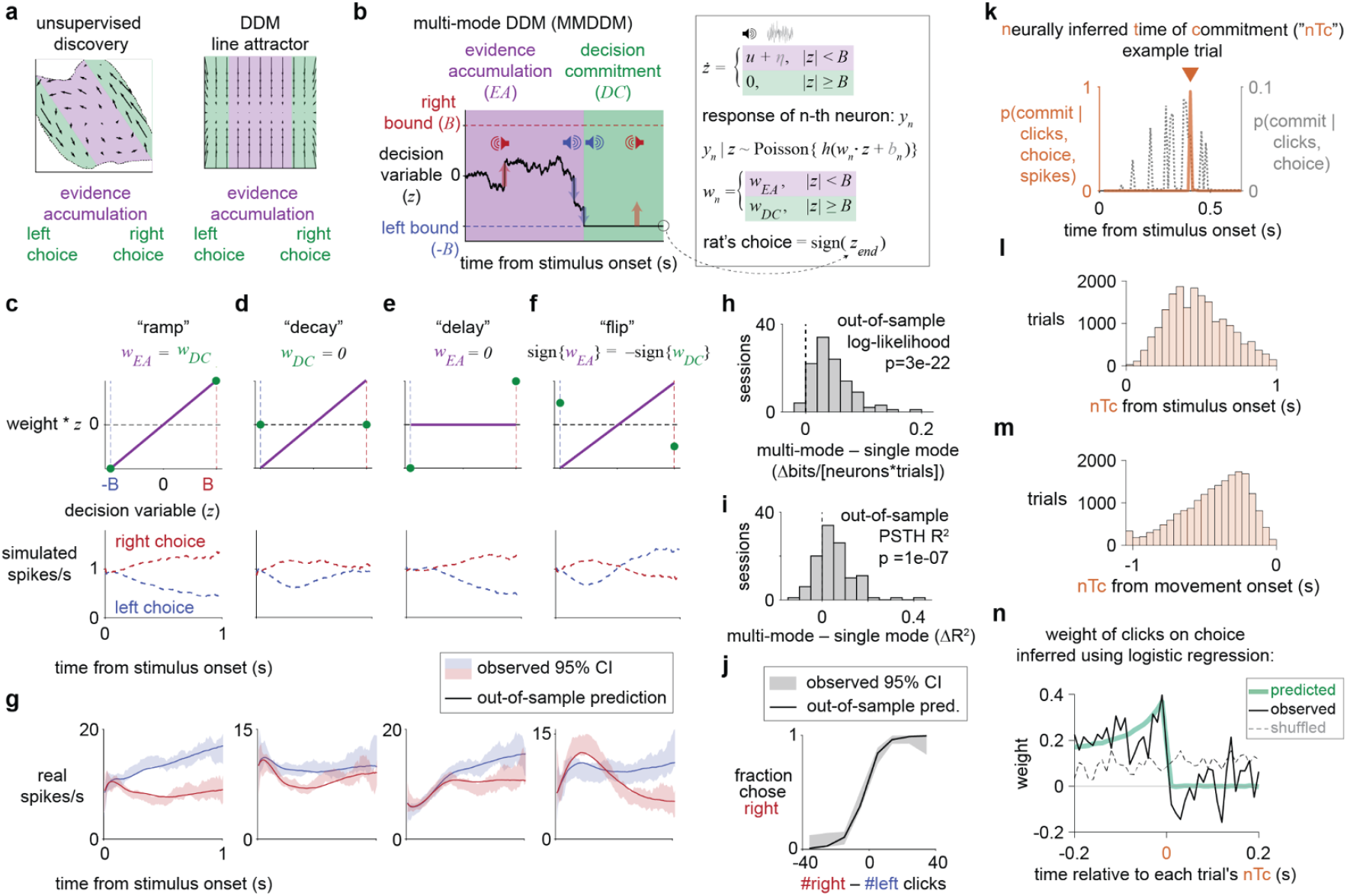
Sensory evidence abruptly ceases to affect the choice after the neurally inferred time of decision commitment (nTc), which is an internal event estimated using a simplified model of the discovered dynamics. **a**, The velocity vector field of both the discovered dynamics and the DDM line attractor can be partitioned into an evidence accumulation regime (EA) and a decision commitment regime (DC). **b**, The multi-mode drift-diffusion model (MMDDM), a simplified model of the discovered dynamics. As in the behavioral DDM, momentary evidence (*u*) and noise (*η*) are accumulated over time in the decision variable (*z*) until *z* reaches either *-B* or *+B*. Abruptly at this moment, the animal commits to a decision: *z* becomes fixed and insensitive to evidence. Here, also at this moment, each neuron’s encoding weight (*w*), mapping *z* to the neuron’s predicted Poisson firing rate *y*, abruptly changes from *w*_*EA*_ to *w*_*DC*_. The mapping from *z* to *y* passes through the softplus nonlinearity *h* and depends on baseline *b*. **c**, Despite its simplicity, the model can capture heterogeneous profiles of individual neurons. The “ramp” temporal profile in a neuron’s PSTH can be generated by setting *w*_*EA*_ and *w*_*DC*_ to be the same. **d**, The “decay” profile is simulated by setting *w*_*DC*_ to zero because as time passes, it is more and more likely to reach the bound and for the encoding to be mediated by *w*_*DC*_ rather than by *w*_*EA*_. When *w*_*DC*_ is zero, there can be no selectivity. **e**, A “delay” profile is generated by setting *w*_*EA*_ to zero because the probability of reaching the bound is zero for some time after stimulus onset. **f**, The “flip” profile is produced by setting *w*_*EA*_ and *w*_*DC*_ to have opposite signs. **g**, For actual neurons, the MMDDM captures the diversity in choice-related temporal profiles. **h**, The MMDDM has a higher out-of-sample likelihood than a 1-dimensional DDM without a neural mode switch. **i**, The MMDDM achieves a higher goodness-of-fit (coefficient-of-determination, R^2^) of the choice-conditioned PSTH’s. **h-i**, P-values were computed using two-sided sign tests. **j**, Behavioral choices are well predicted. **k**, Representative single trial: the inferred time of commitment is far more precise neural activity is included in the inference process (“nTc”) than when inferred solely from sensory stimulus timing and choice behavior. **l**, Among the 34.7% of trials for which the commitment times could be inferred, the nTc occur at highly variable times relative to time from onset of the auditory click trains. The fraction of nTc declines for longer stimulus durations because the stimulus duration on each trial is randomly drawn from 0.2-1.0s. **m**, nTc timing is also highly varied across trials relative to the moment the rat initiates its motor response by leaving the center fixation port. The leftmost bin contains trials on which the nTc occurred more than 1s before movement. **n**, Supporting the interpretation of nTc as decision commitment, and despite the highly variable timing of nTc, sensory evidence presented before nTc impacts the animal’s decision but evidence presented after it does not. Trials for which the estimated time of commitment occurred at least 0.2s before stimulus offset and 0.2s or more after stimulus onset were included for this analysis (9397 of 55,057 trials across 115 sessions/12 rats).

The regime transition coincides with a rapid reorganization in the representation of the decision process in the neural population. To quantify this reorganization, we can consider the activity of each neuron as a dimension in space, and an axis in this neural space as a “neural mode.” Seen in this way, the regime change from evidence accumulation to decision commitment is coordinated with a fast transition in the neural mode. The proposal is conceptually similar to the rapid change in neural modes from motor preparation to motor execution^52^. Therefore, we ask whether a simplified model based on the idea of a rapid, coordinated transition in dynamical regime and neural mode can capture the key features of FINDR-inferred dynamics as well as a broad range of experimental observations.

In what we will call the “multi-mode” or “minimally-modified” drift-diffusion model (MMDDM), a scalar decision variable *z* evolves as in the behavioral DDM, governed by three parameters (**Fig**. 3b; Extended Data Fig. 8a-b; Methods). The key addition in the MMDDM is that neurons encode this decision variable in a different way before and after the decision commitment bound is reached. For each neuron, two scalar weights, *w*_*EA*_ and *w*_*DC*_, specify the strength of its encoding of *z* during the evidence accumulation state (i.e., before reaching the bound, *w*_*EA*_) and the decision commitment state (i.e., after reaching the bound, *w*_*DC*_). When *w*_*EA*_ and *w*_*DC*_ are constrained to be the same, the MMDDM reduces to a DDM with a single neural mode. In the DDM line attractor hypothesis of **Fig. 1**, if the autonomous dynamics towards the line attractor are strong relative to the noise, trajectories will be largely 1-dimensional, and will be well approximated by a single-mode DDM. Because neurons multiplex both decision-related and unrelated signals^53,54^, MMDDM accounts for spike history and, as with FINDR, decision-unrelated baseline changes (Extended Data Fig. 8c-f). All parameters are learned simultaneously by jointly fitting to all spike trains and behavioral choices.

The MMDDM can account for a broader range of neuronal profiles (**Fig**. 3c-g) than the single mode DDM, which captures only the ramp-like neuronal temporal profiles (Extended Data Fig. 9). For the vast majority of recording sessions, the data are better fit by the MMDDM than the single mode DDM (cross-validated; **Fig**. 3h-i), and we confirmed that the choices can be well captured (**Fig**. 3j; Extended Data Fig. 8g) and that the vector fields inferred from real spike trains approximately match the vector fields inferred from the spike trains simulated by the MMDDM (Extended Data Fig. 8h). Additional validations are shown in Extended Data Fig. 8i-n. Moreover, because the end of the stimulus is the same on every trial aligned to the onset of fixation (1.5 s), stimulus offset was not an input in MMDDM, consistent with the lack of abrupt neural changes at stimulus offset (Extended Data Fig. 15).

In the MMDDM, the state transition from evidence accumulation to decision commitment in the MMDDM, and a consequent switch from *w*_*EA*_ to *w*_*DC*_, directly implements a change in neural mode between accumulation and commitment, which has been previously suggested^7,55^. However, it remains unclear whether a moment of a neural mode change in fact corresponds to the animal making up its mind, in part because no method has been previously developed to estimate the moment of commitment on each trial. MMDDM allows precise inference of each trial’s moment of internal commitment to a decision, based on the recorded neural activity (“nTc”; **Fig**. 3k). The resulting nTc estimates varied widely from trial to trial and are not time locked to the start of the stimulus (**Fig**. 3l), end of the stimulus (Extended Data Fig. 10n), or the onset of the motor response (**Fig**. 3m). nTc’s occur much later than the onset of peri-movement kernels inferred from generalized linear models of single-neuron spike trains (Extended Data Fig. 14)^54,56^, indicating that nTc’s are not the onset of the encoding of action plans.

A core prediction of the MMDDM is that if indeed it correctly infers the time of internal decision commitment (“nTc”), then we should be able to observe the contribution of the auditory click inputs to the behavioral choice ceasing abruptly at the estimated decision commitment time. This prediction can be tested by aligning each trial to the nTc, and then measuring the time-varying weight of the stimulus fluctuations on the behavioral choice (referred to as the psychophysical kernel^57–59^). We used a logistic regression model to estimate the weight of each timepoint’s auditory clicks on the subject’s behavioral choice. Remarkably, consistent with the MMDDM prediction, the psychophysical weight of stimulus fluctuations abruptly diminishes to zero after the MMDDM-inferred time of commitment (**Fig**. 3n; Extended Data Fig. 10). Because these commitment times are broadly distributed on different trials (**Fig**. 3l-m), if we instead align trials to the stimulus onset, the abrupt change is no longer observable, and we obtain a smooth psychophysical kernel (Extended Data Fig. 10e-h). Additional characterizations of the nTc are shown in Extended Data Fig. 10i-q. This result provides behavioral confirmation that there exists an internal event during perceptual decision-making after which sensory inputs are ignored and can be inferred from spiking data using the nTc.

## Abrupt and gradual changes at decision commitment

Perceptual decision-making involves a puzzling diversity in the temporal profiles of choice-selective neurons, with some displaying a ramp-to-bound profile, others exhibiting a step-like profile, and some falling in between a ramp and a step^3–5^. This diversity might be explained by a rapid reorganization in population activity at the time of decision commitment. Not all neurons would be equally coupled to this change: a neuron similarly engaged in evidence accumulation and decision commitment would exhibit a ramp-to-bound profile, whereas a neuron more strongly engaged in commitment would show a steep–almost discontinuous–step. Moreover, neurons that are more strongly engaged in accumulation would reflect a ramp-and-decline profile. Can the changes in neuronal responses around the time of decision commitment explain the continuum of ramping and stepping profiles?

We find evidence for these predictions when we grouped neurons by whether they are more, less, or similarly engaged in evidence accumulation relative to decision commitment (Methods; **Fig**. 4a-d; Extended Data Fig. 11a). The peri-commitment neural response time histogram (PCTH) averaged across neurons similarly engaged in accumulation and commitment shows a ramp-to-bound profile. The PCTH resembles a step for the neurons more engaged in commitment, and shows a ramp-and-decline profile for neurons more engaged in accumulation. Even without grouping neurons, we find support for these predictions in the principal component analysis (PCA) on the PCTH’s (Methods; **Fig**. 4e). The first three principal components correspond to the ramp-to-bound, step, and ramp-and-decline profiles. These results show that concurrent gradual and abrupt changes in neural responses around the time of decision commitment explain the continuum of ramp-to-bound and discrete step-like profiles.

Abrupt changes at decision commitment appear inconsistent with a phenomenon that is observed in many studies of decision-making: smoothly curved trial-averaged trajectories in low-dimensional neural state space^6–8^. Similar phenomena are observed in our data: the trial-averaged trajectories for left and right choices do not separate from each other along a straight line, but rather along curved arcs (**Figs**. 2h, 4f). These smoothly curving arcs may result from averaging over trajectories with an abrupt turn aligned to decision commitment, which occurs at different times across trials (**Fig**. 3l-m). Consistent with this account, the smooth curves in low-dimensional neural state space can be well captured by the out-of-sample predictions of MMDDM (**Fig**. 4g; Extended Data Fig. 12), but not a 1-dimensional DDM without a neural mode switch (**Fig**. 4h). These results indicate that the MMDDM, a simplified model of the discovered dynamics, can well capture the widespread observation of smoothly curved trial-averaged trajectories.

**Figure 4.**
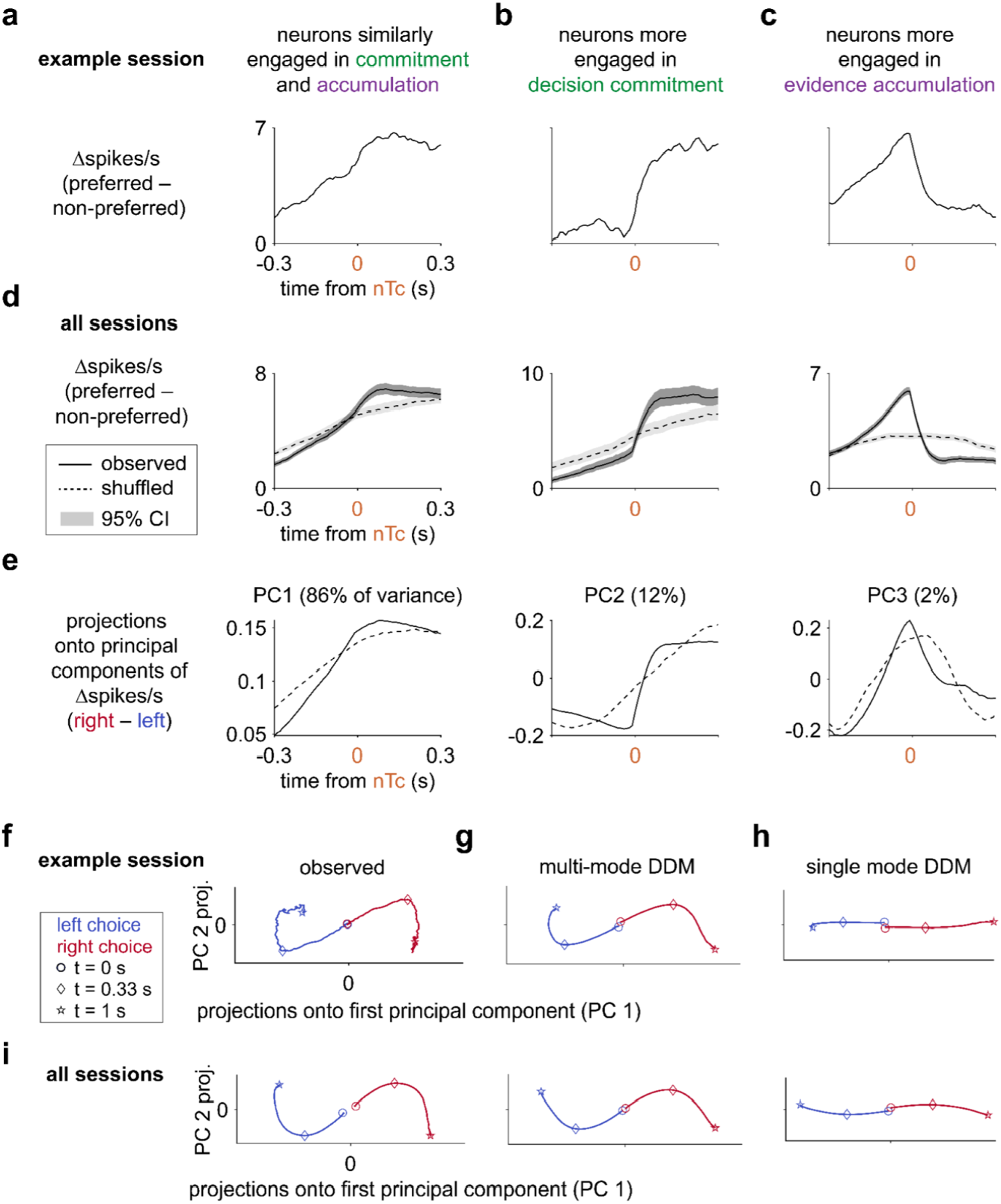
MMDDM captures ramping and stepping profiles and trial-averaged curved trajectories. **a**, The peri-commitment time histogram (PCTH) averaged across neurons that are similarly engaged in decision commitment and evidence accumulation show a ramp-to-bound profile. “Preferred” refers to the behavioral choice for which a neuron emitted more spikes. Example session. N=23 neurons. **b**, In the same session, the PCTH averaged across 10 neurons that are more strongly engaged in commitment has an abrupt, step-like profile. **c**, The ramp-and-decline profile characterizes the PCTH averaged across the 43 neurons from the same session that are more strongly engaged in evidence accumulation. **d**, Across sessions, the ramp-to-bound, step-like, and ramp-and-decline profile characterize the PCTH of neurons similarly engaged in commitment and accumulation (1,116), more engaged in commitment (414), and more engaged in accumulation (1,529), respectively. **e**, The ramp-to-bound, step-like, and ramp-and-decline profiles are observed in the first three principal components of the PCTH’s. **f**, Curved trial-averaged trajectories. **g**, Out-of-sample predictions of the MMDDM. **h**, Out-of-sample predictions of the single mode DDM fail to account for the trial-averaged trajectories. **i**, Results are similar when neurons are pooled across sessions.

## A gradient across brain regions in the strength of neural mode transitions

We observed dynamics with a neural mode transition across the multiple frontal cortical and striatal areas that we recorded: The (cross-validated) MMDDM better captures neural responses in each brain region than the single mode DDM, than an impulsive or a leaky integrator model (**Fig**. 5a; Extended Data Fig. 13a-c). Other signatures of dynamics observed here, including peri-commitment neural changes (Extended Data Fig. 11a) and curved trial-averaged responses (Extended Data Fig. 12a) can be observed within each brain region. These results indicate that the dynamics observed here generalize across frontal cortex and striatum.

**Figure 5.**
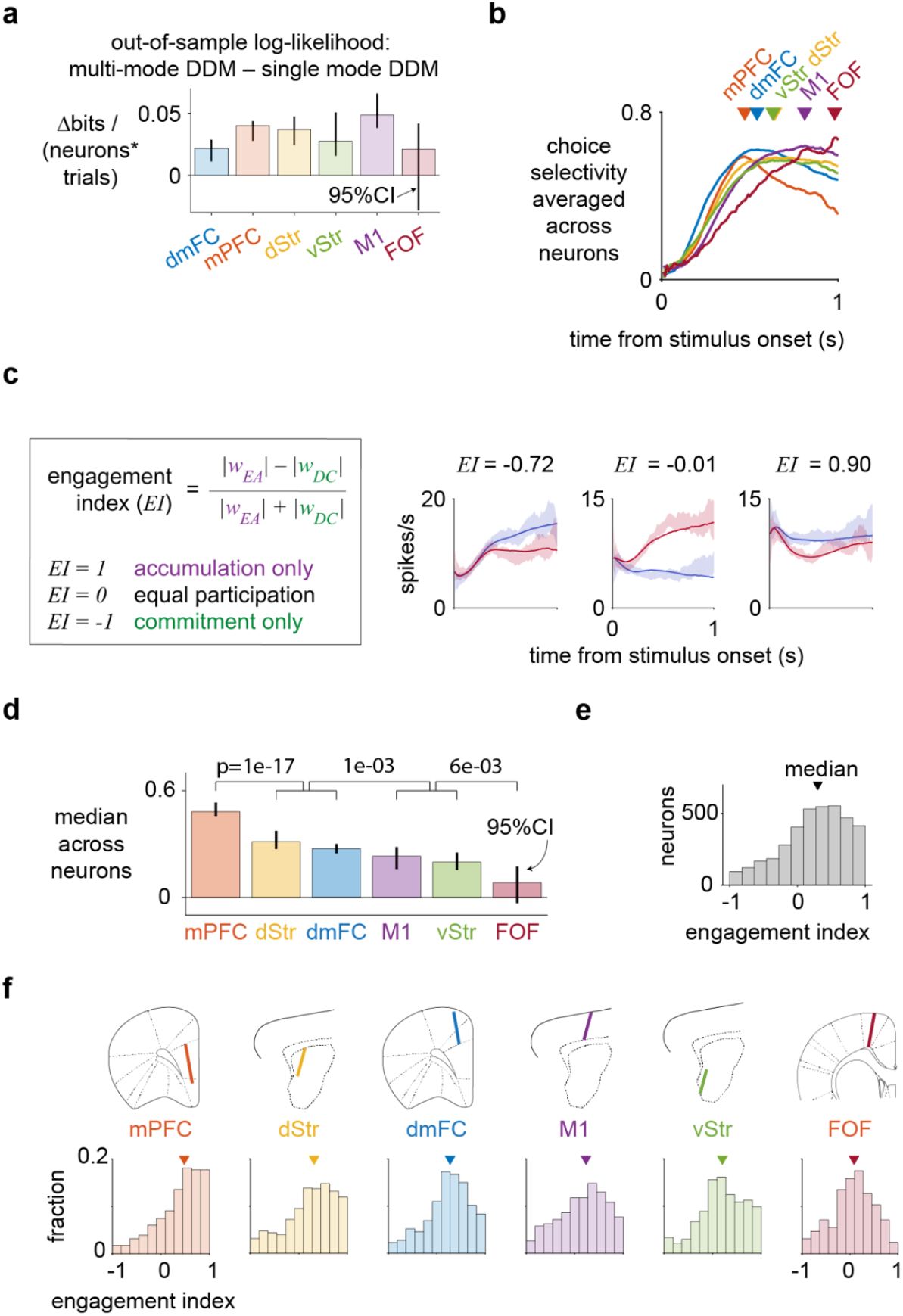
A gradient across brain regions in the strength of neural mode transitions. **a**, MMDDM better captures the data than the single mode DDM. Error bar indicates 95% bootstrapped confidence intervals across sessions. N = 29 dmFC sessions, 29 mPFC, 86 dStr, 74 vStr, 75 M1, and 7 FOF. **b**, The neuron-averaged choice selectivity has different temporal profiles across brain regions: mPFC neurons are most choice-selective near the beginning, while FOF neurons are most choice-selective toward the end. **c**, Using fits from MMDDM, a scalar index quantifies each neuron’s relative level of engagement between evidence accumulation and decision commitment. **d**, A gradient of relative engagement in evidence accumulation and decision commitment across frontal brain regions. The rank order of the median engagement index well matches the rank of the latency to peak choice selectivity. Error bar indicates 95% bootstrapped confidence intervals across neurons. P-values were computed using two-sided sign tests. **e**, The median engagement index is not centered at zero, indicating that frontal cortical and striatal neurons are more strongly engaged in evidence accumulation than in decision commitment. **f**, The differences between the brain regions are apparent in their distribution: whereas the medial prefrontal cortex (mPFC)’s distribution is centered near 1, the distribution for frontal cortical orienting fields (FOF) neurons is centered at zero.

Nonetheless, quantitative differences can be observed across brain regions. The choice selectivity (a measure, ranging from −1 to 1, of the difference in firing rates for right versus left choice trials; Extended Data Fig. 9i) averaged across neurons shows different temporal profiles across brain regions (**Fig**. 5b). Whereas mPFC neurons are most choice-selective near the beginning, FOF neurons are most choice-selective toward the end. Focusing on the latency to the peak of the temporal profiles, the latency is the shortest for mPFC and the longest for FOF. This difference in the latency to peak across brain regions can be explained in the context of separate neural modes for evidence accumulation and decision commitment: neurons that are more strongly engaged in evidence accumulation (*w*_*EA*_ > *w*_*DC*_) have a shorter latency to peak selectivity than neurons that are more strongly engaged in decision commitment (*w*_*DC*_ > *w*_*EA*_). The latencies to peak choice selectivity suggest these frontal brain regions may differ in their relative engagement in evidence accumulation and decision commitment.

To test this possibility, we examined the encoding weights in MMDDM. For each neuron, we computed a scalar index that compares its relative engagement in the two processes, by taking the difference between the absolute values of the *w*_*EA*_ and *w*_*DC*_ and normalizing by the sum (**Fig**. 5c). A neuron with an index near 1 is engaged only in evidence accumulation (and shows a decay), whereas a neuron with an index near −1 is engaged only in decision commitment (and exhibits a delay). A neuron with a ramp profile must have an index near zero, which corresponds to equal levels of engagement. The distribution of the indices was unimodal, indicating the engagement in accumulation and commitment is distributed rather than clustered among neurons (**Fig**. 5e-f). The engagement indices show that brain regions differ in their relative engagement in evidence accumulation: mPFC, dmFC, and dStr are more strongly engaged in evidence accumulation than the regions vStr, M1, and FOF. The rank order of the brain regions by the engagement index is similar to the rank order by time to peak latency. This result indicates that differences in choice-related encoding across frontal cortical and striatal regions can be understood in terms of relative participation in evidence accumulation versus decision commitment.

## Discussion

How attractor dynamics govern the formation of a perceptual choice has been long debated^1,6,9^. Here we suggest that, for decisions on the timescale of hundreds of milliseconds to seconds, an initial input-driven regime mediates evidence accumulation, and a subsequent autonomous-dominant regime subserves decision commitment. This regime transition is coupled to a rapid change in the representation of the decision process by the neural population: the initial neural mode (i.e., direction in neural space) representing evidence accumulation is largely orthogonal to the subsequent mode representing decision commitment. In this sense is reminiscent of other covert cognitive operations, such as attentional selection, that also involve a change in neural mode^60^.

If this coupled transition in dynamical regime and neural mode indeed corresponds to the time of decision commitment, sensory evidence presented after the transition would have minimal impact on the subject’s decision, for the subject would have already committed to a particular choice. Behavioral analysis confirmed this prediction in the experimental data (**Fig**. 3n), leading us to conclude that the transition is indeed a signal for covert decision commitment. We refer to each trial’s estimate of the presence and timing of such a transition, which is based on the sensory stimulus and firing rates of simultaneously recorded neurons, as “nTc”.

How do decisions end? In reaction time paradigms of perceptual decision-making, subjects are trained to respond as soon as they make a decision. The moment the animal initiates its response is then used to operationally define when it commits to a choice^61,62^. In these paradigms decision commitment is overt, as it is closely linked to the onset of the movement subjects make to report their choice^62^. Here, in contrast, using an experimenter-controlled duration paradigm, we found a decision commitment signal (nTc) that is covert in the sense of occurring at a time highly variable with respect to the timing of the external motor action used to report the decision, which it can precede by as much a second or more (**Fig**. 3m). It is also highly variable with respect to stimulus onset (**Fig**. 3l) or offset (Extended Data Fig. 10n). It is thus an internal signal, largely defined by coordination across neurons, not by its timing with respect to external events. The peri-commitment neural responses observed here contrast sharply with the ramp-and-burst neural responses observed in animals trained to couple their decision commitment with response initiation^62^ in a reaction time task.

While the timing of the nTc signal reported here makes it very distinct from motor execution, the signal is also distinct from action preparation or planning. The beginning of action planning carries no implication as to whether sensory evidence presented subsequent to that will or will not be ignored. Indeed, in perceptual decision-making tasks, preliminary action preparation, driven by choice biases induced by previous trials, is often observed to begin even before the sensory stimulus, as reported previously^54,56,63^ and found in our own data (Extended Data Fig. 14). In contrast, commitment to a decision implies that evidence presented subsequent to the commitment will no longer affect the subject’s choice. Here we found that nTc corresponds to such a decision commitment moment. This was the case both at the neural level, where it correlates with a substantial decrease in the effect of sensory inputs on neural responses in the regions we recorded (**Fig**. 2); and at the whole-organism behavioral level, in the sense that sensory evidence before nTc affects the subject’s choices, but sensory evidence after nTc does not (**Fig**. 3n).

While the behavioral DDM is a widely used model of decision-making, other frameworks are also prevalent, such as the linear ballistic accumulator^64^ or urgency gating^65^. It is notable that the dynamics inferred by FINDR, obtained in a data-driven, unsupervised manner from spike times and auditory click times alone, resulted in regimes that match the characteristics of the behavioral DDM, but not of the alternatives. This match led us to explore a simplified model, the MMDDM, in which a scalar latent decision variable evolves as in the DDM, but is represented in different neural modes before versus after decision-commitment. The neural mode change indicates that a downstream decoder of the categorical choice can improve its accuracy by selectively reading out from neurons whose post-commitment weights are large in magnitude. A possible mechanism for the neural mode change is an input from ascending midbrain neurons, which is suggested by a recent finding in a working memory task that midbrain neurons, in response to an external auditory cue, trigger rapid reorganization of motor cortex activity to switch from planning-related activity to a motor command that initiates movement in mice^66^.

We found the MMDDM to provide a parsimonious explanation of a variety of experimental findings from multiple species: across primates and rodents, sensory inputs and choice are represented in separate neural dimensions^6–8,56,67,68^ across time, and neither sensory responses nor the neural dimensions for optimal decoding of the choice are fixed^7,56,68^. These phenomena, along with other observations including diversity in single neuron dynamics^53,54^, curved average trajectories^7,8,67^, choice behavior^36^, and some vigorously debated phenomena such as a variety of single-neuron ramping versus stepping temporal profiles^4,5,69^, are all captured by the MMDDM. However, we do not see MMDDM as a unique or a unified model of perceptual decision-making. Rather, we see it as a simple yet useful approximation, a “minimally-modified” DDM, and a stepping stone toward a unified model of decision-making.

Single trial trajectories, taken together, filled out the two-dimensional latent space inferred by FINDR. But when averaged over trials of a given evidence strength (**Fig**. 2h), they evolved along a one-dimensional curved trajectory. Looking exclusively along this 1-d manifold, the dynamics resemble those of the bistable attractor hypothesis (**Fig**. 1f)^1^ in the sense of a 1-d unstable point at the origin, with autonomous dynamics growing stronger the further the system is from the origin. However, the bistable attractor hypothesis, as well as the other two hypotheses in **Fig**. 1g,h posit a one-dimensional manifold of slow autonomous dynamics, along which evidence accumulation evolves, and towards which other states are attracted^1,35^. In contrast, the FINDR-inferred dynamics --which are inferred from single trials, not averaged trials,-- suggest an initial *two*-dimensional manifold of slow autonomous dynamics, with an origin that is not a saddle point, but is instead weakly but strictly unstable (Extended Data Fig. 16a,e). Sensory evidence inputs drive evidence accumulation along one of these slow dimensions. The other slow dimension corresponds to the decision commitment axis, along which autonomous dynamics will become dominant later in the process. During the initial evidence accumulation, why would there be slow autonomous dynamics along this second dimension? It is well known that during perceptual decision-making, non-sensory factors such as which choice option was rewarded on the previous trial^70–72^ have a significant effect on subjects’ choices. But the neural mechanisms for how such other factors and sensory evidence interact to produce the subject’s final choice remain unclear. We speculate that the slow autonomous dynamics along the decision commitment axis provide a mechanism for inputs driven by these other factors to influence choice, in a manner that is orthogonal and independent of the evolving representation of the accumulating evidence. A recent study^73^ described neural trajectories that were well-described by non-normal dynamics^74,75^. Consistent with this, the two-dimensional FINDR-inferred autonomous dynamics around the origin are also non-normal (Extended Data Fig. 16b,c), although with a key difference with respect to refs. ^73–75^, which is that here the origin is unstable (Extended Data Fig. 16a,e).

One recently proposed method to infer autonomous dynamics, applied to data from a task that did not require accumulating evidence over time, proposed that variety across individual neurons’ tuning curves could lead to curved 1-d decision manifolds^21^. However, their method cannot yet infer input dynamics and thus cannot yet analyze data from tasks with evidence that arrives gradually over time; such an extension would have to be realized before we can assess whether the curvature their approach could infer would correspond to the curvature we described here for accumulation of evidence. Importantly, inferring input dynamics in addition to the autonomous dynamics was critical to our observation that a change in dynamical regime, from input-dominated to autonomous-dominated, appeared to coincide with the change in neural mode (**Fig**. 2). This observation was key for our hypothesis that this event (“nTc”) could correspond to decision commitment, for the development of the MMDDM simplified model to estimate nTc, and for the experimental confirmation that nTc is indeed the moment when sensory evidence ceases impacting the subject’s decision (**Fig**. 3n).

Finally, our approach expands the classic repertoire of techniques used to study perceptual decision-making. We inferred the decision dynamics directly from neural data rather than assuming a specific hypothesis, and we took steps to enhance the human-interpretability of the discovered dynamics: the unsupervised method (FINDR) focuses on low-rather than high-dimensional decision dynamics, and the mapping from latent to neural space (before each neurons’ activation function) preserves angles and distances. Based on key features of the inferred latent dynamics, we developed a highly simplified, tractable model (MMDDM) that is directly relatable to the well-known drift diffusion model framework. We found that the MMDDM, despite its simplicity, was able to describe a broad variety of previously-observed phenomena, and allowed us to infer the subject’s internal decision commitment times on each trial. Pairing deep learning-based unsupervised discovery with simplified, parsimonious models may be a promising approach for studying not only perceptual decision-making but also other complex phenomena.

## Extended Data Figures

**Extended Data Figure 1.**
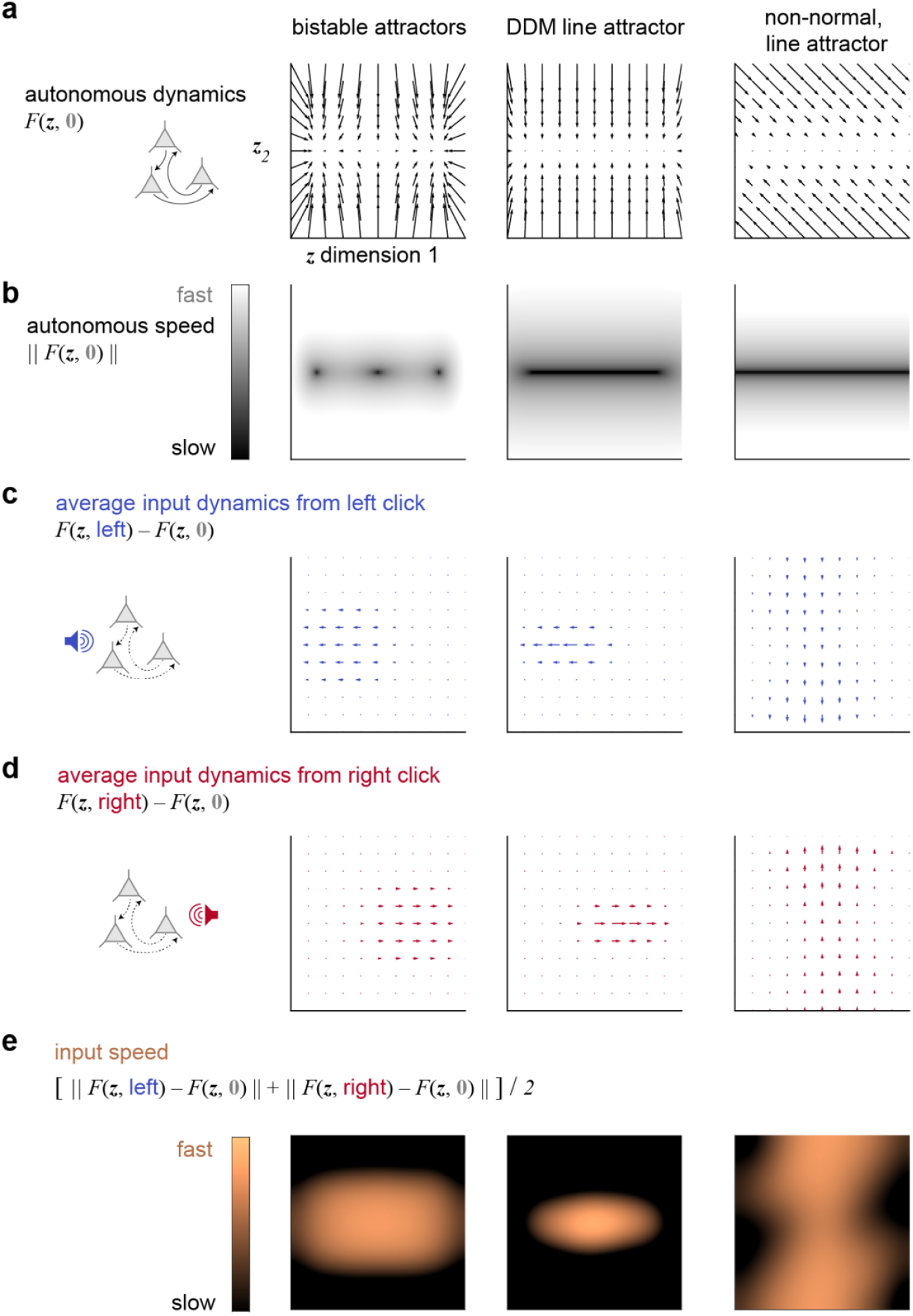
Attractor hypotheses of perceptual decision-making. In these hypotheses, the decision process is represented by the state of a dynamical system, which we refer to as the “decision variable (***z***)” and is depicted as two-dimensional here but may have fewer or more dimensions. An attractor is a set of states for which the dynamical system tends to move toward, from a variety of starting states. When ***z*** is in an attractor state, small perturbations away from the attractor tend to return the system toward the attractor. An attractor can implement the commitment to a choice and the maintenance of the choice in working memory. **a**, In all these hypotheses, the attractors are implemented by the autonomous dynamics, which corresponds to the deterministic dynamics *F* in the absence of inputs and depends only on ***z*** itself. In the bistable attractors hypothesis, there are two discrete attractors, each of which corresponds to a choice alternative. In the DDM line attractor hypothesis, the autonomous dynamics form not only two discrete attractors but also a line attractor in between. The intervening line attractor allows an analog memory of the accumulated evidence when noise is relatively small. In the line attractor hypothesis with non-normal dynamics, the autonomous dynamics form a line attractor, and a separate readout mechanism is necessary for the commitment to a discrete choice. **b**, The autonomous speed is the magnitude of the autonomous dynamics. A dark region corresponds to a steady state, which can be an attractor, repeller, or saddle point. In the bistable attractors hypothesis, the left and right steady states are each centered on an attractor, and the middle is a saddle point. In both the DDM line attractor hypothesis and the hypothesis that has non-normal dynamics with a line attractor, the steady states correspond to attractors. **c-d**, Input dynamics corresponding to a left and right auditory pulse, respectively. Here we show the “effective” input dynamics, which is multiplied by the frequency *p*(***u***|***z***) to account for the pulsatile nature and the statistics of the stimuli in our task (in contrast to **Fig**. 1e, in which the input dynamics were presented without the multiplication of the frequency, which is appropriate for stimuli that are continuous over time, such as the random dot motion patches^49^). Whereas in the bistable attractor and DDM line attractor, the inputs are aligned to the attractors, in the hypothesis that has non-normal dynamics with a line attractor, the inputs are not aligned. **e**, The input speed is the average of the magnitude of the average left input dynamics and the magnitude of the average right input dynamics.

**Extended Data Figure 2.**
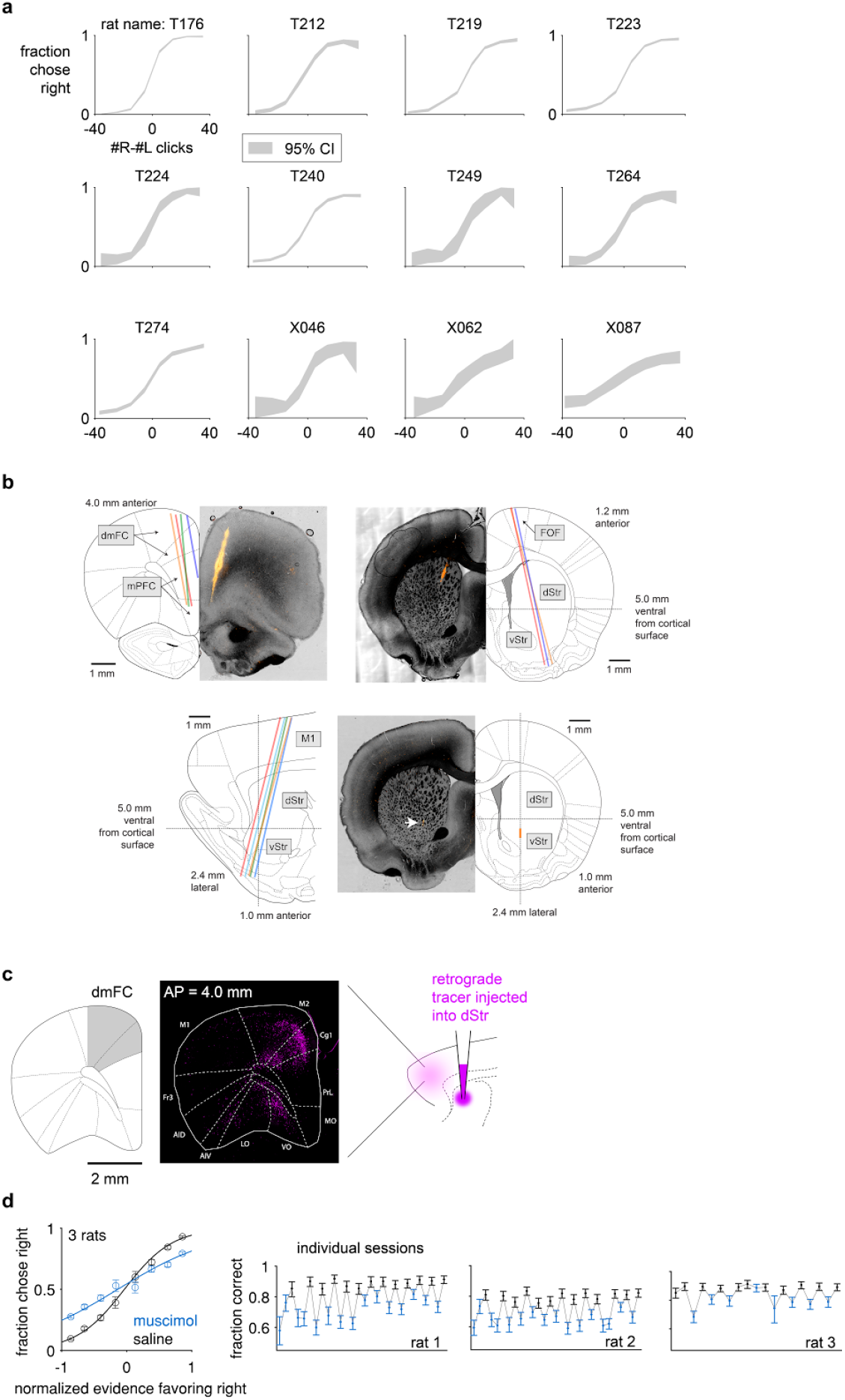
Behavioral performance, histological slices, anatomical tracing, and the causal necessity of dmFC. **a**, Psychometric functions of each of the twelve rats recorded aggregated across recording sessions. **b**, Histological images of probe tracks. Each color indicates a probe chronically implanted in a rat. **c**, Dorsomedial frontal cortex (dmFC) provides a major input to the anterior dorsal striatum (dStr). **d**, dmFC is causally necessary for the auditory decision-making task studied here.

**Extended Data Figure 3.**
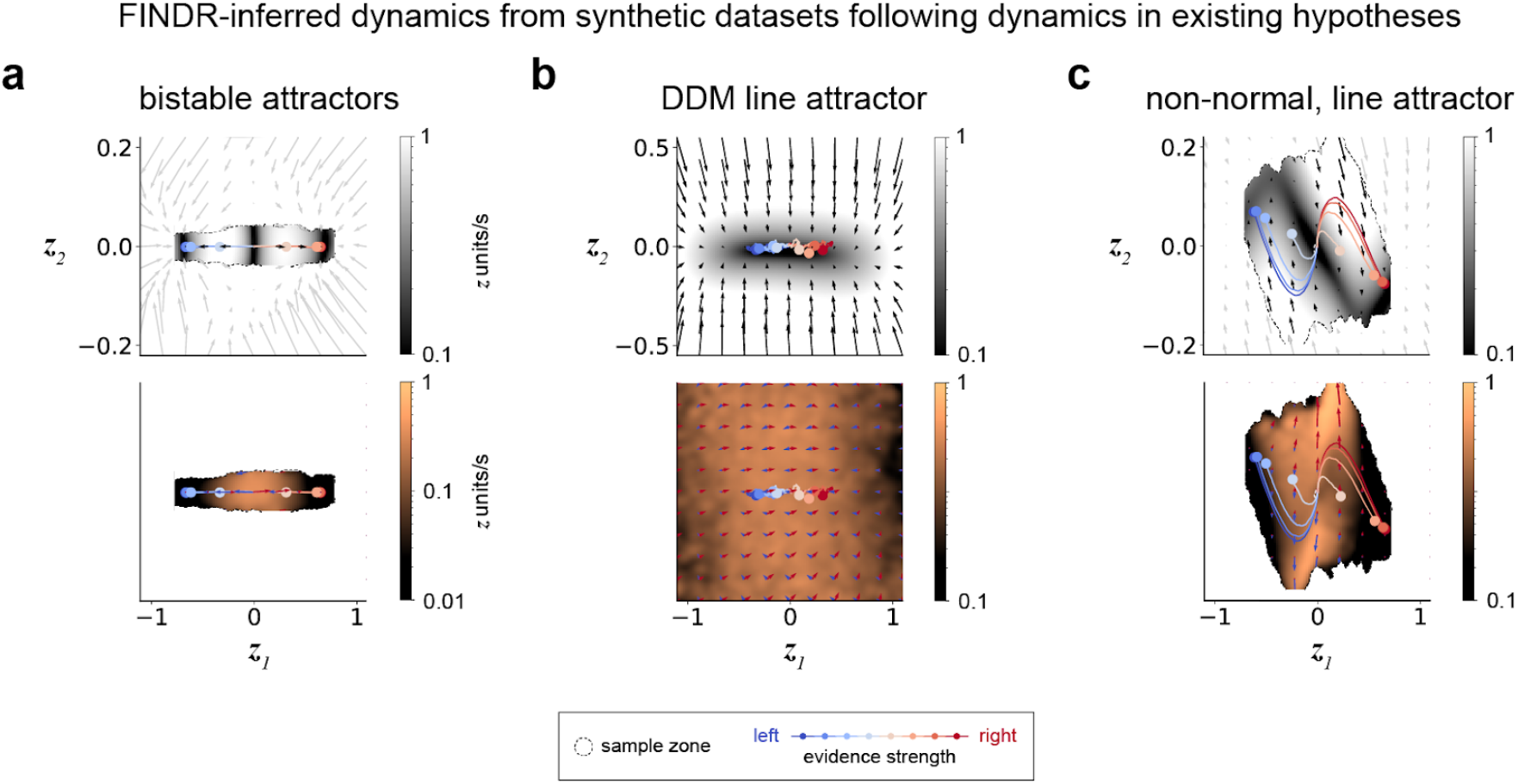
FINDR can be used to distinguish between the dynamical systems hypotheses of perceptual decision-making. **a**, We simulated spikes that follow the bistable attractor dynamics in Extended Data Fig. 1 to create a synthetic dataset with the number of trials, number of neurons, and firing rates that are typical of the values observed in our datasets. Then, we fit FINDR to this synthetic dataset from random initial parameters. The autonomous and input dynamics inferred by FINDR qualitatively match the bistable attractors hypothesis. **b-c**, FINDR-inferred dynamics qualitatively match the dynamics in **Fig**. 1f-h and Extended Data Fig. 1. In panel **b**, the sample zone covers the entirety of the plotted area.

**Extended Data Figure 4.**
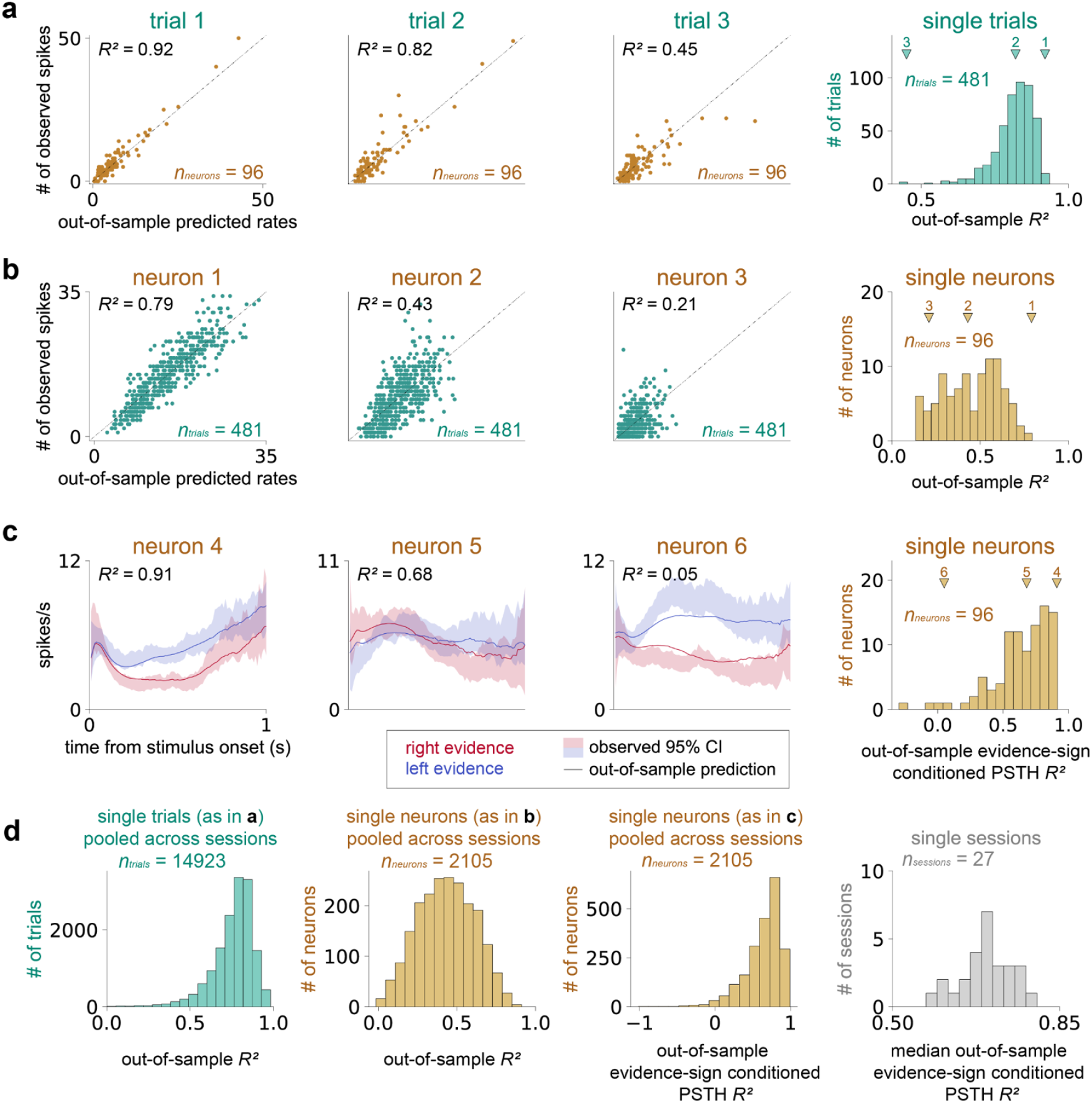
FINDR can well capture the neural responses. **a**-**b**, FINDR captures the underlying firing rates of the single-trial responses of individual neurons from the representative session in **Fig**. 2. **c**, FINDR captures the complex trial-averaged dynamics of individual neurons from the representative session in **Fig**. 2 as can be seen in the peristimulus time histograms (PSTH). The goodness-of-fit is measured using the coefficient of determination (*R*^2^). **d**, FINDR captures the single-trial and trial-averaged responses of individual neurons pooled across 27 sessions. For the histogram showing single trials pooled across sessions, 34 trials that had *R*^2^ < 0 are not shown. Results in **a**-**d** are 5-fold cross-validated.

**Extended Data Figure 5.**
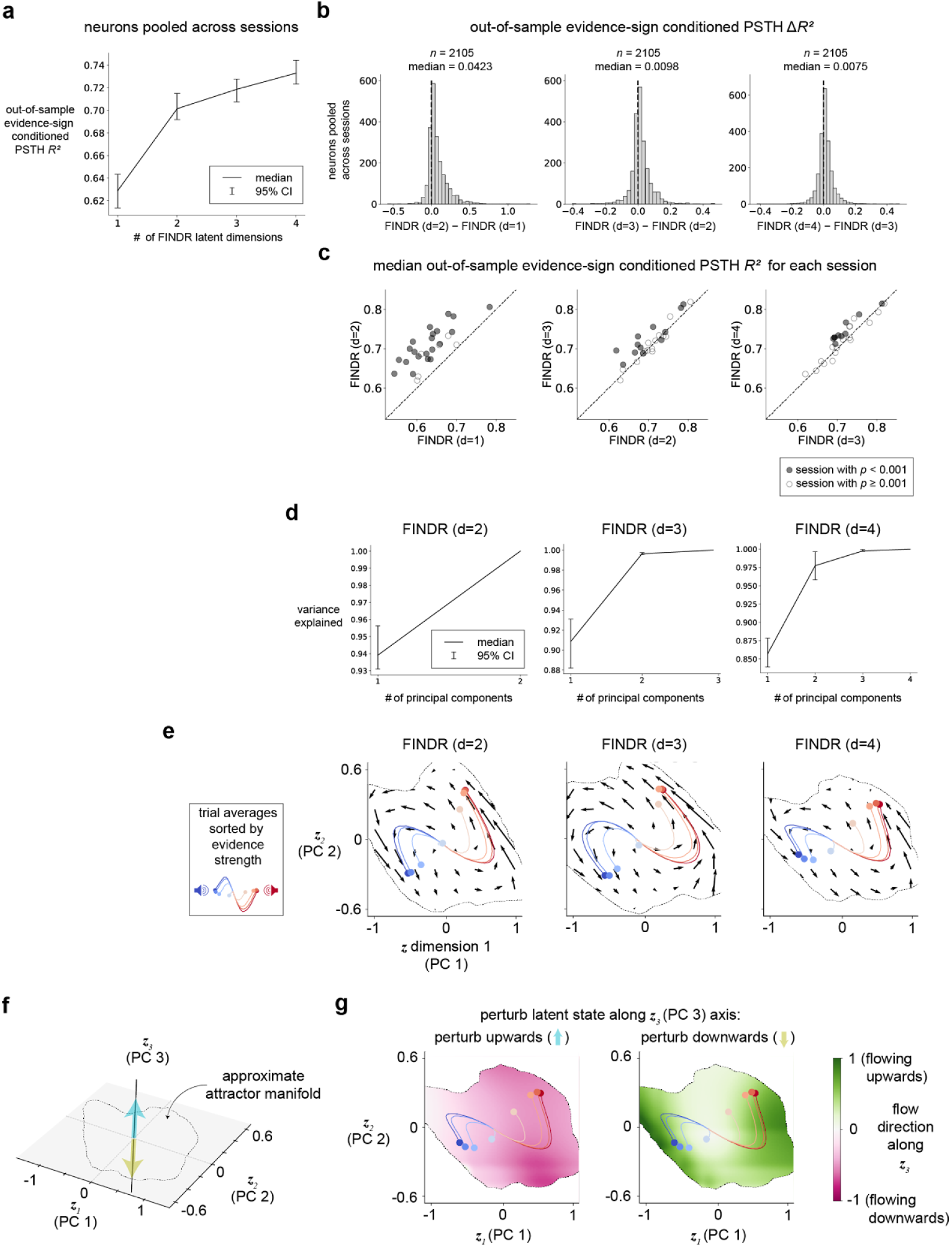
FINDR reveals 2-dimensional decision-making dynamics. **a**, Across different FINDR models with latent dimensions (d) ranging from 1 to 4, we computed the median of the coefficient of determination (R^2^) of the evidence-sign conditioned peri-stimulus time histogram (PSTH) of neurons pooled across sessions. **b**, The median difference in the R^2^ between d=2 and d=1 is significantly different from zero (p<0.001; Wilcoxon signed-rank test). Although the median differences are also significant for the comparison between d=3 and d=2 and the comparison between d=4 and d=3, the magnitude of the difference is relatively small (0.0098 and 0.0075, respectively) compared to the median difference between d=2 and d=1 (0.0423). **c**, We repeated the analysis in **b** without pooling neurons across sessions. Instead, for each session, we computed the median PSTH R^2^ across neurons recorded within that session. Each circle corresponds to a session, and a filled circle indicates a significant difference in the PSTH R^2^ between FINDR models of different dimensionalities (p<0.001; Wilcoxon signed-rank test). **d**, For FINDR models with either 3 or 4 latent dimensions, more than 97% of the variance is captured by the first two principal components (PC’s). PCA was done separately for each session, and the error bars indicate the 95% confidence interval of the median across sessions. **e**, For models with 2 or more dimensions, the vector fields and trajectories projected onto the first two dimensions are qualitatively similar. The vector fields and trajectories were shown for the representative session in **Fig**. 2. The dashed lines demarcate the well-sampled subregion of the state space (i.e., the sample zone). **f**, We evaluated FINDR models with latent dimensions higher than two to see whether the two-dimensional manifold relevant to decision-making dynamics is an approximate attractor manifold. The variance explained by the third PC in the FINDR model with three-dimensional latent dynamics was less than 0.5% (as shown in **d**-**e**), so we turned to the FINDR model with four-dimensional dynamics. In this model, the variance explained by the third PC was around 1.3%. We perturbed the latent states on the manifold along the PC 3 direction. **g**, When the latent states are perturbed (but not too far that the latent states go outside the range along the PC 3 axis covered by the sample zone; see Methods for details), the latent states flow toward the manifold.

**Extended Data Figure 6.**
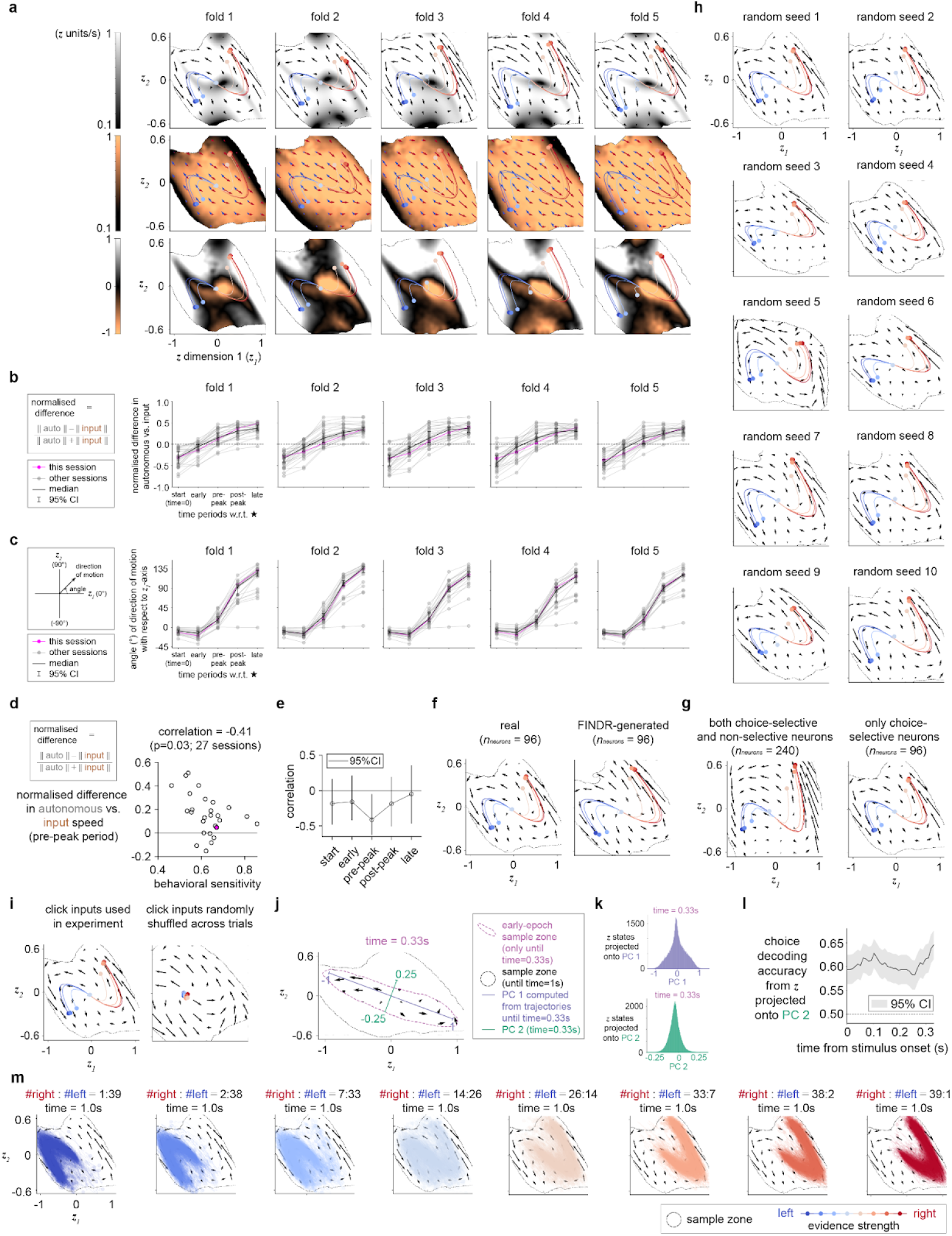
Consistency in FINDR-inferred dynamics. **a**, FINDR-inferred input and autonomous dynamics are consistent across 5 different cross-validation folds as shown for the same session in **Fig**. 2. **b**, Normalized difference in the speed between autonomous and input dynamics for five different time periods (“start (time=0s)” “early”, “pre-peak”, “post-peak”, and “late”) is consistent across folds. **c**, The direction of motion of the trial-averaged trajectories and its angle with respect to the ***z***_*1*_-axis for different time periods is consistent across folds. **d**, Variability in the dynamics across sessions depends in part on the variability in the behavioral performance. For each each session, behavioral sensitivity was estimated as the parameter β in a probit model p(y | x) = Φ(β*x + c), where y is the rat’s left vs. right choice on each trial, x the log-ratio of the right vs. left click rate on that trial, Φ the normal cumulative distribution function, c the constant term in the probit model. The p-value of the Pearson’s correlation was computed using a Student’s t-distribution for a transformation of the correlation. Pink marker indicates the example session. **e**, The linear correlation between the difference in autonomous vs. input dynamics and behavioral sensitivity was negative for all epochs, but reliable only for the pre-peak epoch. The 95% confidence intervals were computed by bootstrapping across sessions. **f**, FINDR reliably recovers the FINDR-inferred dynamics. After fitting FINDR to a dataset, the model parameters were used to simulate a synthetic dataset using the exact same set of sensory stimuli in the real dataset and containing the same number of neurons and trials. From new initial parameter values, FINDR was fit to the simulated data to infer the “FINDR-generated” vector fields. **g**, FINDR is fit to both choice-selective and non-selective neurons. We find similar dynamics to when FINDR is fit to only choice-selective neurons. **h**, We find vector fields that are consistent across multiple different random seeds that change the initialization in the deep neural networks of FINDR and the order in which the mini-batches of the training data are supplied to FINDR during training. **i**, Curved trial-averaged latent trajectories predicted by FINDR depend on the click inputs. When FINDR was fit to data in which the click inputs were randomly shuffled across trials, the trial-averaged latent trajectories remain near the origin. **j**, The dynamics are two-dimensional even in the beginning of the decision period. An early-epoch sample zone indicated by the dotted line was computed using trajectories that were truncated at time=0.33s. The early-epoch sample zone delimits the portion of the state occupied by at least 50 of 5000 simulated single-trial trajectories. **k**, When we compute the PCs for the trajectories truncated at time=0.33s and project the trajectories onto PC 2, the standard deviation along this direction is 20.4% of the standard deviation along PC 1. **l**, We can decode behavioral choice from logistic regression significantly better than chance (dashed line) from the projections of the truncated trajectories onto PC 2. **m**, Single-trial latent trajectories extending to time=1.0s, simulated using stimuli of different evidence strength, which is quantified by the ratio of right and left inputs.

**Extended Data Figure 7.**
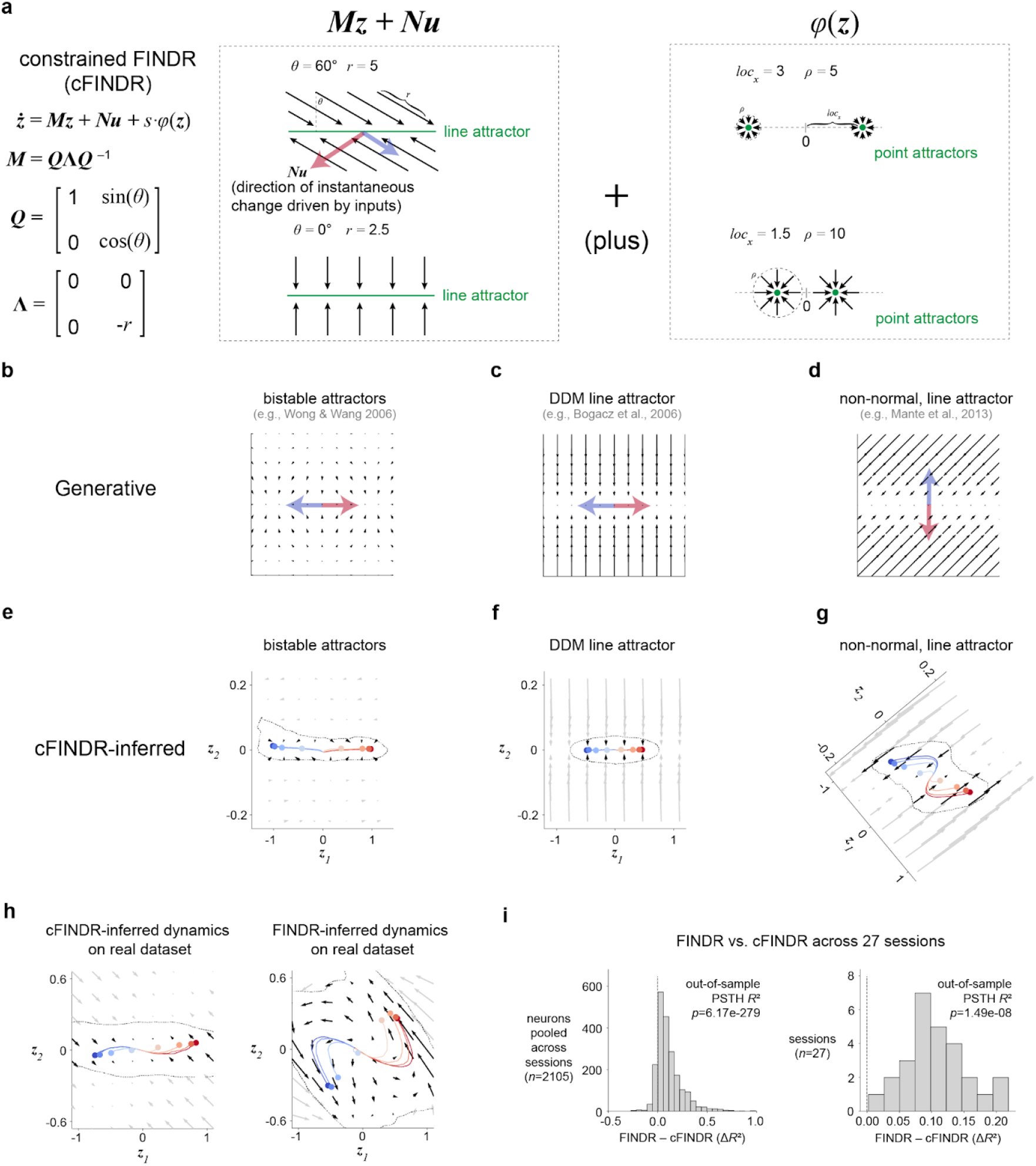
The data are better captured by FINDR than by a variant of FINDR constrained to parametrize the dynamics described by previously proposed hypotheses. **a**, The constrained FINDR (cFINDR) model replaces the neural networks parametrizing *F* in FINDR with a linear combination of affine dynamics, specified by ***M*** and ***N***, and bistable attractor dynamics specified by *φ*. The dynamics are furthermore constrained to be two-dimensional. **b**-**g**, cFINDR model can generate and infer dynamics described by previous hypotheses. **b**, Example bistable attractor dynamics generated from cFINDR. **c**, Example DDM line attractor dynamics generated from cFINDR. **d**, Example non-normal dynamics with a line attractor generated from cFINDR. **e**, cFINDR-inferred dynamics from a synthetic dataset generated using the bistable attractor dynamics in **b. f**, cFINDR-inferred dynamics from a synthetic dataset generated using the DDM line attractor dynamics generated in **c. g**, cFINDR-inferred dynamics from a synthetic generated using the non-normal dynamics with a line attractor in **d. h**, cFINDR-inferred dynamics and FINDR-inferred dynamics on a real representative session. **i**, The coefficient-of-determination (*R*^*2*^) of the evidence-sign conditioned peri-stimulus time histogram (PSTH) computed using fits of FINDR to the data is significantly greater than the *R*^*2*^’s computed using fits of cFINDR (Wilcoxon signed-rank test).

**Extended Data Figure 8.**
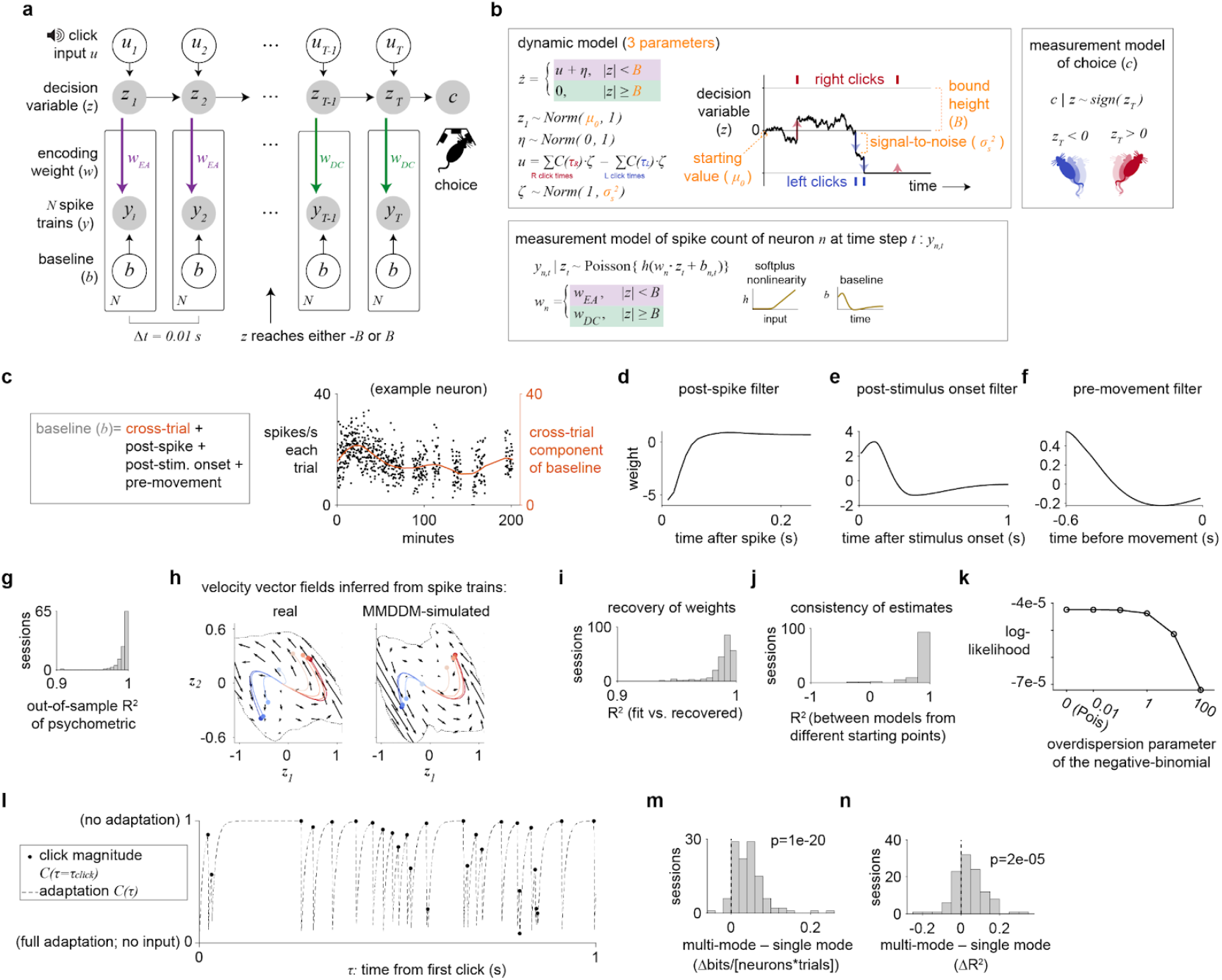
Multi-mode drift-diffusion model (MMDDM). **a**, Directed graph of the MMDDM for a trial with *T* time steps and *N* simultaneously recorded neurons. At each time step, the decision variable *z* depends on external click input (*u*) and its value in the previous time step. The spike train depends on *z* and also a time-varying baseline input. The behavioral choice (*c*) is the sign of the decision variable at the last time step. In this example trial, *z* reaches the bound, and the encoding weight of *z* of each neuron changes from *w*_*EA*_ to *w*_*DC*_. **b**, The MMDDM is an instance of a state-space model, which consists of a dynamic model governing the probability distributions of the latent states (here, scalar decision variable *z*) and measurement models specifying the conditional distributions of the emission (here, spike counts *y* and the rat’s choice *c*) given the value of the latent states. In the dynamic model, *z*’s time derivative (*ż*) is a piecewise linear function. When the absolute value of *z* is less than the bound height *B*, the velocity depends on external click input (*u*) and i.i.d. Gaussian noise (*η*). When *z* reaches either *-B* or *B*, the time derivative is zero. The input of each click emitted at time τ on *z* is scaled by the depressive adaptation from previous clicks, parametrized by *C(*τ), and it is corrupted by i.i.d. multiplicative Gaussian noise ζ with variance 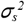. The parameter 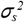 is one of the three parameters learned during fitting and represents the signal-to-noise of the system. The behavioral choice (*c*) is the sign of the decision variable at the last time step. The mapping from *z* to spike train response (*y*) passes through the softplus nonlinearity *h* and depends on baseline *b* and encoding weight *w*. The encoding weight is either *w*_*EA*_ and *w*_*DC*_ depending on *z*. The three parameters that are fit in MMDDM consist of the bound height *B*, the mean *μ*_0_ of starting distribution, and the signal-to-noise of each momentary input. **c**, The baseline input consists of a cross-trial component, parametrized by smooth temporal basis functions, as shown for an example neuron. **d**, The spike history filter of the same neuron. **e**, The post-stimulus filter of the neuron. This filter does not depend on the content of the click train and only depends on the timing of the first click, which is always a simultaneous left and right click. **f**, The kernel of the same neuron to account for movement anticipation. The kernel does not depend on the actual choice of the animal. **g**, The psychometric function is well captured across sessions. **h**, The vector field inferred from real spike trains is confirmed to be similar to that inferred from MMDDM-simulated spike trains for the session “T176_2018_05_03.” **i**, After fitting the model to each recording session, the learned parameters are used to simulate a data set, using the same number of trials and the same auditory click trains. The simulations are used to fit a new model, the recovery model, starting from randomized parameter values. The encoding weights of the accumulated evidence of the recovery model are compared against the weights used for the simulation (which were learned by fitting to the data) using the coefficient-of-determination metric. **j**, Consistency in the encoding weights between the training models during five-fold cross-validation. For each session, a coefficient-of-determination was computed for each pair of training models (10 pairs), and the median is included in the histogram. **k**, Whereas the Poisson distribution requires the mean to be the same as the variance, the negative binomial distribution is a count response model that allows the variance to be larger than the mean μ, with an additional parameter *α*, the overdispersion parameter, that specifies the variance to be equal to μ+*α*μ^2^. When the overdispersion parameter is zero, the distribution is equivalent to a Poisson. Fitting the data to varying values of the overdispersion parameter shows that log-likelihood is maximized with a Poisson distribution for the conditional spike count response. Similarly, when the overdispersion parameter was learned from the data, the best-fit values were all close to zero. **l**, The magnitude of the input after sensory adaptation of each click in a simulated Poisson auditory click train. Based on previous findings^36^, the adaptation strength (φ) is fixed to 0.001, and the post-adaptation recovery rate (k) to 100. The generative click rate is 40 hz, as in the behavioral task. **m**, Sensory adaptation is not critical to the improvement in fit by the MMDDM compared to the single mode DDM. Even without modeling sensory adaptation–by setting φ=1 and k=0, such that every click has the same input magnitude–the out-of-sample log-likelihood is reliably improved by the MMDDM compared to the single mode DDM. **n**, The out-of-sample goodness-of-fit of the PSTH’s is also reliably improved even in the absence of sensory adaptation.

**Extended Data Figure 9.**
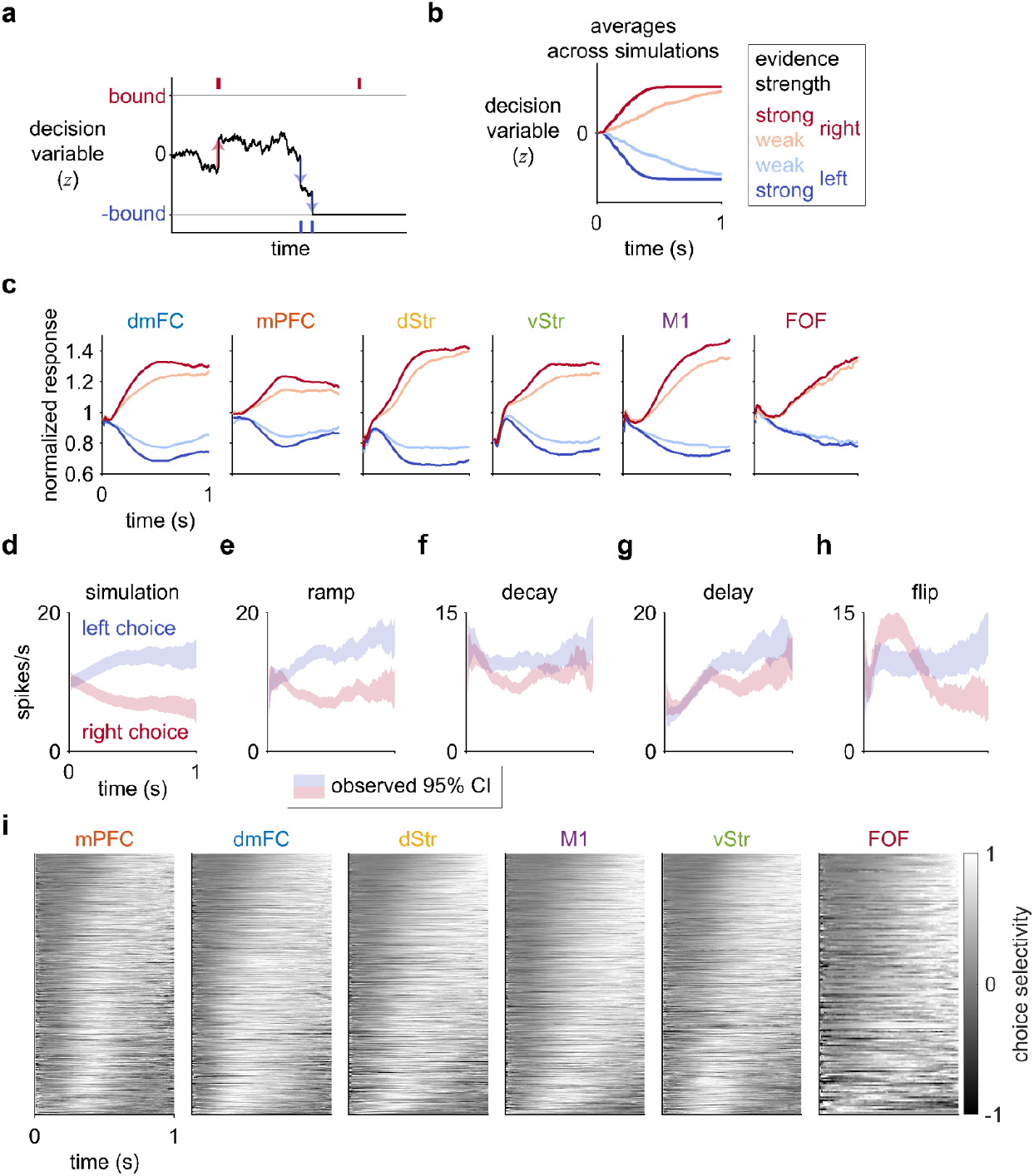
The temporal profiles of the choice selectivity of individual neurons are heterogeneous, and this diversity is not consistent with an one-dimensional neural encoding of the latent variable in the drift-diffusion model (DDM). **a**, In the DDM, noisy inputs are accumulated over time through a scalar latent variable (*z*) until the value of *z* reaches a fixed bound, which triggers the commitment to a choice. **b**, In simulations of the DDM, *z* ramps quickly when the evidence strength is strong and more slowly when the strength is weak. **c**, Responses averaged across both trials and neurons resemble the trajectories of *z* averaged across simulations. Only choice-selective neurons were included. Spikes after the animal began movement (i.e., removed its nose in the center port) were excluded. For this analysis only, error trials were excluded. N = 1324 (dmFC), 1076 (mPFC), 1289 (dStr), 714 (vStr), 822 (M1), 163 (FOF). **d**, The responses of a simulated neuron encoding the DDM with a single neural mode show the ramping dynamics. Shading indicates the bootstrapped 95% confidence interval of the trial-mean of the filtered response. **e**, A neuron with a ramp profile. **f**, A neuron recorded from the session with choice selectivity that decays over time. **g**, A neuron exhibiting a substantial delay in its choice selectivity. **h**, A neuron whose choice selectivity flips in sign. **i**, The diversity of the temporal profile of the choice selectivity of individual neurons is not consistent with a one-dimensional encoding of the DDM.

**Extended Data Figure 10.**
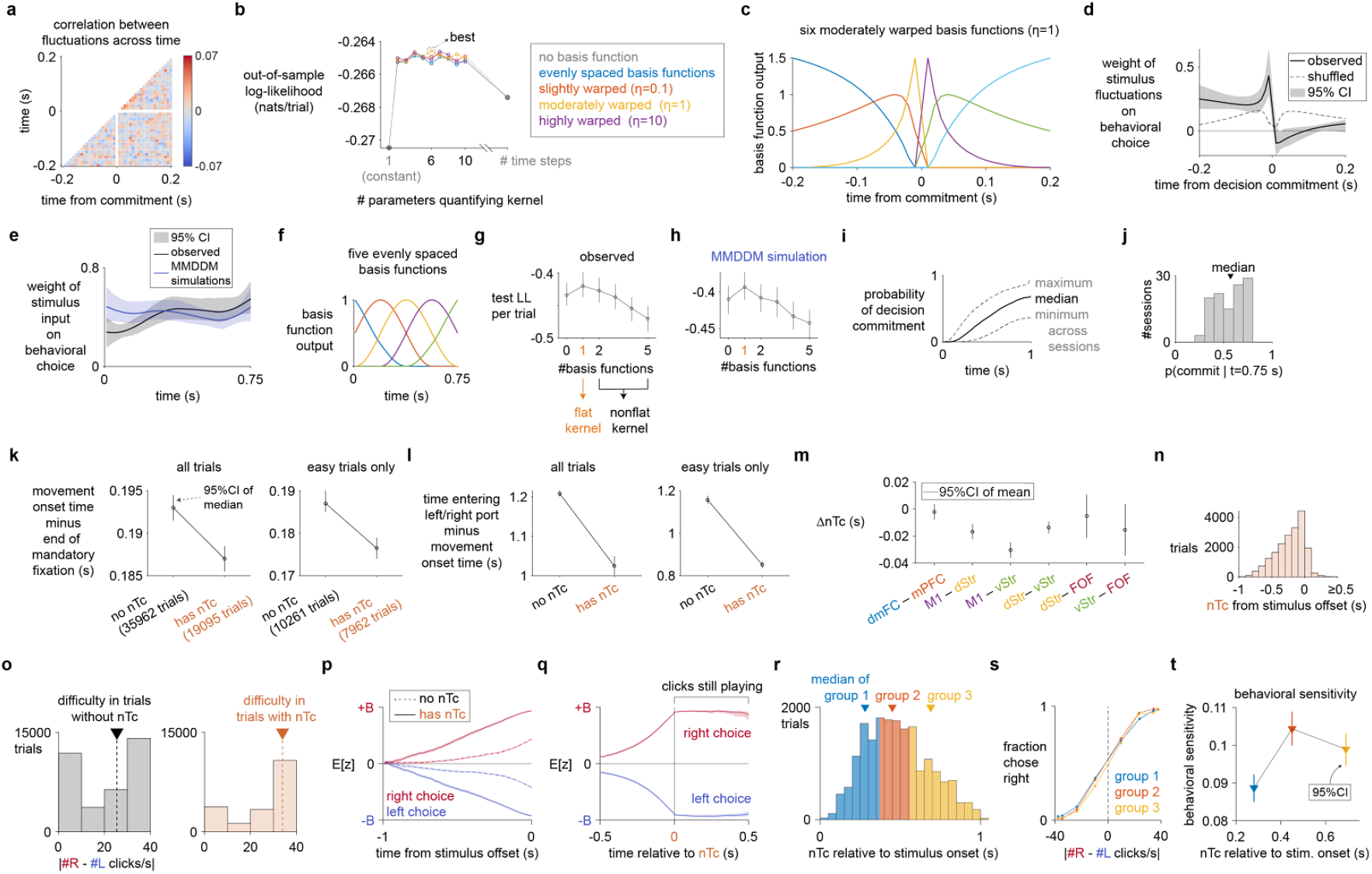
nTc and psychophysical kernels. The psychophysical kernel shown in **Fig**. 3m of the main text was estimated using time bins of 0.01 s, spanning +/−0.2 s around the decision commitment time. In this extended data figure we repeat the psychophysical kernel estimate, but using basis functions and cross-validation to select the basis functions that provide the optimal smoothing of the signal. The results shown here and shown in **Fig**. 3m of the main text are qualitatively similar and lead to the same conclusions. **a**, For the inferred weights of the stimulus fluctuations to be interpretable, the click input fluctuations must not be strongly correlated across time steps. On each time step on each trial, the fluctuation in auditory click input was computed by counting the observed difference in right and left clicks at that time step, and then subtracting from it the expected difference given the random processes used to generate the stimulus. The input fluctuations at time step of t=0 s were excluded because they are strongly correlated with the input fluctuations before decision commitment and strongly anti-correlated with input fluctuations after commitment. **b**, To determine the time resolution of the kernel that best captures the weight of the input fluctuations, 10-fold cross-validation was performed to compare kernels quantified by different numbers of parameters and types of basis functions. The kernel with the lowest temporal resolution is a constant, represented by a single parameter, implying that fluctuations across time have the same weight. At the highest time resolution, the kernel can be parametrized by a separate weight for each time step. At intermediate time resolution, the kernel is parametrized by basis functions that span the temporal window. The basis functions can be evenly spaced across the temporal window, or stretched such that time near t=0 s is represented with higher resolution and time far from t=0 s with lower resolution. The most likely model had six moderately stretched (η=1) basis functions. **c**, The optimal model’s set of six moderately stretched (η=1) basis functions. **d**, Even when using basis functions, the psychophysical kernel is consistent with the core prediction of MMDDM: The psychophysical weight of the stimulus fluctuations on the behavioral choice ceases after the time of decision commitment. Note that no basis function was used in the analysis in **Fig**. 3n. **e**, In contrast to the commitment-aligned kernel, the kernel aligned to the onset of the auditory click trains is smooth. Mean stimulus onset-aligned psychophysical kernel across sessions, estimated using a model with five temporal basis functions. For each session, 10-fold cross-validation was performed on fitting the kernel model to the data, and ten estimated kernels were averaged. Then, the kernels were averaged across sessions. **f**, The onset-aligned psychophysical kernel is parametrized by five evenly spaced radial basis functions. **g**, Cross-validated model comparison shows that a temporally flat psychophysical kernel is most likely given the observed data. **h**, Similarly, given the simulated choices generated by the MMDDM, the out-of-sample log-likelihood is maximized by assuming a flat kernel. **i**, The approximately flat psychophysical kernel inferred from MMDDM-simulated choices is consistent with the MMDDM’s prediction of the probability of decision commitment given the stimulus: throughout the trial, the probability of decision commitment is relatively low, and at no point in the trial is decision commitment an absolute certainty. **j**, At t=0.75 s, the window used to compute the psychophysical kernel, the median probability of decision commitment across sessions is 0.57. **k**, A small but statistically significant effect of whether decision commitment was reached on the “movement onset time,” i.e., the time when the rat withdraws its nose from the fixation port minus the earliest time when the rat is allowed to do so. The effect is not simply due to trial difficulty because it remains when we consider only easy trials (right : left click rate either greater than 38:1 or less than 1:38). **l**, Similar effect of whether commitment was reached on the rat’s “movement execution time,” i.e., the time when the rat reaches either the left or right port minus the time when it withdrew its nose from the fixation port. **m**, Relative timing of decision commitments between pairs of simultaneously recorded brain regions. For each pair of regions, the comparison was made on only the trials on which the threshold for commitment was crossed for both regions. **n**, Inferred times of commitment, relative to stimulus offset. **o**, As expected from the model, nTc’s occur more often in easier trials, i.e., trials with larger generative (experimentally controlled) difference between the left and right click rate. **p**, As expected from the model, the mean value of the latent variable (the expectation under the posterior probability given the spikes and choice) reaches values of larger magnitude on trials on which nTc could be inferred compared to trials on which an nTc could not be inferred. Shading indicates 95% bootstrapped confidence intervals across sessions. **q**, Consistent with the model, even when considering only the period before the clicks were still playing, the mean of the latent variable abruptly plateaus after the nTc. **r**, The trials on which nTc could be estimated were separated into three groups using the terciles of the distribution of nTc relative to stimulus onset. **s**, Psychometric function of each group, showing the fraction of a right choice against the generative (i.e.. experimentally specified) difference between the right and left click rates. **t**, Behavioral sensitivity is higher for trials with longer nTc. A logistic model with two terms (bias and slope) was fit to regress the choice on each trial against the generative difference in click rate. The 95% confidence intervals were computed by bootstrapping across trials.

**Extended Data Figure 11.**
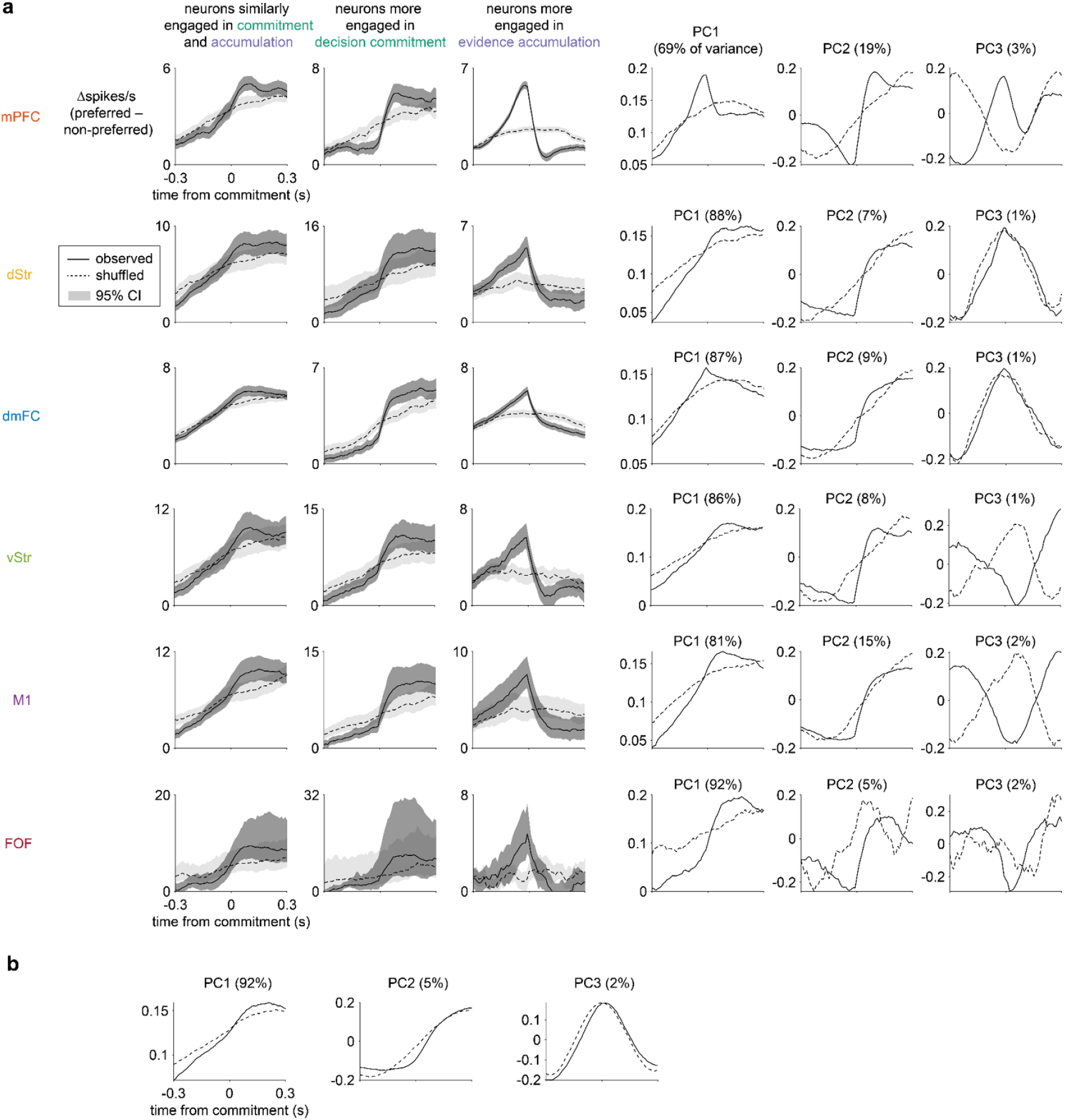
The ramp-to-bound, step-like, and ramp-and-decline profiles can be observed in individual brain regions (panel **a**), and the ramp-to-bound, step-like profiles can be observed in neurons not choice-selective enough to be included in model analysis (panel **b**).

**Extended Data Figure 12.**
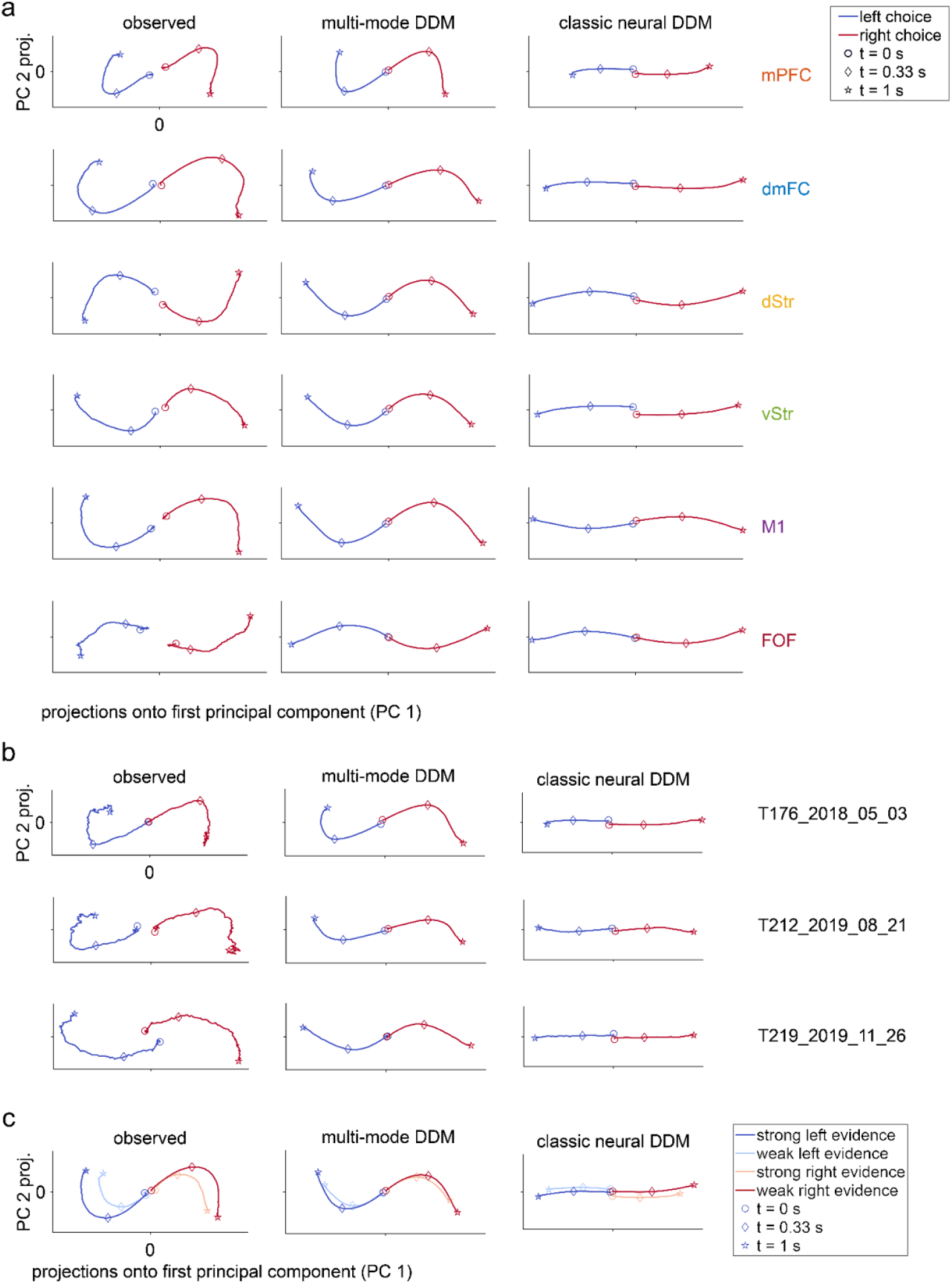
The multi-mode drift-diffusion model (MMDDM) captures curved population trajectories when the analysis is performed on data from **a**, individual brain regions; **b**, individual sessions; and **c**, on trial conditions that depend on not only the choice but also the evidence.

**Extended Data Figure 13.**
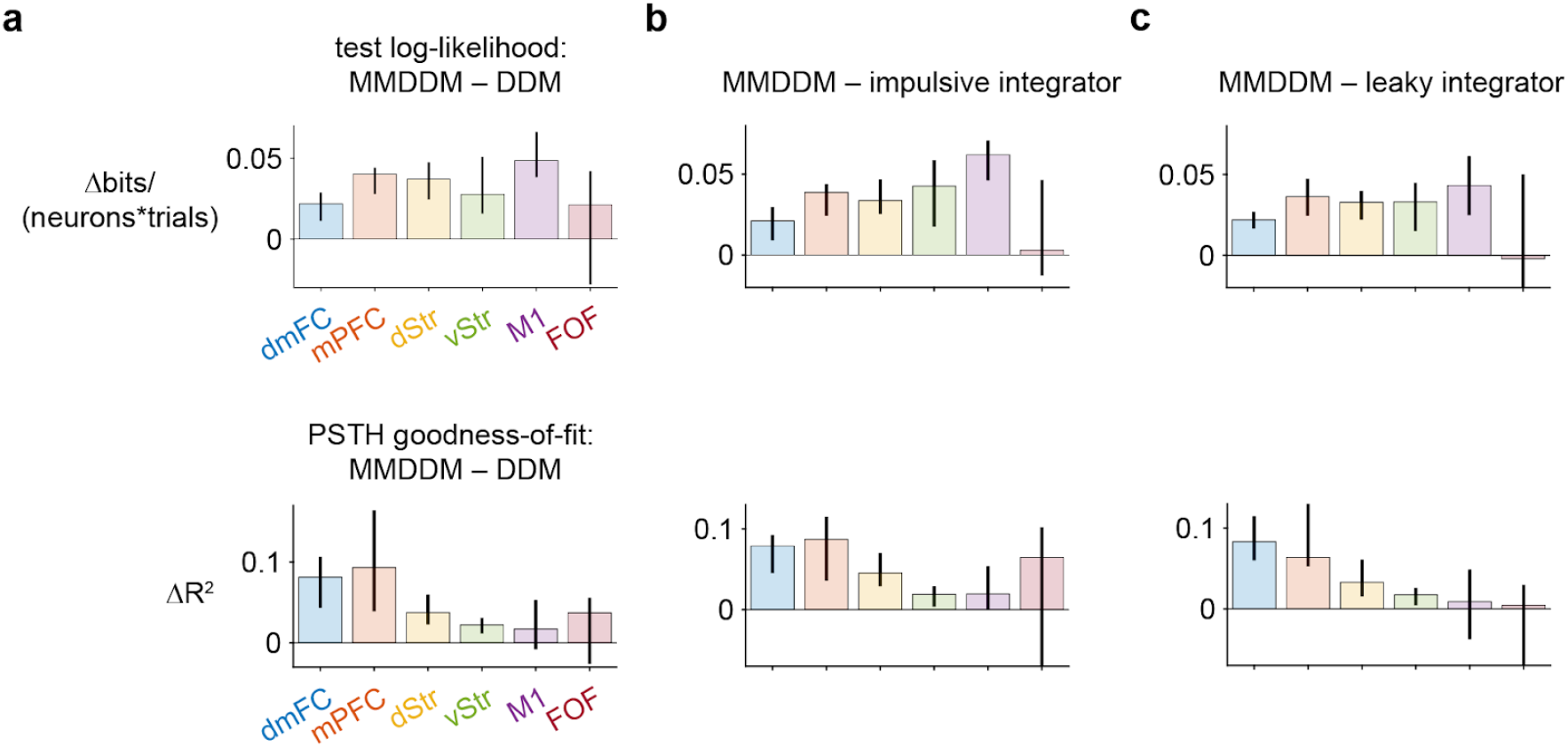
Dynamics across brain regions. **a**, Top, comparison of the out-of-sample log-likelihood between the MMDDM and the single mode DDM. Bottom, comparison of the out-of-sample R^2^ of the PSTH across brain regions. Error bar indicates 95% bootstrapped confidence intervals across sessions. **b**, Comparison between MMDDM and a single-mode impulsive integrator (an approximation of the bistable attractor hypothesis), which is implemented identically to the single-mode DDM except that the feedback parameter of the latent decision variable is not constrained to be zero but allowed to vary between 0 and 5 and is fit to the data. **c**, Comparison between MMDDM and bistable attractor model, which is implemented identically to the single mode DDM except that the feedback parameter of the latent decision variable is not constrained to be zero but allowed to vary between 0 and −5 and is fit to the data.

**Extended Data Figure 14.**
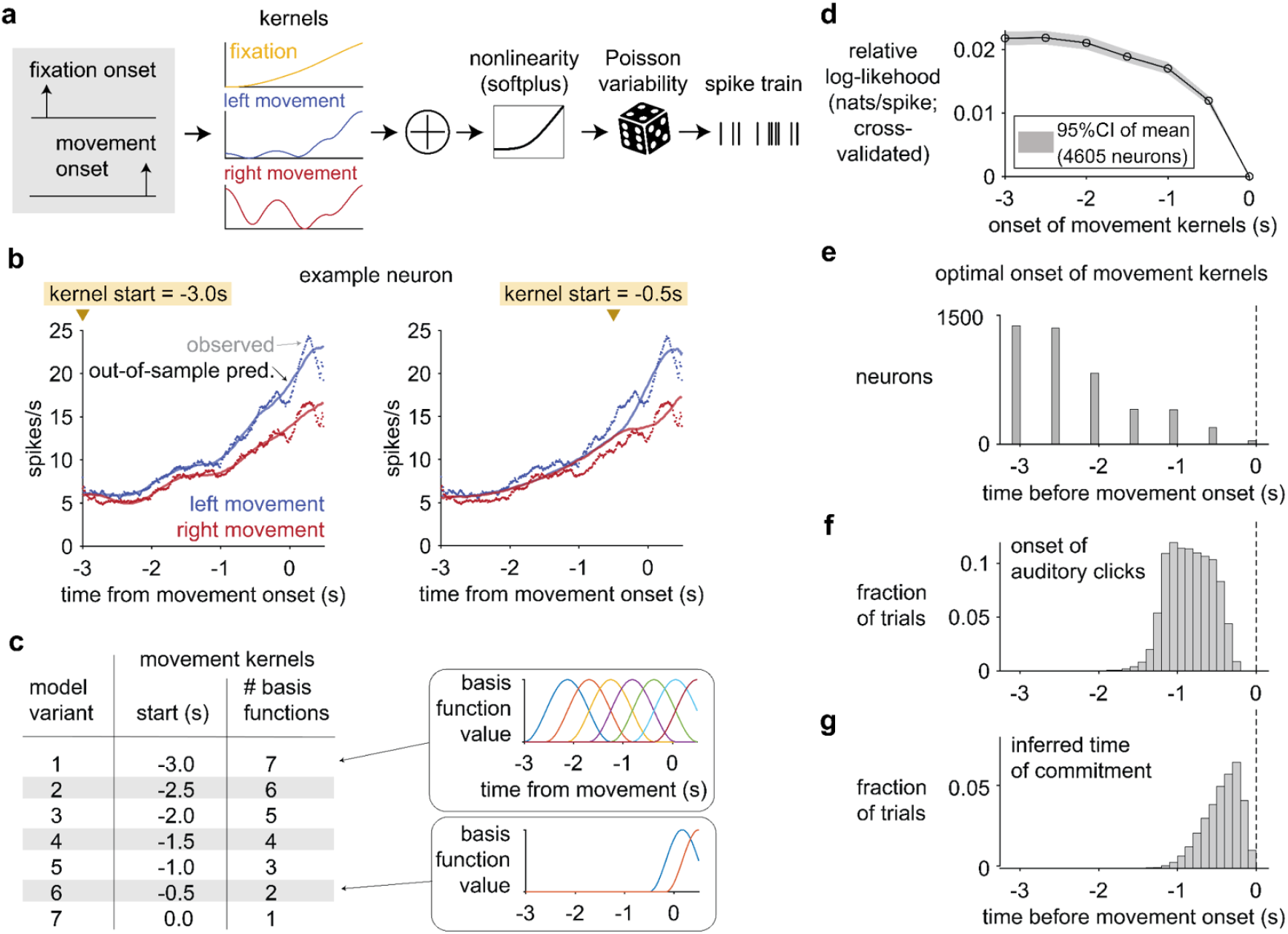
The distribution of commitment times inferred from MMDDM does not match the distribution of start time of peri-movement kernels. **a**, Separately for each choice-selective neuron (N=4605), peri-movement kernels are estimated using Poisson generalized linear models (GLM) ^54,56^. The inputs (i.e., regressors) to the model depend on two events that occur on each trial: onset of fixation (i.e., when the rat inserts its nose into the center port), and the time when the rat leaves the center port and begins to move toward the side port. An impulse (i.e., delta function) at the time of each event is convolved with a linear filter, or kernel, to parametrize the time-varying input related to that event. At each time step, the sum of the inputs is fed through a rectifying nonlinearity (softplus) to specify the neuron’s Poisson firing rate at that time. Three kernels, related to fixation, leftward movement, and rightward movement, are learned by maximizing the marginal likelihood ^54^. **b**, Example neuron. Two GLM variants were fitted to the same neuron, and for each GLM variant, the observed peri-event time histogram (PETH) is overlaid the cross-validated, model-predicted PETH. The choice-dependence of the PETH of this neuron is well captured by the model variant whose peri-movement kernels start −3.0s before and 0.5s after movement onset (left), but less well captured by another variant whose peri-movement kernels time base are limited to −0.5 to 0.5 s (right). **c**, To identify the optimal start of the movement kernel for each neuron, cross-validated (5-fold) model comparison was performed on seven model variants that vary in the start time of the movement kernels and the number of radial basis functions used to parametrize the kernels. The end time of the movement kernel (0.5s), and the parametrization of the fixation-related kernel (−1.5s to 2.0s and 4 basis functions) are identical for all variants. **d**, The out-of-sample log-likelihood is highest for the model variant whose peri-movement kernels start at −3.0 s. **e**, For each neuron, the GLM variant with the highest out-of-sample log-likelihood determines the optimal start of the peri-movement kernels. The mode of the distribution is at −3.0s. **f**, The start of peri-movement kernels for most neurons precede the time of the first click. **g**, The start of peri-movement kernels for most neurons precede the earliest commitment time inferred from MMDDM.

**Extended Data Figure 15.**
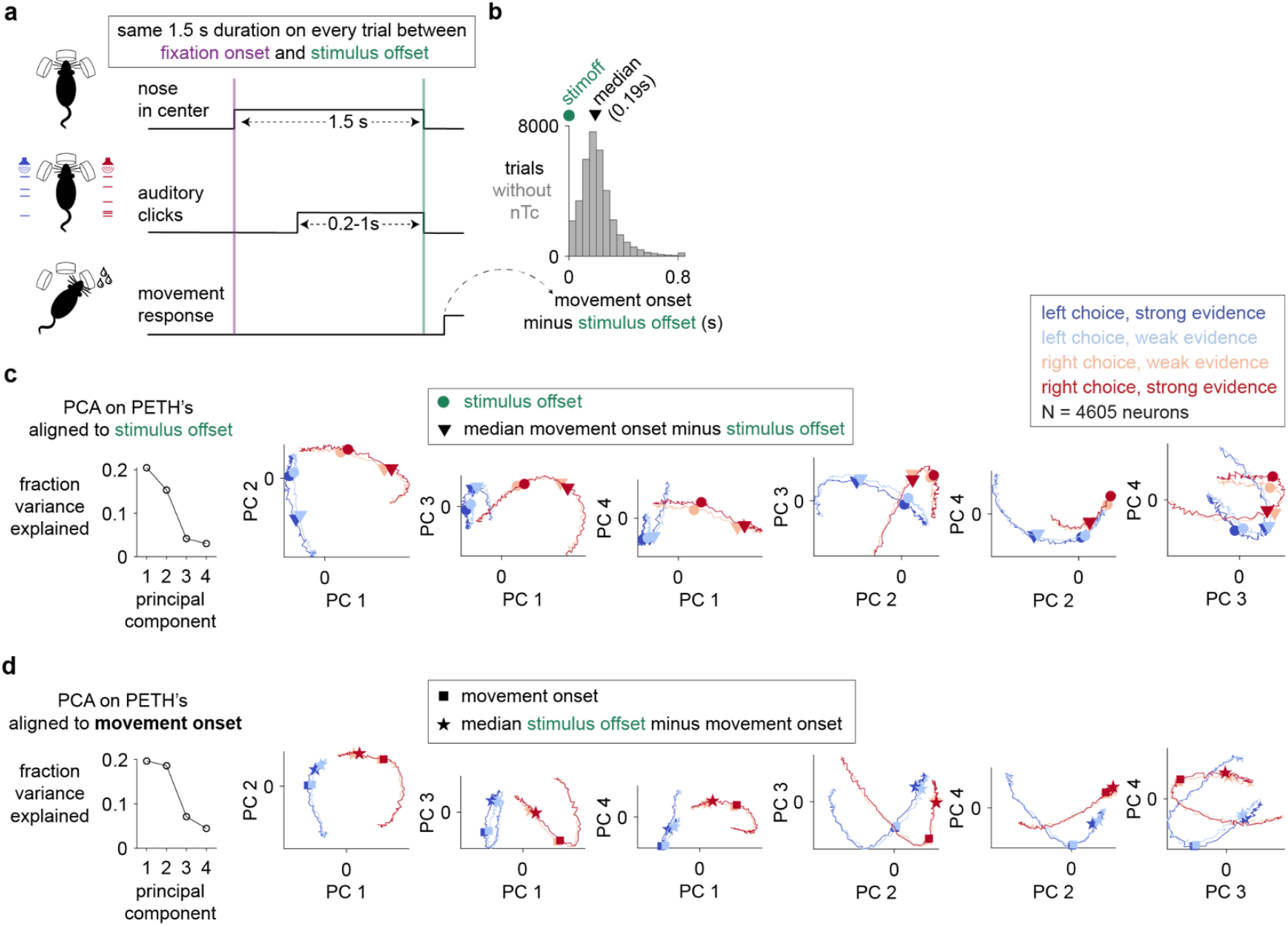
Changes in neural responses after stimulus offset are more closely aligned to movement onset than stimulus offset. **a**, Relative timing of task events. The offset of the auditory click train stimulus always occurred at the end of the 1.5 s minimum fixation period on every trial. **b**, The median time of movement onset relative to stimulus offset across trials without a neurally inferred time of commitment (nTc) is 0.192s. The rightmost bin contains trials for which the movement onset is 0.8s or more after stimulus offset. **c**, Principal component analysis (PCA) was performed on peri-event time histograms (PETH’s) aligned to stimulus offset (circles) and averaged across trials without a neurally inferred time of commitment (nTc). Spikes were counted in 10 ms bins, and the PETH was not additionally filtered. Spikes after the animal moved away from the fixation port (i.e., movement onset) were included. For each neuron and each trial condition, the PETH is a 100 element vector. Concatenating across 4605 choice-selective neurons and 4 trial conditions gave a 4605-by-400 matrix. The mean of each row (i.e., the average response of each neuron) was subtracted from the matrix, and PCA was performed on the resulting matrix. Triangles indicate the median time of movement onset. Projections are scaled by the standard deviation explained by each PC. **d**, PCA performed PETH’s aligned to movement onset offset and averaged across trials without nTc.

**Extended Data Figure 16.**
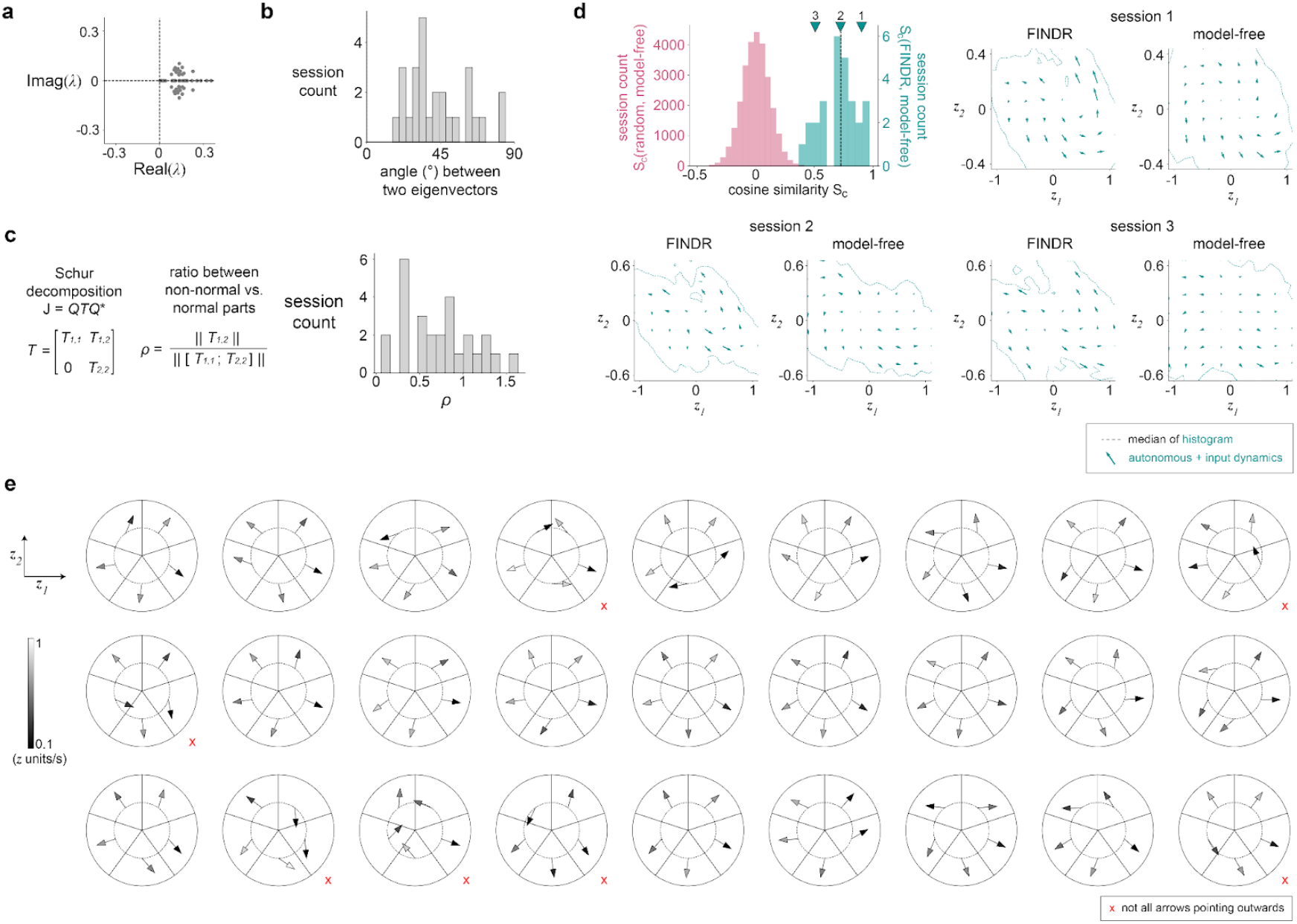
Further analyses and validation of the dynamics discovered by FINDR. **a**, When we computed the eigenvalues of the numerical Jacobian J obtained from the detected slow point around the origin, the real components of both eigenvalues were greater than zero for all sessions (*n*=27), indicating that the origin is not a stable point. Units of λ are sec^−1^. **b**, To quantify how non-normal the dynamics are around the origin, we computed the angle between the two eigenvectors of J. 90° indicates that the dynamics are normal, and angle less than 90° indicates that the dynamics are non-normal. **c**, We further evaluated the non-normality of the discovered dynamics around the origin by taking the Schur decomposition J = *QTQ*\* and computing the ratio between the non-normal part and the normal part of the dynamics, *ρ* = || *T*_1,2_ || / || [*T*_1,1_; *T*_2,2_] ||. *ρ* > 0 indicates that the dynamics are non-normal, with higher values of *ρ* indicating stronger non-normality. **d**, Here we estimated the low-dimensional vector field for each session using a method that does not specify a dynamical model (“model-free” approach). We compared the vector fields estimated using this approach to the FINDR-inferred vector fields. To obtain the model-free vector field, we first estimated single-trial firing rates of individual neurons by binning the spike trains in Δ*t* = 10ms bins and convolving the spike trains with a Gaussian of σ=100ms. Then, we projected the estimated single-trial population firing rate trajectories onto the subspace spanned by the FINDR latent axes. This allows direct comparisons between vector fields. For each evaluation point (*i, j*) on a 8-by-8 grid of the latent state space ***z***, we estimated the velocity arrow by taking the average of ***ż*** ≈ (***z***_*t*_ − ***z***_*t*-1_) / Δ*t* for all *t* across all trajectories that fall inside the cell corresponding to the point (*i, j*). To compare vector fields, we measured S_c_, the mean of the cosine similarities between the vector arrows of the model-free approach and the vector arrows from FINDR inside the sample zone. The median of the S_c_’s across all sessions was 0.73. Three example sessions from across the distribution are shown, with session 2 around the median S_c_ of the histogram. For both FINDR and the model-free approach, the colored trajectories were obtained by trial-averaging based on the evidence strength. To compare between a random vector field and the model-free vector field, for each session, we generated 1,000 random vector fields by randomizing the direction of each arrow in the 8-by-8 grid. **e**, We assessed the dynamical stability around the origin using a model-free approach similar to **d**. We estimated the autonomous velocity around the initial starting point (indicated as the center of the circle) of the model-free latent trajectories by taking the average of ***ż*** ≈ (***z***_*t*_ − ***z***_*t*-1_) / Δ*t* for all *t* across all trajectories that fall inside each of the 5 pie slices. Here we excluded time points where clicks affect the dynamics (***z***_*t*_ − ***z***_*t*-1_) / Δ*t*, and only considered the trajectories with #L clicks = #R clicks during the epoch when they are in the pie slice, when computing the estimate of the autonomous dynamics arrow. When computing the average of (***z***_*t*_ − ***z***_*t*-1_) / Δ*t* for one of the pie slices, we required ***z***_*t*-1_ to be inside the pie slice. The circles have a radius of 0.2 (in the units of ***z***). We found that all five arrows were pointing outwards (p<0.5^5^=0.03125) for 20 out of 27 sessions, consistent overall with the stability analysis in **a**.

**Extended Data Figure 17.**
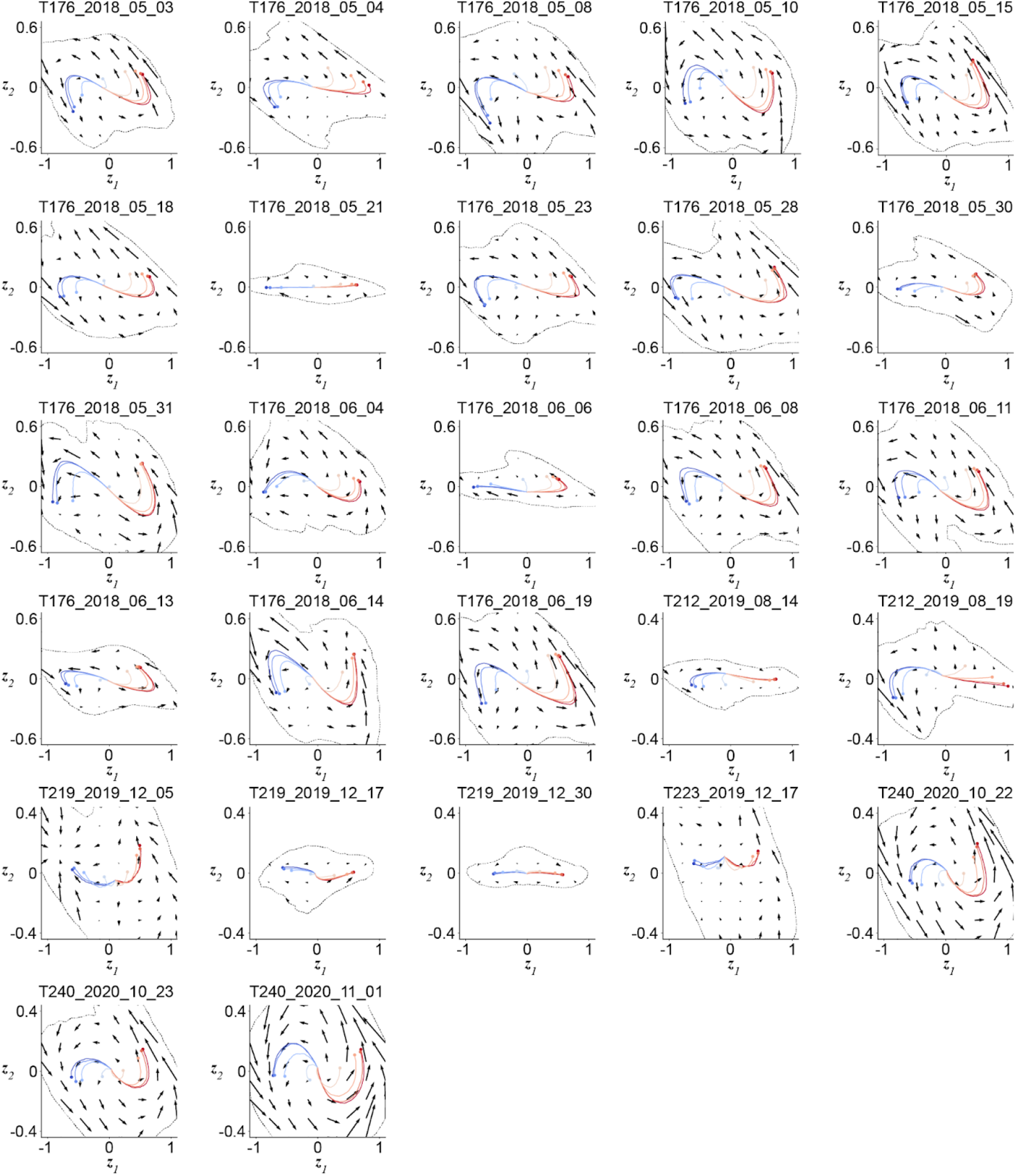
FINDR-inferred vector fields for all recording sessions (*n*=27) with than 30 neurons and 400 trials, and sessions where the animal performed with greater than 80% accuracy. These fits were used for the summary plots in **Fig**. 2k-l. The vector field represents the autonomous dynamics and the trajectories are trial averages sorted by the evidence strength of each trial.

## Supporting information

Detailed descriptions of modeling procedures

## Author contributions

T.Z.L. and D.G. collected data. T.D.K. developed the FINDR model. T.Z.L. developed the MMDDM model. T.Z.L. and T.D.K. developed the model of the baseline used in fitting FINDR and MMDDM. T.Z.L. and T.D.K. performed analyses. A.G.B. and C.D.K. assisted with data collection. V.A.E. and B.D. assisted with analyses. T.Z.L, T.D.K., and C.D.B. wrote the manuscript and conceptualized the study.

## Acknowledgments

We thank Julie Charlton, Long Ding, Joshua Gold, Robbe Goris, John Maunsell, and Jonathan Pillow for their suggestions and comments. We also thank Jessica Morrison, Klaus Osario, Jovanna Teran, and Emily Valance for technical assistance. Analyses reported in this work were substantially performed using the Princeton Research Computing resources at Princeton University which is a consortium of groups led by the Princeton Institute for Computational Science and Engineering (PICSciE) and Office of Information Technology’s Research Computing. This work was supported by grants from NIH F32 MH115416 and NIH R01MH108358.

## Data availability

The experimental data that support the findings of this study are available in Dryad with the identifier https://doi.org/10.5061/dryad.sj3tx96dm.

## Code availability

Custom acquisition, post-processing and analysis code is available at https://github.com/Brody-Lab/tzl_state_dependent_encoding. Code implementing FINDR^31^ is available at https://github.com/Brody-Lab/findr. Code implementing MMDDM is available at https://github.com/Brody-Lab/fhmddm, and code for estimating the baseline is available at https://github.com/Brody-Lab/tzl_spglm.

## Inclusion and Ethics Statement

The animal procedures described in this study were approved by the Princeton University Institutional Animal Care and Use Committee.

## 1 Experiments

### 1.1 Subjects

The animal procedures described in this study were approved by the Princeton University Institutional Animal Care and Use Committee and were carried out according to the standards of the National Institutes of Health. All subjects were adult male Long-Evans rats (Taconic, NY) that were housed in Technoplast cages in pairs and kept in a 12 hour reversed light-dark cycle. All training and testing procedures were performed during the dark cycle. The rats had free access to food, but they had restricted access to water. The amount of water that the rats obtained daily was at least 3% of their body weight.

### 1.2 Behavioral task

Rats performed the behavioral task in custom-made training enclosures (Island Motion, NY) placed inside sound- and light-attenuated chambers (IAC Acoustics, Naperville, IL). Each enclosure consisted of three straight walls and one curved wall in which three nose ports were embedded (one in the center and one on each side). Each nose port also contained one light emitting diode (LED) that was used to deliver visual stimuli, and the front of the nose port was equipped with an infrared (IR) beam to detect the entrance of the rat’s nose into the port. A loudspeaker was mounted above each of the side ports and used to present auditory stimuli. Each of the side ports also contained a silicone tube that was used for water reward delivery, with the amount of water controlled by valve opening time.

Rats performed an auditory discrimination task in which optimal performance required the gradual accumulation of auditory clicks [1]. At the start of each trial, rats inserted their nose in the central port and maintained this placement for 1.5 s (“fixation period”). After a variable delay of 0.5-1.3 s, two trains of randomly timed auditory clicks were presented simultaneously, one from the left speaker and one from the right speaker. At the beginning of each click train, a click was played simultaneously from the left and right speakers (“stereoclick”). Regardless of onset time, the click trains ended at the end of the fixation period, resulting in stimuli whose duration ranged from 0.2-1 s. The train of clicks from each speaker was generated by an underlying Poisson process, with different click rates for each side. The combined mean click rate was fixed at 40 Hz, and trial difficulty was manipulated by varying the ratio of the generative click rate between the two sides. The generative click rate ratio varied from 39:1 clicks/s (easiest) to 26:14 (most difficult). At the end of the fixation period, the rats could orient toward the nose port on the side where more clicks were played and obtain a water reward.

Psychometric functions were calculated by grouping the trials into eight bins of similar size according to the difference in the total number of right and left clicks and, for each group, computing the fraction of trials ending in a right choice. The confidence interval of the fraction of right response was computed using the Clopper-Pearson method.

### 1.3 Electrophysiological recording

Neurons were recorded using chronically implanted Neuropixels 1.0 probes that are recoverable after the experiment [6]. In each of four animals, a probe was implanted at 4.0 mm anterior to Bregma, 1.0 mm lateral, for a distance of 4.2 mm, and at an angle of 10 degrees relative to the sagittal plane that intersects the insertion site (the probe tip was more medial than the probe base). In each of five other animals, a probe was implanted to target primary motor cortex, dorsal striatum, and ventral striatum, at the site 1.0 mm anterior, 2.4 mm lateral, for a distance of 8.4 mm, and at an angle of 15 degrees relative to the coronal plane intersecting the insertion site (the probe tip was more anterior than the probe base). In each of three final sets of rats, a probe was implanted to target the frontal orienting fields and anterior dorsal striatum at 1.9 mm anterior, 1.3 mm lateral, for a distance of 7.4 mm, and at angle of −10 degree relative to the sagittal plane intersecting the insertion site (the probe tip was more lateral than the probe base). Spikes were sorted into clusters using Kilosort2 [9], and clusters were manually curated.

### 1.4 Muscimol inactivation

Infusion cannulas (Invivo1) were implanted bilaterally over dorsomedial frontal cortex (4.0 mm AP, 1.2 mm ML). After the animal recovered from surgery, the animal was anesthetized and on alternate days, a 600 nL solution of either only saline or muscimol (up to 150 nG) was infused in each hemisphere. Half an hour after the animal wakes up from anesthesia, the animal is allowed to perform the behavioral task.

### 1.5 Retrograde tracing

To characterize the anatomical inputs into dorsal striatum, 50 nL of cholera toxin subunit B conjugate (ThermoFisher Scientific) was injected into dorsal striatum at 1.9 mm AP, 2.4 ML, 3.5 mm below the cortical surface. The animal was perfused seven days after surgery.

### 1.6 Histology

The rat was fully anesthetized with 0.4 mL ketamine (100 mg/ml) and 0.2 mL xylazine (100 mg/ml) IP, followed by transcardial perfusion of 100 mL saline (0.9% NaCl, 0.3x PBS, pH 7.0, 0.05 mL heparin 10,000 USP units/mL), and finally transcardial perfusion of 250 mL 10% formalin neutral buffered solution (Sigma HT501128). The brain was removed and post fixed in 10% formalin solution for a minimum period of 7 days. 100 micrometer sections were prepared on a Leica VT1200S vibratome, and mounted on Superfrost Pus glass slides (Fisher) with Fluoromount-G (Southern Biotech) mounting solution and glass coverslips. Images were acquired on a Hamtasu NanoZoomer under 4x magnification.

## 2 Autonomous and input dynamics

The class of dynamical systems we study here is specified by

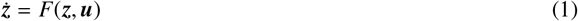

for some generic function *F*, with ***z*** the latent decision variable and ***u*** the external input to the system from the auditory clicks in the behavioral task. At each moment, there may be no click, a click from the left, or a right click. When time is discretized to sufficiently short steps, ***u*** is one of three values

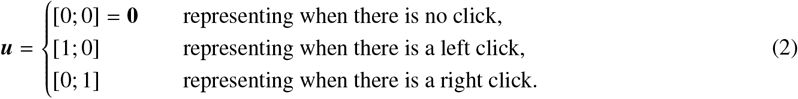

We define the autonomous dynamics of the system as

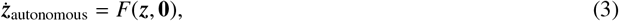

and the average input dynamics as

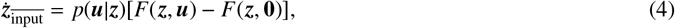

and specifically, the average left and right input dynamics as

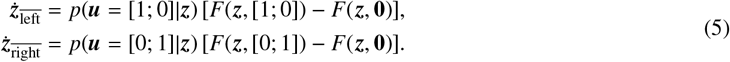

The sum of the autonomous dynamics and the average input dynamics is equal the expected value of ***ż*** computed over the distribution *p*(***u***|***z***):

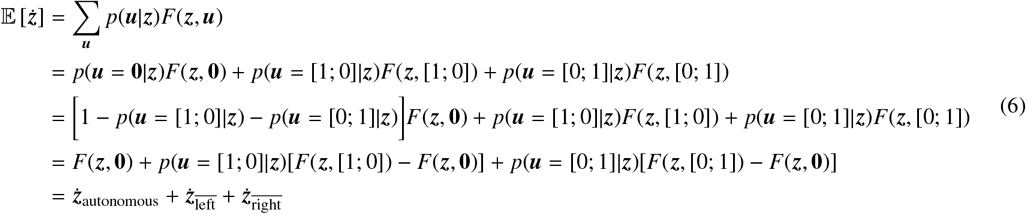

In the main text, Figure 2c plots ***ż***_autonomous_, and Figure 2e plots 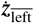 and 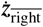. *F*(***z***, left) in the main text is defined as *p*(***u*** = [1; 0]|***z***)*F*(***z***, [1; 0]) + [1 − *p*(***u*** = [1; 0]|***z***)]*F*(***z*, 0**) and *F*(***z***, right) as *p*(***u*** = [0; 1]|***z***)*F*(***z***, [0; 1]) + [1 − *p*(***u*** = [0; 1]|***z***)]*F*(***z*, 0**).

Since *p*(***u***| ***z***) = *p*(***z***| ***u***)*p*(***u***)/*p*(***z***), and *p*(***z***) in general does not have an analytical form, we estimate *p*(***u z***) numerically. To do this, we train FINDR (Section 3, [4]) to learn *F*, and generate click trains for 5000 trials in a way that is similar to how clicks are generated for the task done by our rats. Then, we simulate 5000 latent trajectories from the learned *F* and the generated click trains. We then bin the state space of ***z*** and ask, for a single bin, how many times the latent trajectories cross that bin in total and how many of the latent trajectories, when crossing that bin had ***u*** = [1; 0] (or ***u*** = [0; 1]). That is, we estimate *p*(***u*** = [1; 0] ***z***) with 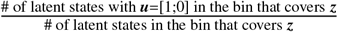. For Figure 2 in the main text, because ***z*** is 2-dimensional, we use bins of 8-by-8 that cover the state space traversed by the 5000 latent trajectories, and weigh the flow arrows of the input dynamics with the estimated *p*(***u*** |***z***). Similarly, for the background shading that quantifies the speed of input dynamics in Figure 2, we use bins of 100-by-100 to estimate *p*(***u*** |***z***), and apply a Gaussian filter with σ = 2 (in the units of the grid) to smooth the histogram. A similar procedure was performed in Extended Data Figures 3 and 6 to estimate *p*(***u***|***z***) numerically.

### 2.1 Speed of autonomous and input dynamics

To compute the normalised difference in the speed of autonomous and input dynamics in Figure 2k, similar to previous sections, we first generated latent trajectories from the learned *F* for 5000 different trials with generative click rate ratios used in our experiments with rats. Then, we computed the magnitude of the autonomous dynamics ||***ż***_autonomous_|| and the magnitude of the average input dynamics 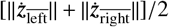 for each time point for each of the 5000 trajectories, and then averaged across the trajectories and across time periods defined in Figure 2j to obtain Figure 2k.

## 3 FINDR

Detailed descriptions are provided in [4]. Briefly, to infer velocity vector fields (or flow fields) from the neural population spike trains, we used a sequential variational autoencoder (VAE) called Flow-field Inference from Neural Data using deep Recurrent networks (FINDR).

FINDR minimizes a linear combination of two losses: one for neural activity reconstruction (ℒ _1_) and the other for vector-field inference (ℒ _2_). To reconstruct neural activity, FINDR uses a deep neural network *G* that takes the spike trains of *N* simultaneously recorded neurons ***y*** and the sensory click inputs ***u*** on a given trial, and reconstructs ***y*** from the *d*-dimensional latent decision variable ***z***:

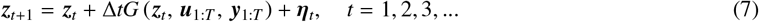

Here, *T* is the number of time steps on a given trial, ***u***_*t*_ is a two-dimensional vector representing the number of left and right clicks played on a time step (Δ*t* = 0.01s), ***y***_*t*_ an *N*-dimensional vector of the spike counts on a time step, and η_*t*_ noise drawn from *N* (**0**, Δ*t*Σ) on each time step. Σ is a *d*-dimensional diagonal matrix, where the diagonal elements need not be equal to each other. At each time step, firing rates of *N* simultaneously recorded neurons ***r***_*t*_ are given by

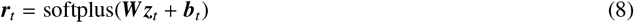

where softplus is a function approximating the firing rate-synaptic current relationship (f-I curve) of neurons, ***W*** a × *N d* matrix representing the encoding weights, ***b***_*t*_ a *N*-dimensional vector representing the putatively decision-irrelevant baseline input. The baseline ***b***_*t*_ is learned prior to fitting FINDR using the procedure described in Section 5.4 and in detail in the Supplementary Information. The reconstruction loss is given by

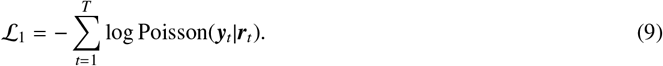

For vector field inference, we parametrize the vector field *F* with a gated feedforward neural network [3, 4]:

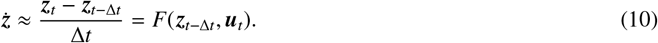

*F* gives the discretized time derivative of ***z***. We find the vector field *F* that captures the latent trajectories ***z*** inferred from *G* in Eq. (7) by minimizing

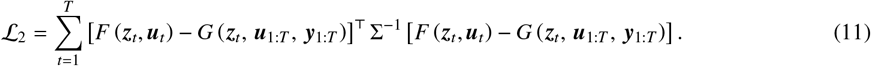

The total loss that is minimized by FINDR is

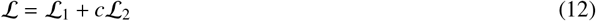

where *c* = 0.1 is a fixed hyperparameter (*c* = 0.0125 in Extended Data Figure 3b). FINDR minimizes ℒ by using stochastic gradient descent (SGD) to learn ***W***, Σ, the parameters of the neural network representing *F*, and the parameters of the neural network *G*. It can be shown that ℒ is an approximate upper bound on the marginal log-likelihood of the data, and that training FINDR this way is equivalent to performing inference and learning via a sequential auto-encoding variational Bayes (AEVB) algorithm that straightforwardly extends the standard AEVB [5, 4].

After training, we plot the vector field (i.e., a grid of ***ż***) using the learned *F*, and generate FINDR-predicted neural responses using Eq. (8) and

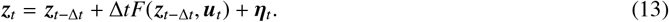

Eq. (13) is an Euler-discretized gated neural stochastic differential equation (gnSDE [3, 4]).

### 3.1 Parameters

The total number of free parameters *P* of the FINDR model is given by

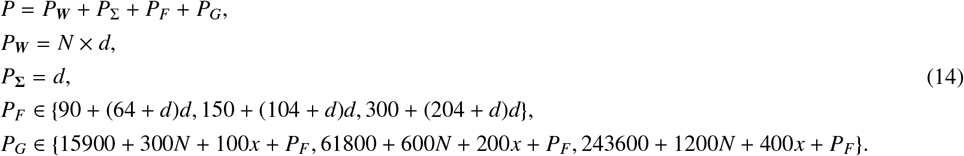

*P*_***W***_ is the number of parameters in the encoding weight matrix ***W***, whose dimensions are the number of neurons *N* and latent dimensionality *d. P*_Σ_ is the parameter count in the diagonal covariance Σ of the additive Gaussian noise of the latent ***z***. The number of parameters in the neural networks parametrizing *F* (*P*_*F*_) and *G* (*P*_*G*_) are separate hyperparameters. Here, 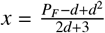.

### 3.2 Hyperparameters

The hyperparameters that were optimized (*P*_*F*_, *P*_*G*_, α) include the number of parameters of the network *F* (*P*_*F*_), the number of parameters of the network *G* (*P*_*G*_), and the learning rate α ∈ {10^−2^, 10^−1.625^, 10^−1.25^, 10^−0.875^, 10^−0.5^}. We identify the optimal values for these hyperparameters in 3 × 3 × 5 = 45 grid search. The grid search was done separately for each set of training data for each of five cross-validation folds. Within each each training set, 3/4 of the trials were used to the optimize the parameters under a given set of hyperparameters, and the remaining 1/4 were held out to evaluate the model’s performance for that set of hyperparameters. Test data were never used in the grid search.

### 3.3 Latent space transformation

Because the encoding weight matrix ***W*** is not constrained to semi-orthogonal and can take only any real values, different combinations of ***W*** and ***z***_*t*_ can give rise to the same firing rate vector ***r***_*t*_, even when baseline ***b***_*t*_ is fixed. To uniquely identify the latent trajectories (except for redundancy from rotations and reflections), after optimization, we linearly transformed the latent space ***z*** to 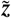

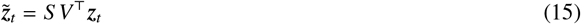

where *S* a *d* × *d* diagonal matrix containing the singular values of ***W*** and *V* a *d* × *d* matrix containing the right singular vectors

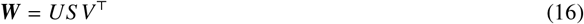

*U* is an *N* × *d* matrix containing the left singular vectors of ***W*** (where *N* is the number of neurons). In the space of 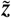, the encoding weight matrix is a linear transformation that preserves angles and distances because *U* is semi-orthogonal and can only give rise to an isometry such as rotation and reflection.

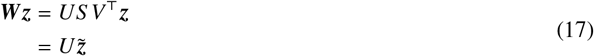

To obtain meaningful axes for the transformed latent space 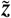, we generate 5000 different trajectories of 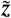 in generative mode (i.e., using *F* and Σ in Eq. (13), but not *G*), and perform principal component analysis (PCA) on the trajectories. The principal components (PCs) were used to define the axes of the decision variable 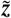. In the main text, the PC 1 axis of 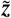 was denoted as ***z***_1_, and the PC 2 axis of 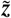 was denoted as ***z***_2_. In all our analyses, the latent trajectories and vector fields inferred by FINDR are shown in the transformed latent space of 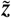, and scaled so that the latent trajectories along PC 1 lie between −1 and 1.

### 3.4 Model evaluation

The goodness-of-fit of the PSTH was quantified using the coefficient of determination (*R*^2^) of the evidence-sign conditioned PSTH as defined in Eq. (33), using five-fold cross-validation. We used 3/5 of the trials in a session as training dataset, 1/5 of the trials as validation dataset to optimize the hyperparameters of FINDR, and 1/5 of the trials as test (i.e., out-of-sample) dataset to evaluate performance of FINDR. Therefore, when we compute the goodness-of-fit, we also obtain 5 different vector fields inferred by FINDR for each fold, which we confirm are consistent across folds (Extended Data Figure 6).

### 3.5 FINDR models with latent dimensions greater than two

We evaluated FINDR models with latent dimensions higher than two to assess whether the two-dimensional manifold we found is approximately an attractor. To show that the sample zone is an approximate attractor manifold, we perturb the latent states on the manifold along the 3rd principal component (PC) direction. When the latent states are perturbed (but not too far that the latent states go outside the range along the PC 3 axis covered by the sample zone), the latent states flow towards the manifold. To get the flow directions along PC 3, we first generated 5000 latent trajectories (similar to Figure 2 for computing the sample zone). We then divided the PC 1 × PC 2 space into an 8-by-8 grid (the grid used for the vector field arrows in Extended Data Figure 5e). For each cell of the grid, we identified the latent states from the 5000 trajectories that were inside the cell, and identified the highest (lowest) PC 3 value 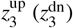. This was to ensure that the perturbation along the PC 3 axis was not too large. Then we computed the flow vector using a 100-by-100 grid on the PC 1 × PC 2 space, assuming that 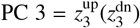 and PC 4 = 0. The space covered by each cell of the grid is colored based on the direction of the flow vector along PC 3—if flowing upwards green, if flowing downwards pink. This heatmap was applied a Gaussian filter with σ = 2 (in the units of the 100-by-100 grid), similar to the heatmap for input dynamics in Figure 2f. The resulting plot is shown on the left (right) panel. Results were similar without the Gaussian filter.

### 3.6 Constrained FINDR (cFINDR)

The constrained FINDR model (cFINDR) replaces the neural network parametrizing *F* in FINDR with a linear combination of affine dynamics, specified by ***M*** and ***N***, and bistable attractor dynamics specified by *φ*. The dynamics are furthermore constrained to be two-dimensional.

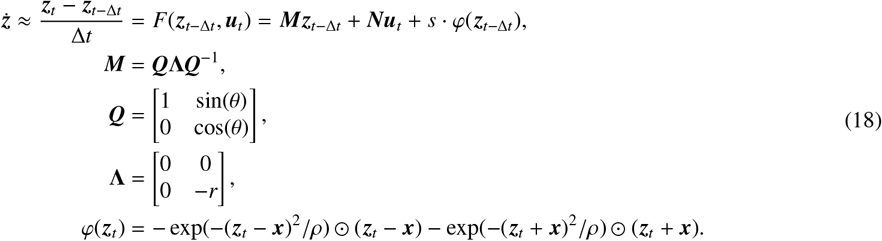

The matrix ***M*** implements a line attractor located at *z*_2_ = 0. The inputs ***u***_*t*_ are the same as in FINDR and represent the auditory clicks. The two discrete attractors are constrained such that *x*_2_ = 0 and implemented through the function *φ*. The shape of the basin of attraction corresponding to each point attractor is specified by the parameter *ρ*. The relative contribution of the discrete attractors and the line attractor to the overall dynamics is specified by the scalar *s*.

The DDM line attractor hypothesis can be implemented in cFINDR by setting *θ* = 0. Non-normal dynamics with a line attractor [7] can be implemented by setting *θ* ≠ 0. The bistable attractor hypothesis can be implemented by increasing *ρ*.

As in FINDR, cFINDR learns ***W***, Σ, and parameters of *G*. Instead of the neural networks parametrizing *F*, cFINDR, learns *s, θ, r*, ***x***, *ρ*, and the 2 × 2 matrix ***N*** to approximate *F*, which has nine parameters. The same objective function and optimization procedure were used in cFINDR. After optimization, as in FINDR, the latent space ***z*** is linearly transformed to uniquely identify the dynamics (except for arbitrary rotations or reflections). As in the analysis of the results from FINDR, the latent trajectories and vector fields inferred by cFINDR are in the transformed latent space 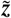.

When we fit cFINDR to the data, we experimented with the different constraints *r* > 0 and *r* > 3. The fits using *r* > 0 were superior to those using *r* > 3 and therefore used in the comparison between cFINDR and FINDR of the data presented in Extended Data Figure 7h-i. The motivation to try both *r* > 0 and *r* > 3 was because we found that in synthetic data, cFINDR under the constraint *r* > 0 could not recover the dynamics generated under the DDM line attractor hypothesis (*r* = 10). For this reason, Extended Data Figure 7f shows results from synthetic data using *r* > 3. When fit to data, FINDR outperforms cFINDR using either *r* > 0 or *r* > 3.

### 3.7 Curvature of trial-averaged trajectories

To compute the curvature of trial-averaged trajectories in Figure 2j, as before, we first generate latent trajectories from FINDR for 5000 different trials with generative click rate ratios used in our experiments with rats. Then, we separate the trials based on whether the generative click ratio on a given trial favors a leftward choice or a rightward choice. We take the average of the latent trajectories over the left-favoring trials, and then convolve the trial-averaged trajectory with a Gaussian filter with σ = 3 (in the units of the time step Δ*t* = 0.01s). We take this smoothed trajectory to numerically compute the planar curvature. We do the same for the right-favoring trials, and take the average between the curvature obtained from left-favoring trials and curvature obtained from right-favoring trials to generate the plot in Figure 2j.

### 3.8 Choice decoding from FINDR

FINDR does not use animal’s choice for reconstructing neural activity. However, after training, we can fit a logistic regression model that predicts the animal’s choice from the decision variable ***z*** at the final time step *T*. When we fit an *l*_2_-regularized logistic regression model using ***z***_*T*_ from the trained network *G* and the animal’s choice on the representative session in Figure 2c-i, we found that the logistic choice decoder achieves 89.7% accuracy in predicting choice on the out-of-sample dataset. We can generate choices from this decoder by generating latent trajectories using *F* and Σ in Eq. (13) as in previous Sections, and supplying ***z***_*T*_ to the trained decoder. 5000 latent trajectories and choices generated from *F* and the choice decoder were used for the analysis in Extended Data Figure 6l. We used a separate logistic regression model for predicting choice from the latent trajectories truncated at time = 0.33s projected onto PC 2. The optimization of the logistic regression models was done using L-BFGS [2].

## 4 MMDDM

The multi-mode drift-diffusion model (MMDDM) is a state-space model, comprising a dynamic model that governs the time evolution of the probability distributions of latent (i.e., hidden) states and measurement models that define the conditional distributions of observations (i.e., emissions) given the latent state.

### 4.1 Dynamic model

The latent variable *z* is one-dimensional (i.e., a scalar), and its time evolution is governed by a piecewise linear function:

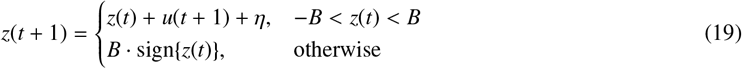

When the absolute value of *z* is less than the bound height *B* (free parameter), its time evolution depends on momentary external input *u* and i.i.d. Gaussian noise *η*.

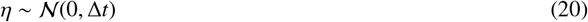

where Δ*t* is the time step and set to be 0.01s (10 milliseconds). When *z* is either less than −*B* or greater than *B*, it becomes to be fixed at the bound. The initial probability distribution of *z* is given by

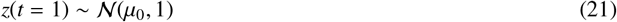

whose mean *µ*_0_ is a free parameter. On time step *t*, the input *u*(*t*) is the total difference in the per-click input *v* between the right and left clicks that occurred in the time interval [*t* − Δ*t, t*)

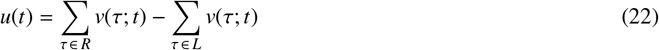

where *L* (*R*) is the set of the left (right) click times and *v*(τ; *t*) is the per-click input of a click occurred time τ at time step *t*. Note that τ ∈ ℝ indicates continuous time, whereas *t* ∈ ℕ indexes a time step. The per-click input is given by

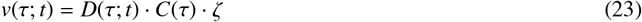

where *D*(τ; *t*) indicates the integral over the interval [*t* − Δ*t, t*) of the Direct delta function *δ* delayed by τ:

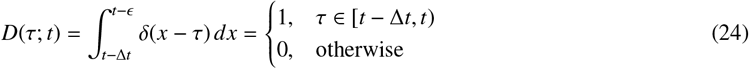

where *ϵ* is the machine epsilon. To account for sensory adaptation, the per-click input is depressed by preceding clicks by a time-varying scaling factor given by the function *C*(τ), implemented according to previous work [1] (Supplemental Information). The per-click input is corrupted by i.i.d. multiplicative Gaussian noise *ζ*:

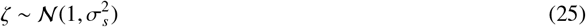

The free parameter 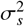 is the variance of the per-click noise. Variability in the dynamic model is fit to the data through the per-click noise *ζ* rather than per-time step noise *η* based on previous findings [1]; our results are similar if we set the variance of *η* rather than the variance of *ζ* to be a free parameter.

The dynamic model has 3 free parameters: bound height *B*, variance 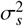 of the per-click noise, and mean *µ*_0_ of the initial state. These parameters are learned simultaneously with the parameters of the measurement models.

### 4.2 Measurement model of behavioral choices

On each trial, the binary behavioral choice *c* (1=right, 0=left) is the sign of *z* on the trial’s last time step *T* (the earlier of 1 s after the onset of the clicks or immediately before the animal leaves the fixation port):

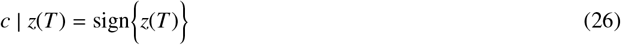

### 4.3 Measurement model of spike counts

On each time step *t*, given the value of *z*, the spike count *y* of neuron *n* is a Poisson random variable

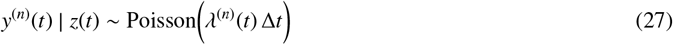

The firing rate *λ* is given by

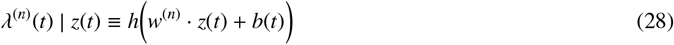

where *h*(·) is the softplus function used to approximate the neuronal frequency-current curve of a neuron:

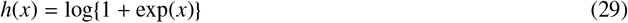

The encoding weight *w* depends on *z* itself:

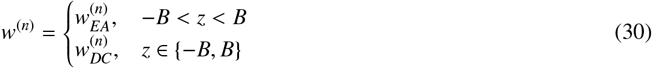

Each neuron has two scalar weights, *w*_*EA*_ and *w*_*DC*_, that specify the encoding of the latent variable during the evidence accumulation regime (EA) and the decision commitment regime (DC), respectively. When the latent variable has not yet reached the bound (−*B* or *B*), all simultaneously recorded neurons are in the evidence accumulation regime and encode the latent variable through their own private *w*_*EA*_. When the bound is reached, all neurons transition to the decision commitment regime and encode *z* through their own *w*_*DC*_.

The bias *b* accounts for factors that are putatively independent of the decision, including a component that varies only across trials and another component that varies both across and within trials:

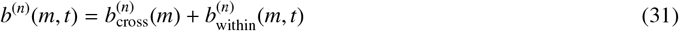

The cross-trial trial component, 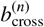 is a function of time from the first trial of the session, whereas *t* indicates time within each trial relative to that trial’s stimulus onset. The within-trial component consists of time-varying influence from spike history, post-stimulus onset, and pre-movement onset.

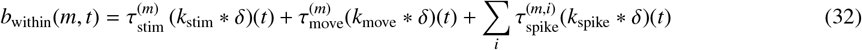

where the symbol * indicates convolution, τ_*x*_ indicates translation τ_*x*_*k*(*t*) = *k*(*t* − τ_*x*_) by the time of event *x*, and *δ* is the Dirac delta function. The functions 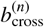, *k*_stim_, *k*_move_, *k*_spike_ are learned and each parametrized as a linear combination of radial basis functions, following [10, 11] (Supplemental Information). Each neuron’s measurement model of the spike train has 19 parameters that are learned simultaneously with the parameters of the dynamic model (i.e., model of the latent variable).

### 4.4 Parameter learning

All parameters, including the 3 parameters of the latent variable, as well as the 19 parameters private to each neuron, are learned simultaneously by jointly fitting to all spike trains and choices using maximum a posteriori estimation. Gaussian priors were placed on the model parameters to ensure the optimization reached a critical point and confirmed to not change the results in separate optimizations using maximum likelihood estimation (i.e., optimization without Gaussian priors). Out-of-sample predictions were computed using five-fold cross-validation.

### 4.5 nTc

The time step when decision commitment occurred is selected to be when the posterior probability of the latent variable at either the left or the right bound, given the click times, spike trains, and behavioral choice, is greater than 0.8. Results were similar for other thresholds, and the threshold of 0.8 was chosen to balance between the accuracy of the prediction and the number of trials for which commitment is predicted to have occurred. Using this definition, commitment occurred in 34.6% of the trials.

### 4.6 Engagement index

The engagement index (*EI*) was computed for each neuron to quantify its involvement in evidence accumulation and decision commitment. The index was defined using the neuron’s *w*_*EA*_ and *w*_*DC*_: *EI* ≡ (|*w*_*EA*_|− |*w*_*DC*_|)/(|*w*_*EA*_| + |*w*_*DC*_|). It ranges from −1 to 1. A neuron with an *EI* of −1 encodes the latent variable only during decision commitment, an *EI* of 1 indicates involvement only during evidence accumulation, and an *EI* of 0 represents a similar strength of encoding the latent variable during evidence accumulation and decision commitment.

## 5 Analyses

### 5.1 Neuronal selection

Only neurons that meet a pre-selected threshold for being reliably choice-selective are included for analysis. For each neuron, reliable choice selectivity is measured using the area under the receiver operating characteristic (auROC) indexing how well an ideal observer can classify between a left- and right-choice trial based on the spike counts of the neuron. Spikes were counted in four non-overlapping time windows (0.01-0.21 s after stimulus onset, 0.21-0.4, 0.41-0.6, and 0.61-0.9), and an auROC was computed for each time window. A neuron with an auROC < 0.42 or > 0.58 for any of these windows is considered choice selective and included for other analyses. Additionally, neurons must have an average firing rate of at least 2 spikes/s. Across sessions, the median fraction of neurons included under this criterion is 10.4%.

### 5.2 Peri-stimulus time histogram (PSTH)

Spike times were binned at 0.01 s and were included up to one second after the onset of the auditory stimulus (click trains) until one second after the stimulus onset, or until when the animal removed its nose from the central port, whichever came first. The time-varying firing rate of each neuron in each group of trials (i.e., task condition) was estimated with a peristimulus time histogram (PSTH), which was computed by convolving the spike train on each trial with a causal Gaussian linear filter with a standard deviation of 0.1 s and a width of 0.3 s and averaging across trials. The confidence interval of a PSTH was computed by bootstrapping across trials.

The goodness-of-fit of the model predictions of the PSTH was quantified using the coefficient of the determination (*R*^2^), computed using five-fold cross-validation. The *R*^2^ was computed using the by conditioning the PSTH on either the sign of the evidence (i.e., whether the generative click ratio on a given trial favors a leftward choice or a rightward choice) or the animal’s choice:

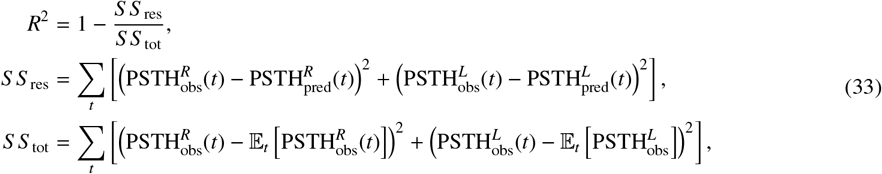

where *t* is time within a trial that goes from 0s to 1s, with 0s being the stimulus onset. The superscripts *L* and *R* indicate either the sign of the difference in the total number of right and left clicks or the animal’s choice. Definitions for 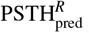 and 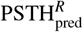 are similar, but for the model-predicted neural activity. When conditioning on the animal’s choice, *L* and *R* refer to the left and right choices of the animal, respectively.

A normalised PSTH was computed by dividing the PSTH by the mean firing rate of that neuron across all time steps across all trials. When PSTH’s were separated by “preferred” and “null”, the “preferred” task condition was defined as the group of trials with the behavioral choice when the neuron responded more strongly, and a “null” task condition was defined as the trials associated with the other choice.

### 5.3 Choice selectivity

In Figure 5b and Extended Data Figure 9i, for each neuron, and for each time step *t* aligned to the onset of the auditory click trains, we computed choice selectivity *c*(*t*):

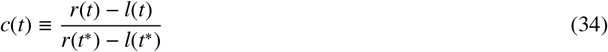

where *r* and *l* are the PSTHs computed from trials ending in a right and left choice, respectively. The time step *t*^*^ is time of the maximum absolute difference:

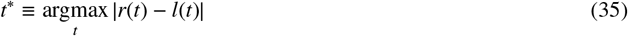

In Extended Data Figure 9i, neurons are sorted by the center of mass of each neuron’s absolute value of the choice selectivity.

### 5.4 Baseline

In FINDR, cFINDR, and MMDDM, the firing rate of a neuron depends on a time-varying scalar baseline. On timestep *t* of trial *m*, conditioned on the value of the latents on a given time step, the spike count *y* of each neuron is given by

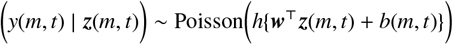

where *h* is the softplus function and ***w*** is the encoding weight of the latent. The baseline *b* incorporates putatively decision-independent variables as input to the neural spike trains including slow drifts in firing rates across trials and faster changes within each trial that are aligned to either the time from stimulus onset or the time from the animal leaving the fixation port. The baseline is learned using a Poisson generalized linear model fit separately to the spike counts of each neuron. Details are provided in the Supplementary Information.

### 5.5 Peri-commitment time histograms (PCTH)

On trials for which a time of decision commitment (nTc) could be inferred (see section 4.5), the spike trains are aligned to the predicted time of commitment and then averaged across those trials. The trial-average is then filtered with a causal Gaussian kernel with a standard deviation of 0.05s. The PCTHs were averaged within each of three groups of neurons: 1) neurons that are similarly engaged in evidence accumulation and choice maintenance; 2) neurons more strongly engaged in evidence accumulation; and 3) neurons more strongly engaged in choice maintenance. Each neuron was assigned to one of these groups according to its engagement index (*EI*; see section 4). Neurons with 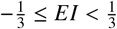 are considered to be similarly engaged in evidence accumulation and choice maintenance; neurons with 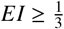 are considered to be more strongly engaged in evidence accumulation; and those with 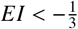 are considered to be more strongly engaged in choice maintenance. Principal component analysis on the PCTHs is performed as described in section 5.6.

For this analysis, we focused on only the 65/115 sessions for which the MMDDM improves the *R*^2^ of the PSTHs and also that the inferred encoding weights are reliable across cross-validation folds (*R*^2^ > 0.9). From this subset of sessions, there were 1,116 neurons similarly engaged in maintenance and accumulation, 414 neurons that are more engaged in maintenance, and 1,529 neurons more engaged in accumulation.

To compute the shuffled PCTH, the predicted times of commitment are shuffled among only the trials on which commitment were detected. If the randomly assigned commitment time extends beyond the length of the trial, then the time of commitment is assigned to be the last time step of that trial.

### 5.6 Trial-averaged trajectories in neural state space

To measure the trial-averaged dynamics in neural state space, principal component analysis (PCA) was performed on a data matrix made by concatenating the peri-stimulus time histograms. The data matrix *X* has dimensions *T* × *C*-by-*N*, where *T* is the number of time steps (*T* = 100), *C* is the number of task conditions (*C* = 2 for choice-conditioned PSTHs, *C* = 4 for PSTHs conditioned on both choice and evidence strength), and *N* is the number of neurons. The mean across rows is subtracted from *X*, and singular value decomposition is performed: *US V*^⊤^ = *X*. The principal axes correspond to the columns of the right singular matrix *V*, and the projections of the original data matrix *X* onto the principal axes correspond to the left singular matrix (*U*) multiplied by *S*, the rectangular diagonal matrix of singular values. The first two columns of the projections *US* are plotted as the trajectories in neural state space.

### 5.7 Psychophysical kernel

Kernels were time locked to either each trial’s nTc (Fig. 3n; Extended Data Fig. 10a-d) or the first click on each trial (Extended Data Fig. 10e-h). We extended the logistic regression model presented in [8] to include a lapse parameter (Supplemental Information), and we confirmed that results were similar using generic logistic regression. A shuffling procedure was used to randomly permute the inferred time of commitment across trials, without changing the behavioral choice and the times of the auditory clicks on each trial. Within this randomly permuted sample, we selected trials for which the auditory stimuli were playing at least 0.2s before and at least 0.2s after the inferred time of commitment to compute the psychophysical kernel in the “shuffled” condition. In Fig. 3n, the prediction was generated using the MMDDM parameters that were fit to the data and the same set of trials in the data. In Extended Data Fig. 10, temporal basis functions were used to parametrize the kernel, and the optimal number and type of basis functions were selected used cross-validated model comparison.

### 5.8 Statistical tests

Binomial confidence intervals were computed using the Clopper-Pearson method. All other confidence intervals were computed through a bootstrapping procedure using the bias-corrected and accelerated percentile method [12]. Unless otherwise specified, *P*-values comparing medians were computed using a two-sided Wilcoxon rank sum test, which tests the null hypothesis that two independent samples are from continuous distributions with equal medians, against the alternative that they are not.

### 5.9 Estimating the low-dimensional vector field without specifying a dynamical model

In Extended Data Figure 16d, we estimated the low-dimensional velocity vector field for each session using a method that does not specify a dynamical model (“model-free” approach). To obtain the model-free vector field, we first estimated single-trial firing rates of individual neurons by binning the spike trains into Δ*t* = 10ms bins, and convolving the spike trains with a Gaussian of σ = 100ms centered at 0. Results were similar for other values of σ around 100ms. Then, for each neuron we took the average across all trials in the session and subtracted this average from single-trial firing rate trajectories. These baseline-subtracted firing rate trajectories were then projected to the low-dimensional subspace spanned by the FINDR latent axes. We projected the estimated firing rates to the same subspace as FINDR to allow direct comparisons between the FINDR-inferred vector field and the model-free vector field.

We treated this low-dimensional projection of the baseline-subtracted firing rates as the latent trajectories in this model-free approach. To obtain velocity vector fields from the latent trajectories, we first estimated the instantaneous velocity ***ż*** at time point *t* by computing ***ż***_*t*_ = (***z***_*t*_ − ***z***_*t*−Δ*t*_)/Δ*t* for all *t* for all latent trajectories. We then divided the two-dimensional latent space into an 8-by-8 grid. For each cell (*i, j*) from this 8-by-8 grid, we identified all states ***z***_*t*_ from all trajectories that fall inside the cell (*i, j*). We took the corresponding ***ż***_*t*_ of the identified ***z***_*t*_’s and took the average to compute the velocity for the cell (*i, j*). We computed the velocity vectors for all 64 cells. To compare vector fields, we took the cosine similarity between the velocity vector for cell (*i, j*) from FINDR and the velocity vector for cell (*i, j*) from the model-free approach, and took the mean of these cosine similarities, *S* _*c*_(FINDR, model-free). In computing *S* _*c*_(FINDR, model-free), only cells that had the number of states greater than 1% of the total number of states were included. When the number of states used to estimate the velocity vector was less than 1% of the total number of states, we considered that cell (*i, j*) to be outside the “sample zone”, analogous to the sample zone in Figure 2.

To compare between a random vector field and the model-free vector field, we generated 1,000 random vector fields (with each of the 64 arrows in the 8-by-8 grid going in random directions) for each session, and computed *S* _*c*_(random, model-free) for each random vector field.

In Extended Data Figure 16e, we estimated the autonomous dynamics vector field around the origin as a model-free way of confirming Extended Data Figure 16a. Similar to Extended Data Figure 16d, we convolved the spike trains with a Gaussian, and projected the baseline-subtracted firing rate trajectories to the low-dimensional subspace spanned by the FINDR latent axes. However, to separate the autonomous dynamics from the input dynamics, we used a Gaussian with a smaller σ (20ms), with the window size ±3σ around 0, and then excluded any ***ż***_*t*±3σ_ with time *t* where a click occurred, from the estimation of the autonomous dynamics. When computing the average of (***z***_*t*_ − ***z***_*t*−1_)/Δ*t* for one of the five pie slices, we required ***z***_*t*−1_ to be inside the pie slice. For all sessions, the circle was of radius 0.2 (in the units of ***z***). To further ensure that we estimate the autonomous dynamics, when computing the average, we only considered the trajectories where the number of left clicks was equal to the number of right clicks during the epoch when they were in the pie slice.

## Notes

### Competing Interest Statement

The authors have declared no competing interest.

### Summary of Updates

Updates to abstract, extended data figures, and supplementary materials.

## References

1. Wong K.-F. & Wang, X.-J. A recurrent network mechanism of time integration in perceptual decisions. J. Neurosci. 26, 1314–1328 (2006).

2. Dubreuil, A., Valente, A., Beiran, M., Mastrogiuseppe, F. & Ostojic, S. The role of population structure in computations through neural dynamics. Nat. Neurosci. 25, 783–794 (2022).

3. Bollimunta, A., Totten, D. & Ditterich, J. Neural dynamics of choice: single-trial analysis of decision-related activity in parietal cortex. J. Neurosci. 32, 12684–12701 (2012).

4. Latimer, K. W., Yates, J. L., Meister, M. L. R., Huk, A. C. & Pillow, J. W. NEURONAL MODELING. Single-trial spike trains in parietal cortex reveal discrete steps during decision-making. Science 349, 184–187 (2015).

5. Zoltowski, D. M., Latimer, K. W., Yates, J. L., Huk, A. C. & Pillow, J. W. Discrete Stepping and Nonlinear Ramping Dynamics Underlie Spiking Responses of LIP Neurons during Decision-Making. Neuron 102, 1249–1258.e10 (2019).

6. Mante, V., Sussillo, D., Shenoy, K. V. & Newsome, W. T. Context-dependent computation by recurrent dynamics in prefrontal cortex. Nature 503, 78–84 (2013).

7. Aoi, M. C., Mante, V. & Pillow, J. W. Prefrontal cortex exhibits multidimensional dynamic encoding during decision-making. Nat. Neurosci. 23, 1410–1420 (2020).

8. Yao, J. D. et al. Transformation of acoustic information to sensory decision variables in the parietal cortex. Proc. Natl. Acad. Sci. U. S. A. 120, e2212120120 (2023).

9. Khona, M. & Fiete, I. R. Attractor and integrator networks in the brain. Nat. Rev. Neurosci. 23, 744–766 (2022).

10. Inagaki, H. K., Fontolan, L., Romani, S. & Svoboda, K. Discrete attractor dynamics underlies persistent activity in the frontal cortex. Nature 566, 212–217 (2019).

11. Gardner, R. J. et al. Toroidal topology of population activity in grid cells. Nature 602, 123–128 (2022).

12. Sorscher, B., Mel, G. C., Ocko, S. A., Giocomo, L. M. & Ganguli, S. A unified theory for the computational and mechanistic origins of grid cells. Neuron 111, 121–137.e13 (2023).

13. Briggman, K. L., Abarbanel, H. D. I. & Kristan, W. B., Jr. Optical imaging of neuronal populations during decision-making. Science 307, 896–901 (2005).

14. Nieh, E. H. et al. Geometry of abstract learned knowledge in the hippocampus. Nature 595, 80–84 (2021).

15. Churchland, M. M. et al. Neural population dynamics during reaching. Nature 487, 51–56 (2012).

16. Kaufman, M. T., Churchland, M. M., Ryu, S. I. & Shenoy, K. V. Cortical activity in the null space: permitting preparation without movement. Nat. Neurosci. 17, 440–448 (2014).

17. Chaudhuri, R., Gerçek, B., Pandey, B., Peyrache, A. & Fiete, I. The intrinsic attractor manifold and population dynamics of a canonical cognitive circuit across waking and sleep. Nat. Neurosci. 22, 1512–1520 (2019).

18. Wang, X.-J. Decision making in recurrent neuronal circuits. Neuron 60, 215–234 (2008).

19. Machens, C. K., Romo, R. & Brody, C. D. Flexible control of mutual inhibition: a neural model of two-interval discrimination. Science 307, 1121–1124 (2005).

20. Wei, Z., Inagaki, H., Li, N., Svoboda, K. & Druckmann, S. An orderly single-trial organization of population dynamics in premotor cortex predicts behavioral variability. Nat. Commun. 10, 216 (2019).

21. Genkin, M., Shenoy, K. V., Chandrasekaran, C. & Engel, T. A. The dynamics and geometry of choice in premotor cortex. bioRxiv 2023.07.22.550183 (2023) doi:10.1101/2023.07.22.550183.

22. Rajan, K., Harvey, C. D. & Tank, D. W. Recurrent Network Models of Sequence Generation and Memory. Neuron 90, 128–142 (2016).

23. Prat-Ortega, G., Wimmer, K., Roxin, A. & de la Rocha J. Flexible categorization in perceptual decision making. Nat. Commun. 12, 1283 (2021).

24. Thura, D. & Cisek, P. Deliberation and commitment in the premotor and primary motor cortex during dynamic decision making. Neuron 81, 1401–1416 (2014).

25. Mohan, K., Zhu, O. & Freedman, D. J. Interaction between neuronal encoding and population dynamics during categorization task switching in parietal cortex. Neuron 109, 700–712.e4 (2021).

26. Galgali, A. R., Sahani, M. & Mante, V. Residual dynamics resolves recurrent contributions to neural computation. Nat. Neurosci. (2023) doi:10.1038/s41593-022-01230-2.

27. Finkelstein, A. et al. Attractor dynamics gate cortical information flow during decision-making. Nat. Neurosci. 24, 843–850 (2021).

28. Wimmer, K., Nykamp, D. Q., Constantinidis, C. & Compte, A. Bump attractor dynamics in prefrontal cortex explains behavioral precision in spatial working memory. Nat. Neurosci. 17, 431–439 (2014).

29. Pandarinath, C. et al. Inferring single-trial neural population dynamics using sequential auto-encoders. Nat. Methods 15, 805–815 (2018).

30. Schneider, S., Lee, J. H. & Mathis, M. W. Learnable latent embeddings for joint behavioural and neural analysis. Nature 617, 360–368 (2023).

31. Kim, T. D. et al. Flow-field inference from neural data using deep recurrent networks. bioRxiv (2023).

32. Ratcliff, R. & McKoon, G. The diffusion decision model: theory and data for two-choice decision tasks. Neural Comput. 20, 873–922 (2008).

33. Vickers, D. Evidence for an accumulator model of psychophysical discrimination. Ergonomics 13, 37–58 (1970).

34. Gold, J. I. & Shadlen, M. N. The neural basis of decision making. Annu. Rev. Neurosci. 30, 535–574 (2007).

35. Bogacz, R., Brown, E., Moehlis, J., Holmes, P. & Cohen, J. D. The physics of optimal decision making: a formal analysis of models of performance in two-alternative forced-choice tasks. Psychol. Rev. 113, 700–765 (2006).

36. Brunton, B. W., Botvinick, M. M. & Brody, C. D. Rats and humans can optimally accumulate evidence for decision-making. Science 340, 95–98 (2013).

37. Hyafil, A. et al. Temporal integration is a robust feature of perceptual decisions. Elife 12, (2023).

38. Kopec, C. D. et al. To integrate or not to integrate: Testing degenerate strategies for solving an accumulation of perceptual evidence decision-making task. bioRxiv (2024) doi:10.1101/2024.08.21.609064.

39. Erlich, J. C., Brunton, B. W., Duan, C. A., Hanks, T. D. & Brody, C. D. Distinct effects of prefrontal and parietal cortex inactivations on an accumulation of evidence task in the rat. Elife 4, (2015).

40. Hanks, T. D. et al. Distinct relationships of parietal and prefrontal cortices to evidence accumulation. Nature 520, 220–223 (2015).

41. Yartsev, M. M., Hanks, T. D., Yoon, A. M. & Brody, C. D. Causal contribution and dynamical encoding in the striatum during evidence accumulation. Elife 7, (2018).

42. Hunnicutt, B. J. et al. A comprehensive excitatory input map of the striatum reveals novel functional organization. Elife 5, (2016).

43. Anastasiades, P. G. & Carter, A. G. Circuit organization of the rodent medial prefrontal cortex. Trends Neurosci. (2021) doi:10.1016/j.tins.2021.03.006.

44. Sussillo, D., Jozefowicz, R., Abbott, L. F. & Pandarinath, C. LFADS - Latent Factor Analysis via Dynamical Systems. arXiv [cs.LG] (2016).

45. Keshtkaran, M. R. et al. A large-scale neural network training framework for generalized estimation of single-trial population dynamics. Nat. Methods 19, 1572–1577 (2022).

46. Weinan. A proposal on machine learning via dynamical systems. Commun. Math. Stat. 5, 1–11 (2017).

47. Chen, R. T. Q., Rubanova, Y., Bettencourt, J. & Duvenaud, D. K. Neural Ordinary Differential Equations. in Advances in Neural Information Processing Systems 31 (eds. Bengio, S.et al.) 6571–6583 (Curran Associates, Inc., 2018).

48. Kim, T. D., Can, T. & Krishnamurthy, K. Trainability, Expressivity and Interpretability in Gated Neural ODEs. in Proceedings of ICML (2023).

49. Newsome, W. T., Britten, K. H. & Movshon, J. A. Neuronal correlates of a perceptual decision. Nature 341, 52–54 (1989).

50. Kiani, R., Hanks, T. D. & Shadlen, M. N. Bounded integration in parietal cortex underlies decisions even when viewing duration is dictated by the environment. J. Neurosci. 28, 3017–3029 (2008).

51. Kiani, R. & Shadlen, M. N. Representation of confidence associated with a decision by neurons in the parietal cortex. Science 324, 759–764 (2009).

52. Inagaki, H. K. et al. Neural Algorithms and Circuits for Motor Planning. Annu. Rev. Neurosci. 45, 249–271 (2022).

53. Meister, M. L. R., Hennig, J. A. & Huk, A. C. Signal multiplexing and single-neuron computations in lateral intraparietal area during decision-making. J. Neurosci. 33, 2254–2267 (2013).

54. Park, I. M., Meister, M. L. R., Huk, A. C. & Pillow, J. W. Encoding and decoding in parietal cortex during sensorimotor decision-making. Nat. Neurosci. 17, 1395–1403 (2014).

55. Charlton, J. A. & Goris, R. L. T. Abstract deliberation by visuomotor neurons in prefrontal cortex. Nat. Neurosci. 27, 1167–1175 (2024).

56. Yates, J. L., Park, I. M., Katz, L. N., Pillow, J. W. & Huk, A. C. Functional dissection of signal and noise in MT and LIP during decision-making. Nat. Neurosci. 20, 1285–1292 (2017).

57. Neri, P., Parker, A. J. & Blakemore, C. Probing the human stereoscopic system with reverse correlation. Nature 401, 695–698 (1999).

58. Okazawa, G., Sha, L., Purcell, B. A. & Kiani, R. Psychophysical reverse correlation reflects both sensory and decision-making processes. Nat. Commun. 9, 3479 (2018).

59. Odoemene, O., Pisupati, S., Nguyen, H. & Churchland, A. K. Visual Evidence Accumulation Guides Decision-Making in Unrestrained Mice. J. Neurosci. 38, 10143–10155 (2018).

60. Panichello, M. F. & Buschman, T. J. Shared mechanisms underlie the control of working memory and attention. Nature 592, 601–605 (2021).

61. Roitman, J. D. & Shadlen, M. N. Response of neurons in the lateral intraparietal area during a combined visual discrimination reaction time task. J. Neurosci. 22, 9475–9489 (2002).

62. Stine, G. M., Trautmann, E. M., Jeurissen, D. & Shadlen, M. N. A neural mechanism for terminating decisions. Neuron (2023) doi:10.1016/j.neuron.2023.05.028.

63. Mochol, G., Kiani, R. & Moreno-Bote, R. Prefrontal cortex represents heuristics that shape choice bias and its integration into future behavior. Curr. Biol. (2021) doi:10.1016/j.cub.2021.01.068.

64. Brown, S. D. & Heathcote, A. The simplest complete model of choice response time: linear ballistic accumulation. Cogn. Psychol. 57, 153–178 (2008).

65. Thura, D., Beauregard-Racine, J.Fradet, C.-W. & Cisek, P. Decision making by urgency gating: theory and experimental support. J. Neurophysiol. 108, 2912–2930 (2012).

66. Inagaki, H. K. et al. A midbrain-thalamus-cortex circuit reorganizes cortical dynamics to initiate movement. Cell 185, 1065–1081.e23 (2022).

67. Charlton, J. A. & Goris, R. Abstract deliberation by visuomotor neurons in prefrontal cortex. bioRxiv 2022.12.06.519340 (2022) doi:10.1101/2022.12.06.519340.

68. Scott, B. B. et al. Fronto-parietal Cortical Circuits Encode Accumulated Evidence with a Diversity of Timescales. Neuron 95, 385–398.e5 (2017).

69. Shadlen, M. N. et al. Comment on ‘Single-trial spike trains in parietal cortex reveal discrete steps during decision-making’. Science vol. 351 1406–1406 Preprint at 10.1126/science.aad3242 (2016).

70. Gold, J. I., Law, C.-T., Connolly, P. & Bennur, S. The relative influences of priors and sensory evidence on an oculomotor decision variable during perceptual learning. J. Neurophysiol. 100, 2653–2668 (2008).

71. Urai, A. E., de Gee, J. W., Tsetsos, K. & Donner, T. H. Choice history biases subsequent evidence accumulation. Elife 8, (2019).

72. Mendonça, A. G. et al. The impact of learning on perceptual decisions and its implication for speed-accuracy tradeoffs. Nat. Commun. 11, 2757 (2020).

73. Daie, K., Fontolan, L., Druckmann, S. & Svoboda, K. Feedforward amplification in recurrent networks underlies paradoxical neural coding. bioRxiv (2023) doi:10.1101/2023.08.04.552026.

74. Goldman, M. S. Memory without feedback in a neural network. Neusron 61, 621–634 (2009).

75. Murphy, B. K. & Miller, K. D. Balanced amplification: a new mechanism of selective amplification of neural activity patterns. Neuron 61, 635–648 (2009).

## References

[1] Bingni W. Brunton, Matthew M. Botvinick, and Carlos D. Brody. Rats and humans can optimally accumulate evidence for decision-making. Science, 340(6128):95–98, 2013.

[2] Richard H. Byrd, Peihuang Lu, Jorge Nocedal, and Ciyou Zhu. A limited memory algorithm for bound constrained optimization. SIAM Journal on Scientific Computing, 16(5):1190–1208, 1995.

[3] Timothy Doyeon Kim, Tankut Can, and Kamesh Krishnamurthy. Trainability, Expressivity and Interpretability in Gated Neural ODEs. Proceedings of the 40th International Conference on Machine Learning, 2023.

[4] Timothy Doyeon Kim, Thomas Zhihao Luo, Tankut Can, Kamesh Krishnamurthy, Jonathan W. Pillow, and Carlos D. Brody. Flow-field inference from neural data using deep recurrent networks. bioRxiv, 2023.

[5] Diederik P Kingma and Max Welling. Auto-encoding variational bayes, 2014.

[6] Thomas Zhihao Luo, Adrian Gopnik Bondy, Diksha Gupta, Verity Alexander Elliott, Charles D. Kopec, and Carlos D. Brody. An approach for long-term, multi-probe neuropixels recordings in unrestrained rats. Elife, 2020.

[7] V. Mante, D. Sussillo, K. V. Shenoy, and W. T. Newsome. Context-dependent computation by recurrent dynamics in prefrontal cortex. Nature, 503(7474):78–84, 2013.

[8] Onyekachi Odoemene, Sashank Pisupati, Hien Nguyen, and Anne K. Churchland. Visual evidence accumulation guides decision-making in unrestrained mice. Journal of Neuroscience, 2018.

[9] Marius Pachitariu, Nick A. Steinmetz, Shabnam N. Kadir, Matteo Carandini, and Kenneth D. Harris. Fast and accurate spike sorting of high-channel count probes with kilosort. Advances in Neural Information Processing Systems, 29, 2016.

[10] Il Memming Park, Miriam L. Meister, Alexander C. Huk, and Jonathan W. Pillow. Encoding and decoding in parietal cortex during sensorimotor decision-making. Nature Neuroscience, 17, 2014.

[11] Neil C. Rabinowitz, Robbe L. Goris, Marlene Cohen, and Eero P. Simoncelli. Attention stabilizes the shared gain of v4 populations. Elife, 2015.

[12] Ryan J. Tibshirani and Bradley Efron. An Introduction to the Bootstrap. Chapman Hall/CRC, 1994.

