## Supplementary material for "Transitions in dynamical regime and neural mode underlie perceptual decision-making": Detailed descriptions of modeling procedures

### Supplementary Information

### 1 Simulations of dynamics consistent with existing hypotheses

The dynamical equations for each of the attractor hypotheses used to generate the flow fields in Figure 1 and Extended Data Figure 1, and synthetic datasets in Extended Data Figure 3 are as follows.

Bistable attractors:

$$\begin{aligned} dz_1 &= 10z_1(0.7 + z_1)(0.7 - z_1)dt + cudt + dW \\ dz_2 &= -10z_2dt + dW \end{aligned} \tag{1}$$

DDM line attractor:

$$
\begin{aligned}
dz_1 &= \begin{cases} cudt + 0.5dW, & z_1 \in (-0.7, 0.7) \\ 10z_1(0.7 - z_1)(0.7 + z_1)dt + 0.5dW, & z_1 \notin (-0.7, 0.7) \end{cases} \\
dz_2 &= -30z_2 + 0.5dW
\end{aligned} \tag{2}
$$

Non-normal, line attractor:

$$
\begin{aligned}
dz_1 &= 5z_2 + dW \\
dz_2 &= -5z_2dt + cudt + dW
\end{aligned} \tag{3}
$$

Here, $u$ at each time point can take a value from the set $\{-1, 0, 1\}$, where $-1$ indicates a leftward click, 0 indicates no click, and 1 indicates a rightward click. $c = 1/10$ for non-normal line attractor, and $c = 1/40$ for others. Similar to the task done by our rats, for each trial, $\gamma$ was drawn from the set $\{-3.5, -2.5, -1.5, -0.5, 0.5, 1.5, 2.5, 3.5\}$. We generated left and right clicks with rates $40/(1 + \exp(\gamma))$ and $40/(1 + \exp(-\gamma))$, respectively. The total duration of each trial was $1s$, and the duration of the click stimulus was randomly chosen to be one of $0.5s$, $0.7s$, and $0.9s$ for each trial. We generated 500 trajectories from each of Eq. (1–3), and assumed that the initial condition $\boldsymbol{z}(0) = \mathbf{0}$. Each trajectory corresponds to a trial. The spikes were generated from

$$
\begin{aligned}
W_{ij} &\sim 1 + 5\mathcal{N}(0, 1) \\
r_i &= \text{softplus}\left(\sum_j W_{ij}z_j + 5\right) \\
x_i &\sim \text{Poisson}(\Delta t \cdot r_i)
\end{aligned} \tag{4}
$$

where $\boldsymbol{W} \in \mathbb{R}^{80\times2}$, and $x_i$ is the number of spikes in a time bin with width $\Delta t = 0.01s$. Therefore, a total of 80 neurons and 500 trials were simulated for each dataset.

#### 2 Measurement model of spike counts

Poisson generalized linear models (GLM) are used as the measurement model of the spike counts in the FINDR model, the multi-mode drift-diffusion model (MMDDM), and in Extended Data Figure 14. For each neuron and on each time step $t$ with duration $\Delta t = 0.01$ s, number of spikes $y$ in the time interval $[t - \Delta t, t)$ is modelled as Poisson random variable:

$$
p(y \mid \lambda) = \frac{(\lambda\Delta t)^y \exp(-\lambda\Delta t)}{y!}. \tag{5}
$$

The firing rate $\lambda$ is the nonlinear output of a linear combination of inputs $x$

$$
\lambda = h\{x\} \tag{6}
$$

of the softplus function ($h$), which approximates the neuronal frequency-current curve of a neuron:

$$
h(x) = \log\{1 + \exp(x)\}. \tag{7}
$$

When the input $x$ is very negative, $h(x)$ is approximately zero. When $x$ is large and positive, $h(x)$ is approximately $x$, i.e., identity. The input $x$ varies on each time step $t$ on each trial and is a linear combination of the latent variable $z(t)$ and a putatively decision-independent baseline (or "bias") input $b(t)$

$$
x(t) = w_z \cdot z(t) + b(t). \tag{8}
$$

The bias $b$ incorporates nuisance variables as input to the neural spike trains and is the sum of a component that varies only across trials and a component that varies both across trials and within each trial:

$$
b(m, t) = b_{\text{cross}}(m) + b_{\text{within}}(m, t) \tag{9}
$$

where $m$ indicates trials and $t$ time step relative to stimulus onset.

#### 2.1 Measurement model in FINDR

In FINDR, the cross-trial component of the baseline of each neuron $n$ is a linear combination of the pre-trial activity of the entire recorded neuronal population:

$$b^{(n)}_{\text{cross}}(m) = w^{(n)}_c \cdot \left(\boldsymbol{q}^{(n)}\right)^\top \boldsymbol{p}_m \tag{10}$$

$\boldsymbol{p}_m$ is a vector whose element $i$ of the number of spikes emitted by neuron $i$ on the period of $[-2, 0)$ relative the beginning of the fixation period on trial $m$. The length of $\boldsymbol{p}$ is equal to the number of simultaneously recorded neuron and we include all neurons in $\boldsymbol{p}$ regardless of choice selectivity, as well neuron $n$ itself. The vector $\boldsymbol{q}$ is the solution to a regularized linear regression problem

$$\boldsymbol{q}^{(n)} = (\boldsymbol{P}^\top \boldsymbol{P} + \beta \boldsymbol{I})^{-1} \boldsymbol{P}^\top \boldsymbol{r} \tag{11}$$

where $\boldsymbol{P}$ concatenates the pre-trial spike count across trials

$$\boldsymbol{P} = \begin{bmatrix} \boldsymbol{p}_1 & \boldsymbol{p}_2 & \cdots & \boldsymbol{p}_M \end{bmatrix}$$

and $\boldsymbol{r}$ is the overall firing rate averaged across the epoch of the trial from stimulus onset to the earlier of either 1 s after stimulus onset or immediately before the animal leaves the fixation port. $\boldsymbol{I}$ is the identity matrix, and we choose the L2 regularization penalty $\beta$ over a grid of values $\beta \in$ {1e0, 1e2, ... 1e10} using five-fold cross-validation.

Having learned $\boldsymbol{q}$, the scalar weight $w_c$ is learned together with the parameters of other component of the baseline $b_{\text{within}}$, by maximizing the marginal likelihood of a Poisson generalized linear model of each individual neuron's spike count. On time step $t$ of trial $m$, the firing rate of neuron $n$ is given by

$$\lambda^{(n)}(m, t) = h\left\{b^{(n)}_{\text{cross}}(m) + b^{(n)}_{\text{within}}(m)\right\} \tag{12}$$

The within-trial component is a sum of the filtered output of an impulse occurring at each of the following events in the trial: onset of the auditory click trains and the initiation of the animal's movement away from the fixation.

$$b_{\text{within}}(m, t) = \sum_{\text{event}} \tau^{(m)}_{\text{event}} (k_{\text{event}} * \delta)(t) \tag{13}$$

where the symbol $*$ indicates convolution, $\tau_x$ indicates translation $\tau_x k(t) = k(t - \tau_x)$ by the time of event $x$, $\delta$ is the Dirac delta function. For each type of event we estimate a linear filter, or kernel, $k_{\text{event}}$, parametrized as the linear combination of a set of smooth temporal basis functions:

$$k_{\text{event}} = \Phi_{\text{event}} \boldsymbol{w}_{\text{event}} \tag{14}$$

where each $\Phi_i$ is a $T \times D$ temporal basis matrix representing $D$ orthogonal temporal basis functions (see section 5), and $\boldsymbol{w}_i$ is a $D$-dimensional vector of weights that are learned from the data.

**Post-stimulus onset kernel** The beginning of each trial and the onset of the auditory click trains is indicated by an auditory click played simultaneously by the left and right speaker, which we refer to as a "stereoclick." The kernel extends from 0-1.0 s after the stereoclick and is parameterized using $D = 8$ basis functions with a time-warping parameter $\eta_{warp} = 1$ (see section 5). A larger $\eta_{warp}$ indicates a greater over-representation of the time close to the event at the expense of under-representation of time far from the event.

**Pre-movement kernel** This kernel extends from -1.0 s to -0.01 s relative to when the animal departs from the fixation port and is parameterized by $D = 3$ temporal basis functions time warping $\eta_{warp} = 0.1$. No choice-related or evidence-related information is included in this kernel.

On time step $t$ of trial $m$, the firing rate is given by

$$\lambda(m, t) = h\left\{w_c \cdot \boldsymbol{q}^\top \boldsymbol{p}_m + \tau^{(m)}_{\text{stim}} (\Phi_{\text{stim}} \boldsymbol{w}_{\text{stim}} * \delta)(t) + \tau^{(m)}_{\text{move}} (\Phi_{\text{move}} \boldsymbol{w}_{\text{move}} * \delta)(t)\right\} \tag{15}$$

where $\boldsymbol{w} = (w_c, \boldsymbol{w}_{\text{stim}}, \boldsymbol{w}_{\text{stim}})$ are free parameters. We place an isotropic Gaussian prior on the $\boldsymbol{w}$ with precision parameter $\alpha$ (functionally equivalent to L2 regularization). We learn both $\boldsymbol{w}$ and $\alpha$ directly from the data by maximizing the log of the marginal likelihood

$$\boldsymbol{w}^* = \arg\max_{\boldsymbol{w}} \sum_m \sum_t \log \int p(y\{m,t\} \mid \boldsymbol{w})\, p(\boldsymbol{w} \mid \alpha)\, d\boldsymbol{w} \tag{16}$$

Because the integral has no closed form (Poisson and Gaussian are not conjugate distributions), this is done using a variational approach described in [14].

During cross-validation, not only $\boldsymbol{w}$ but also $\boldsymbol{q}$ is learned using only training trials. To learn $\boldsymbol{q}$, each set of training trials is partitioned again into training trials and test trials.

### 3 Multi-mode drift-diffusion model (MMDDM)

#### 3.1 Dynamic model

Following previous work [3, 4], we formulate the dynamics of $z$ as an Ornstein-Uhlenbeck process, whose time evolution can be represented using the Fokker-Planck equation

$$dz = \begin{cases} (\lambda z + u)dt + \sigma_z dW, & z \in (-B, B) \\ 0, & \text{otherwise} \end{cases} \tag{17}$$

When $z$ is at either the left bound $-B$ or the right $B$, its value does not change. The bound height parameter $B \in (5, 20)$ is learned from the data.

The hyperparameter $\lambda$ quantifies the consistent drift in $z$. For $\lambda < 0$, the integration is leaky, $z$ drifts toward 0, and early inputs affect $z$ less than later inputs. For $\lambda > 0$, $z$ positively feedbacks to itself, $z$ is impulsive, and early inputs affect $z$ more than later inputs. The time constant of $z$ is $1/\lambda$. In the case of $\lambda = 0$, $z$ behaves in the regime of the drift-diffusion model. To implement either the MMDDM or the single mode DDM, we set $\lambda$ to be 0. To implement a leaky integrator, we fit $\lambda$ but constrain it to be negative, and to implement an impulsive integrator, we fit $\lambda$ but constrain it to be positive.

The momentary input $u(t)$ is the total difference in the per-click input between the right and left clicks that occurred in the time interval $[t - \Delta t, t)$

$$u(t) = \sum_{\tau \in R} v(\tau; t) - \sum_{\tau \in L} v(\tau; t) \tag{18}$$

where $L(R)$ is the set of the left (right) click times and $v(\tau; t)$ is the per-click input of a click emitted at time $\tau$ on the latent variable at time step $t$. Note that $\tau \in \mathbb{R}$ indicates continuous time, whereas $t \in \mathbb{N}$ indexes a time step. The per-click input is given by

$$v(\tau; t) = D(\tau; t) \cdot C(\tau) \cdot \zeta \tag{19}$$

where $D(\tau; t)$ indicates the integral over the interval $[t - \Delta t, t)$ of the Direct delta function $\delta$ delayed by $\tau$:

$$D(\tau; t) = \int_{t-\Delta t}^{t-\epsilon} \delta(x - \tau)\, dx = \begin{cases} 1, & \tau \in [t - \Delta t, t) \\ 0, & \text{otherwise} \end{cases} \tag{20}$$

where $\epsilon$ is the machine epsilon. To account for sensory adaptation, the per-click input is depressed by preceding clicks by a time-varying scaling factor given by the function $C(\tau)$ (see section 3.1.4).

The per-click input is corrupted by i.i.d. multiplicative Gaussian noise $\zeta$:

$$\zeta \sim \mathcal{N}(1, \sigma_s^2) \tag{21}$$

The free parameter $\sigma_s^2 \in (0.1, 20)$ quantifies the noise-to-signal ratio of the sensory input and is optimized during model fitting.

In the absence of feedback ($\lambda = 0$) and no momentary input, the variable $z$ evolves stochastically due to the diffusion noise $\sigma_z dW$, where $dW$ is a white noise Weiner process. Following previous findings [3], diffusion noise is fixed to be small by setting $\sigma_z = 1$ to be a hyperparameter.

##### 3.1.1 Transition probability

The solution to the Fokker-Planck equations in Eq. 17 provides the transition probability of $z$:

$$p(z_t \mid z_{t-1}) = \begin{cases} 1, & z_{t-1} = z_t = -B \\ 1, & z_{t-1} = z_t = B \\ f(z_t \mid \mu_t, \sigma_t^2), & (z_{t-1}, z_t) \in (-B, B) \\ \int_{-\infty}^{-B} f(x \mid \mu_t, \sigma_t^2)\, dx, & z_t = -B, z_{t-1} \in (-B, B) \\ \int_{B}^{\infty} f(x \mid \mu_t, \sigma_t^2)\, dx, & z_t = B, z_{t-1} \in (-B, B) \\ 0, & \text{otherwise} \end{cases} \tag{22}$$

If $z$ is at either of the two absorbing bounds (i.e., $-B$ or $B$) in the previous time step, then $z$ must retain the same value for the current time step. If both the previous and present values are within the bounds, the probability is given by a Gaussian distribution with mean $\mu_t$ and variance $\sigma_t^2$:

$$f(x \mid \mu_t, \sigma_t^2) \sim \mathcal{N}(\mu_t, \sigma_t^2) \tag{23}$$

$$\mu_t = e^{\lambda dt}\left(z_{t-1} + \frac{v_t}{\lambda dt}\right) - \frac{v_t}{\lambda dt} \tag{24}$$

where $v$ is used here to indicate the expectation of the difference between the sum of the mean of the right and left click inputs:

$$v = \sum_{\tau_R \in [t-dt, t)} C(\tau_R) - \sum_{\tau_L \in [t-dt, t)} C(\tau_L) \tag{25}$$

In the limit of $\lambda$ toward 0, the mean $\mu_t$ is given by

$$\lim_{\lambda \to 0} \mu_t = z_{t-1} e^{\lambda dt} + v_t \left(1 - e^{-\lambda dt}\right) \tag{26}$$

The variance $\sigma_t^2$ is a sum of the stimulus-independent diffusion noise and the noise emanating from each click

$$\sigma_t^2 = \sigma_z^2 dt + \sigma_s^2 \left[ \sum_{\tau_R \in [t-dt, t)} C(\tau_R) + \sum_{\tau_L \in [t-dt, t)} C(\tau_L) \right] \tag{27}$$

##### 3.1.2 Prior probability

The prior probability is given by

$$p(z_1) = \begin{cases} f(z_1 \mid \mu_1, \sigma_1^2), & B > z_1 > -B \\ \int_{-\infty}^{-B} f(x \mid \mu_1, \sigma_1^2)\, dx, & z_1 = -B \\ \int_{B}^{\infty} f(x \mid \mu_1, \sigma_1^2)\, dx, & z_1 = B \end{cases} \tag{28}$$

Where

$$f(x \mid \mu_1, \sigma_1^2) \sim \mathcal{N}(\mu_0, \sigma_0^2) \tag{29}$$

The mean of the Gaussian distribution is specified by the parameter $\mu_0 \in (-5, 5)$, which is learned from the data, and the variance is a hyperparameter fixed to $\sigma_0^2 = 1$.

##### 3.1.3 Discretization of time and decision variable

Similar to previous studies[3, 4], the model is implemented by discretizing both time and the decision variable $z$. Time is discretized to $\Delta t = 0.01$s, and the value of $z$ is discretized to $\Xi = 53$ values equally spaced in $[-B, B]$. The i-th bin $\xi_i$ is given by

$$\xi_i = \frac{2i - \Xi - 1}{\Xi - 1} B \tag{30}$$

The difference between successive bins is

$$\Delta\xi = \frac{2B}{\Xi - 1} \tag{31}$$

Under this discretization scheme, the probability $p(z)$ at each time step is represented by a $\Xi$-element probability vector $\boldsymbol{p}$ whose first element indicates the probability of $z$ at the left bound, the last element the probability at the right bound, and the intervening elements correspond to the approximate probability at each bin. By definition of being of a probability vector, the elements of $\boldsymbol{p}$ always sum to 1. For each value $z$, the probability $p(z)$ is divided between the two closest bins, weighted by the relative distance between $z$ and the two bins.

**Prior probability vector** To illustrate the discretization in detail, below we demonstrate the computation of the prior probability vector. Each element of the probability vector is computed as a definite integral of the normal distribution in Eq. (28). For the first element of the prior probability vector, which quantifies the probability of $z$ at the left bound:

$$\boldsymbol{p}_{i=1} = \int_{-\infty}^{\xi_1} f(x;\mu_1,\sigma_1^2)\,dx + \frac{1}{\Delta\xi}\left[\int_{\xi_1}^{\xi_2} f(x;\mu_1,\sigma_1^2)\,(\xi_2 - x)\,dx\right] \tag{32}$$

The definite integral is equal to

$$\boldsymbol{p}_{i=1} = \Phi(\bar{z}_1) + \frac{\sigma_1}{\Delta\xi}\left\{\bar{z}_2\left[\Phi(\bar{z}_2) - \Phi(\bar{z}_1)\right] + \phi(\bar{z}_2) - \phi(\bar{z}_1)\right\} \tag{33}$$

where $\bar{z}_i$ is the z-score

$$\bar{z}_i = \frac{\xi_i - \mu_1}{\sigma_1}$$

and $\Phi(x)$ and $\phi$ are the standard normal cumulative distribution function and standard normal probability density function, respectively. The mean $\mu_1$ and the standard deviation $\sigma_1$ are parameters of the model. For the second to second-to-last elements of the probability vector, i.e., $i \in \{2, \ldots \Xi - 1\}$:

$$\boldsymbol{p}_i = \frac{\sigma_1}{\Delta\xi}\left\{\bar{z}_{i+1}\left[\Phi(\bar{z}_{i+1}) - \Phi(\bar{z}_i)\right] + \bar{z}_{i-1}\left[\Phi(\bar{z}_{i-1}) - \Phi(\bar{z}_i)\right] + \phi(\bar{z}_{i+1}) + \phi(\bar{z}_{i-1}) - 2\phi(\bar{z}_i)\right\} \tag{34}$$

Finally, for the last element:

$$\boldsymbol{p}_\Xi = 1 - \Phi(\bar{z}_\Xi) + \frac{\sigma_1}{\Delta\xi}\left\{\bar{z}_{\Xi-1}\left[\Phi(\xi_{\Xi-1}) - \Phi(\xi_\Xi)\right] + \phi(\bar{z}_{\Xi-1}) - \phi(\bar{z}_\Xi)\right\} \tag{35}$$

**Stochastic matrix** The transition probability of $z$ in Eq. (22) is represented as a square $\Xi \times \Xi$ stochastic matrix:

$$Z_t \approx p(z_t \mid z_{t-1}) \tag{36}$$

Each column of the matrix corresponds to the probability vector corresponding to the conditional probability given the value of $z$ on the preceding time step:

$$Z_t[:, j] \approx p(z_t \mid z_{t-1} = \xi_j) \tag{37}$$

Each element of the transition matrix is the conditional probability

$$Z_t[i, j] \approx p(z_t = \xi_i \mid z_{t-1} = \xi_j) \tag{38}$$

The first column of the transition matrix represents the conditional transition probability given that $z$ was at the bound on the previous time step. By the definition of the absorbing bounds, for all time step $t$, the first column of the transition matrix has a value of 1 at the first element and zero elsewhere.

$$Z[i, 1] = \begin{cases} 1, & i = 1 \\ 0, & \text{otherwise} \end{cases} \tag{39}$$

Similarly, the last column has a value of 1 at the last element and zero elsewhere:

$$Z[i, \Xi] = \begin{cases} 1, & i = \Xi \\ 0, & \text{otherwise} \end{cases} \tag{40}$$

Each of the intervening columns is computed similarly to the prior probability vector, but with a different Gaussian mean $\mu_j$ and variance $\sigma^2$. For $j \in \{2, ...\Xi - 1\}$. Both $\mu_j$ and $\sigma^2$ vary across time steps, but the subscript $t$ is omitted for clarity.

$$Z_t[i,j] = \begin{cases} \Phi(\bar{z}_1) + \frac{\sigma_t}{\Delta\xi}\left(\bar{z}_2\left[\Phi(\bar{z}_2) - \Phi(\bar{z}_1)\right] + \phi(\bar{z}_2) - \phi(\bar{z}_1)\right), & i = 1 \\ \frac{\sigma_t}{\Delta\xi}\left(\bar{z}_{i+1}\left[\Phi(\bar{z}_{i+1}) - \Phi(z_i)\right] + z_{i-1}\left[\Phi(\bar{z}_{i-1}) - \Phi(\bar{z}_i)\right] + \phi(\bar{z}_{i+1}) + \phi(\bar{z}_{i-1}) - 2\phi(\bar{z}_i)\right), & \Xi > i > 1 \\ 1 - \Phi(\bar{z}_\Xi) + \frac{\sigma_t}{\Delta\xi}\left(z_{\Xi-1}\left[\Phi(\xi_{\Xi-1}) - \Phi(\xi_\Xi)\right] + \phi(\bar{z}_{\Xi-1}) - \phi(\bar{z}_\Xi)\right), & i = \Xi \end{cases} \tag{41}$$

where

$$\bar{z}_i = \frac{\xi_i - \mu_j}{\sigma}$$

The mean of the Gaussian is given by

$$\mu_j = e^{\lambda\Delta t}\left(\xi_j + \frac{v}{\lambda\Delta t}\right) - \frac{v}{\lambda\Delta t} \tag{42}$$

where $v$ is the expectation of the difference between the sum of the right and left click inputs:

$$v = \sum_{\tau_R \in [t-\Delta t, t)} C(\tau_R) - \sum_{\tau_L \in [t-\Delta t, t)} C(\tau_L). \tag{43}$$

$\tau_R$ are the times when a right click occurred, $\tau_L$ where a left click occurred, and the $C(t)$ the post-adaptation strength of each click (see section 3.1.4).

The variance $\sigma^2$ does not vary across the columns of the transition matrix and a sum of the stimulus-independent diffusion noise and the noise emanating from each click

$$\sigma^2 = \sigma_z^2 \Delta t + \sigma_s^2 \left[\sum_{\tau_R \in [t-\Delta t, t)} C(\tau_R) + \sum_{\tau_L \in [t-\Delta t, t)} C(\tau_L)\right] \tag{44}$$

Because the momentary click input varies across time steps, and because both $\mu_j$ and $\sigma^2$ depend on the momentary input.

##### 3.1.4 Sensory adaptation

Sensory adaptation is conceptualized as nearly full depression immediately after a click and gradually less and less depression after the preceding click. This idea is modeled with the ordinary differential equation parametrizing the gain $C$ of the momentary input

$$\frac{dC}{dt} = C \cdot (\phi - 1) \cdot \delta(\boldsymbol{\tau}_{L,R} - t) + k(1 - C) \tag{45}$$

with initial condition

$$C(t = 0) = 1 \tag{46}$$

At the occurrence of the first click on each trial, which is always the stereoclick (i.e., a click played simultaneously from the left and right speakers), the gain of the momentary input is given by

$$C(dt) = \phi$$

In accordance with previous findings [3] and the behavioral and neural phenomena associated with the precedence effect[2], the hyperparameter $\phi$ is fixed to be 0.001. After the occurrence of a click, in the gap of time before the subsequent click, the gain recovers toward 1 with the exponential change rate $k$. We specify the change to be fast by setting $k = 100$.

After each click at time $\tau$, the dynamics of $C$ at time $\tau+\Delta t$ before any other click's arrival is computed by separately integrating over the intervals $[\tau, \tau+\epsilon]$ and $(\tau+\epsilon, \tau+\Delta t]$ for small $\epsilon \ll \Delta t$:

$$
\begin{aligned}
\int_{\tau}^{\tau+\epsilon} dC &= k\int_{\tau}^{\tau+\epsilon}[1-C(t)]dt + \int_{\tau}^{\tau+\epsilon}(\phi-1)\delta(\tau-t)C(t)dt \\
C(\tau+\epsilon) - C(\tau) &\approx k[1-C(\tau+\epsilon)-1+C(\tau)]\epsilon + (\phi-1)C(\tau) \\
C(\tau+\epsilon) &\approx C(\tau) + k\epsilon[C(\tau)-C(\tau+\epsilon)] + (\phi-1)C(0) \\
C(\tau+\epsilon) &\approx C(\tau)\left[\phi + k\epsilon\left(1-\frac{C(\tau+\epsilon)}{C(\tau)}\right)\right]
\end{aligned}
$$

Integrating over the second interval,

$$
\begin{aligned}
&\int_{\tau+\epsilon}^{\tau+\Delta t}\frac{dC}{1-C(t)} = \int_{\tau+\epsilon}^{\tau+\Delta t}kdt + \int_{\tau+\epsilon}^{\tau+\Delta t}(\phi-1)\delta(\tau-t)\frac{C(t)}{1-C(t)}dt \\
&-\log(|1-C(\tau+\Delta t)|) + \log(|1-C(\tau+\epsilon)|) = k(\Delta t-\epsilon)
\end{aligned}
$$

Using the result obtained by integrating over the first interval,

$$
\begin{aligned}
|1-C(\tau+\Delta t)| &= \exp\left(\log(|1-C(\tau+\epsilon)|) - k(\Delta t-\epsilon)\right) \\
|1-C(\tau+\Delta t)| &= |1-C(\tau+\epsilon)|\exp\left(-k(\Delta t-\epsilon)\right) \\
|1-C(\tau+\Delta t)| &\approx \left|1-C(\tau)\left[\phi+k\epsilon\left(1-\frac{C(\tau+\epsilon)}{C(\tau)}\right)\right]\right|\exp\left(-k(\Delta t-\epsilon)\right)
\end{aligned}
$$

In the limit of $\epsilon \to 0$,

$$
|1-C(\tau+\Delta t)| = |1-\phi C(\tau+\Delta t)|\exp\left(-k\Delta t\right)
$$

Therefore,

$$
C(\tau+\Delta t) = \begin{cases} 1-[1-\phi C(\tau)]\exp\left(-k\Delta t\right) \\ 1+[1-\phi C(\tau)]\exp\left(-k\Delta t\right) \end{cases}
$$

These two possibilities are disambiguated by the requirements

$$
\begin{aligned}
&k \geq 0 \\
&\phi \in [0,1] \\
&C \in [0,1] \\
&C(0) = 1
\end{aligned}
$$

Therefore

$$
C(\tau+\Delta t) = 1-[1-\phi C(\tau)]\exp\left(-k\Delta t\right) \tag{47}
$$

Note that if we take the derivative, we recover

$$
\frac{dC}{dt} = k(1-C)
$$

For click times $\boldsymbol{\tau}$, the dynamics of $C$ at the moment of each click can be computed recursively:

$$
\begin{aligned}
C(\tau_1) &\equiv 1 \\
C(\tau_2) &= 1-[1-\phi]\exp\left[-k(\tau_2-\tau_1)\right] \\
C(\tau_3) &= 1-[1-\phi C(\tau_2)]\exp\left[-k(\tau_3-\tau_2)\right] \\
&\ldots
\end{aligned}
$$

##### 3.1.5 Latency

The clicks have a latency of $\Delta t$=0.01 s, set based on the latency of auditory responses in rat primary auditory cortex [15]. On time step $t$, the latent decision variable $z$ receives input from clicks that occurred during $[t-2\Delta t, t-\Delta t)$. Results are not affected by the exact value of the latency.

#### 3.2 Measurement model of the spike trains

Conditioned on the decision variable $z$, each neuron's spike train response $y$ is modeled as Poisson random variable:

$$p(y) = (\lambda \Delta t)^y \exp(-\lambda \Delta t)/y! \tag{48}$$

The spike train is binned at the time steps of $\Delta t = 0.01$ s, the same as that of the latent variable $z$. The firing rate $\lambda$ is the nonlinearly rectified output of the softplus function ($h$), which approximates the neuronal frequency-current curve of a neuron:

$$h(x) = \log\{1 + \exp(x)\} \tag{49}$$

When the input $x$ is very negative, $h(x)$ is approximately zero. When $x$ is large and positive, $h(x)$ is approximately $x$ itself. The same activation function is used for all neurons.

##### 3.2.1 Decision-dependent predictor

The input $x$ varies on each time step $t$ on each trial and is a linear combination of the latent variable $z(t)$ and decision-independent baseline input $b(t)$

$$x(t) = w_z \cdot \tilde{z}(t) + b(t) \tag{50}$$

To maximize interpretability and minimize trade-offs between parameters, the value of $z$ being encoded is independent of the bound height:

$$\tilde{z}_i \equiv \frac{z_i}{B} = \frac{2i - \Xi - 1}{\Xi - 1} \tag{51}$$

The weight of $z$ depends on the state $i$ of $z$ itself:

$$w_z = \begin{cases} w_{DC} & \text{if } z \text{ is at a bound, i.e., } i = 1 \text{ or } i = \Xi \\ w_{EA} & \text{otherwise} \end{cases} \tag{52}$$

Each neuron has its own scalar $w_{DC}$ and $w_{EA}$. Importantly, the neurons do not receive any direct input from the auditory clicks. The decision-independent baseline input $b(t)$ is a weighted sum of decision-independent predictors.

##### 3.2.2 Within-trial varying component of the baseline

The decision-independent predictors consist of four types of variables, three of which vary both within and across trials, and a fourth that varies only across trials.

Each within-trial varying predictor is aligned to an event in a trial: 1) the stereoclick (a click played from the left and right speakers simultaneously), which defines the start of auditory click trains and the trial itself; 2) the animal's departure from the center port, which ends a trial if it occurs before 1s from the stereoclick; and 3) a neuron's own previous spiking.

On each trial $m$, an impulse input occurs at the time of each event, convolved with a filter, or kernel, $k$ specific to that event.

$$b_{\text{within}}(m, t) = \sum_{\text{event}} \tau_{\text{event}}^{(m)} (k_{\text{event}} * \delta)(t) \tag{53}$$

where the symbol $*$ indicates convolution, $\tau_x$ indicates translation $\tau_x k(t) = k(t - \tau_x)$ by the time of event $x$, $\delta$ is the Dirac delta function. For each type of event, we estimate a linear filter, or kernel, $k_{\text{event}}$, parametrized as the linear combination of a set of smooth temporal basis functions:

$$k_{\text{event}} = \Phi_{\text{event}} \boldsymbol{w}_{\text{event}} \tag{54}$$

where each $\Phi_i$ is a $T \times D$ temporal basis matrix representing $D$ orthogonal temporal basis functions (see section 5), and $\boldsymbol{w}$ is a $D$-dimensional vector of weights that are learned from the data.

**Post-stereoclick kernel** The beginning of each trial and the onset of the auditory click trains is indicated by an auditory click played simultaneously by the left and right speaker, i.e., a stereoclick. The kernel $k$ extends from 0-1.0 s after the stereoclick and is parameterized using $D = 5$ basis functions with time warping $\eta_{\text{warp}} = 0.1$ (see section 5). A positive $\eta_{warp}$ indicates a greater over-representation of the time close to the event at the expense of under-representation of time far from the event.

**Pre-movement kernel** This kernel extends from 0.6 s to 0.01 s before the animal departs from the center port and is parameterized by $D = 2$ temporal basis functions with time warping $\eta_{\text{warp}} = 0.1$

**Post-spike kernel** This kernel extends from 0.01-0.25 s after each spike emitted by the same neuron (but not by other neurons) and is parametrized by $D = 3$ temporal basis functions with a time-warping $\eta_{warp} = 1$.

###### 3.2.3 Cross-trial component of the baseline

The cross-component of the baseline is parameterized as the linear combination of eight smooth temporal basis functions evaluated over the start times of each trial $m$. To reduce the number of parameters fit in the MMDDM, the relative weights of up to $D = 8$ smooth temporal basis functions $\phi$ are first learned using a separate linear-Gaussian model fit to the firing rates $r$ on each trial:

$$r \sim \mathcal{N}(\mu, \sigma) \tag{55}$$

$$\mu \sim \sum_{i}^{D} \phi_i(\tau_m) \tag{56}$$

#### 3.3 Measurement model of the behavioral choice

On each trial, the binary behavioral choice $c$ (1=right, 0=left) is the sign of $z$ on the trial's last time step $T$ (the earlier of 1 s after the onset of the clicks or immediately before the animal leaves the fixation port):

$$c \mid z(T) = \text{sign}\Big\{z(T)\Big\} \tag{57}$$

#### 3.4 Parameter estimation

By discretizing the value of $z$, the MMDDM has the form of an autoregressive, input-output hidden Markov model (HMM) [1]. The "autoregressive" feature refers to the inputs from each neuron's own spike history to capture the long-range correlations between spike train observations of the same neuron, and the "input-output" feature points to the dependence of the spike train observations on multiple baselines that vary on different time scales.

The inference of the most likely parameters of a model of this structure given simultaneously observed spike trains can be efficiently accomplished using the expectation-maximization (EM) algorithm [5]. However, the EM algorithm exhibits slow, first-order convergence when the conditional distributions of the observations, given the latent state, are not well-separated across states [12]. For the MMDDM, a poor separation between the conditional spike train distributions is expected from the approximation of $z$ as many discrete states. Therefore, instead of the EM algorithm, we use the closely related expectation-conjugate-gradient (ECG) algorithm, which shows superior performance for poorly separated conditional distributions [12]. The ECG algorithm computes the exact gradient of the log-likelihood of the observations by solving the expectation step of the standard forward-backward algorithm for HMMs. The gradient can be used by any first-order optimization algorithm to maximize the log-likelihood. The first-order optimization algorithm used here is the limited-memory BFGS [8], which is a quasi-Newton method that efficiently approximates the hessian matrix using the gradients.

###### 3.4.1 Gradient of the log-likelihood

On each trial, the observations on which the likelihood of the MMDDM parameters are computed consist of the behavioral choice $c$:

$$c = \begin{cases} 0, & \text{left choice} \\ 1, & \text{right choice} \end{cases} \tag{58}$$

and the spike trains of $N$ simultaneously recorded neurons:

$$\boldsymbol{Y} = \{y_1, y_2, ...y_N\} \tag{59}$$

$$y = [y_1, y_2, ...y_T]^\top. \tag{60}$$

The spike train of each neuron on each time step $y_{n,t}$ is the spike count of that neuron over $[t, t + \Delta t)$. The gradient of the log-likelihood of the simultaneously recorded spike trains $\boldsymbol{Y}$ and the choice $d$ on each trial is given by

$$\begin{aligned}
\nabla \log p(\boldsymbol{Y}, c) &= \frac{\nabla p(\boldsymbol{Y}, c)}{p(\boldsymbol{Y}, c)} \\
&= \frac{\nabla \sum_{\boldsymbol{z}} p(\boldsymbol{Y}, c, \boldsymbol{z})}{p(\boldsymbol{Y}, c)} \\
&= \frac{\sum_{\boldsymbol{z}} p(\boldsymbol{Y}, c, \boldsymbol{z}) \nabla \log p(\boldsymbol{Y}, c, \boldsymbol{z})}{p(\boldsymbol{Y}, c)} \\
&= \sum_{\boldsymbol{z}} p(\boldsymbol{z} \mid \boldsymbol{Y}, c) \nabla \log p(\boldsymbol{Y}, c, \boldsymbol{z}) \\
&= \sum_{\boldsymbol{z}} p(\boldsymbol{z} \mid \boldsymbol{Y}, c) \left[\nabla \log p(\boldsymbol{Y}, c \mid \boldsymbol{z}) + \nabla \log p(\boldsymbol{z})\right]
\end{aligned} \tag{61}$$

where $\boldsymbol{z}$ is the set of the values of the latent decision variable on each of the $T$ time steps of the trial:

$$\boldsymbol{z} = \{z_1, z_2, ..., z_T\}$$

and conditioning jointly over the latent variable on all time steps. However, because of the first-order Markov dependence, the gradient can be simplified further:

$$\begin{aligned}
\nabla \log p(\boldsymbol{Y}, c) = &\sum_{z_1} p(z_1 \mid \boldsymbol{Y}, c) \nabla \log p(z_1) + \\
&\sum_{t=2}^{T} \sum_{z_t} p(z_t, z_{t-1} \mid \boldsymbol{Y}, c) \nabla \log p(z_t \mid z_{t-1}) + \\
&\sum_{t=2}^{T} \sum_{z_t} p(z_t \mid \boldsymbol{Y}, c) \sum_{n=1}^{N} \nabla \log p(y_{n,t} \mid z_{t-1}) + \\
&\sum_{z_T} p(z_T \mid \boldsymbol{Y}, c) \nabla \log p(c \mid z_T).
\end{aligned} \tag{62}$$

The posterior probabilities $p(z_t \mid \boldsymbol{Y}, c)$ and $p(z_t, z_{t-1} \mid \boldsymbol{Y}, d)$ were computed using the forward-backward algorithm, which can be used even when the spike trains have spike history inputs [5].

##### 3.5 Relationship between MMDDM and rSLDS

The MMDDM can approximately be described in the framework of recurrent switching linear dynamical systems (rSLDS) [6, 16], a model that approximates nonlinear dynamics with a discrete set of linear regimes. The rSLDS framework distinguishes between continuous variables, which we denote here as $\boldsymbol{z}$, and discrete states, which we denote as $\boldsymbol{x}$. The first dimension of the continuous variable $\boldsymbol{z}$ is identical to the one-dimensional decision variable in MMDDM. The second dimension of $\boldsymbol{z}$ is close to zero except when the first dimension of $\boldsymbol{z}$ reaches a bound $B$. The two discrete states $\boldsymbol{x} \in \{[1; 0], [0; 1]\}$ correspond to the dynamical regimes of evidence accumulation ($\boldsymbol{x} = [1; 0]$) and decision commitment ($\boldsymbol{x} = [0; 1]$):

$$\boldsymbol{x}_t \mid \boldsymbol{x}_{t-1}, \boldsymbol{z}_{t-1} = \begin{cases} [1; 0], & \|z_{1,t-1}\| < B \\ [0; 1], & \|z_{1,t-1}\| \geq B \end{cases}. \tag{63}$$

Here

$$\boldsymbol{z}_t = \begin{cases} A_1 \boldsymbol{z}_{t-1} + V_1 \boldsymbol{u}_t + \epsilon_{1,t}, & \boldsymbol{x}_t = [1; 0] \\ A_2 \boldsymbol{z}_{t-1} + V_2 \boldsymbol{u}_t + \epsilon_{2,t}, & \boldsymbol{x}_t = [0; 1] \end{cases}, \tag{64}$$

where $A_1 = [1, 0; 0, 1]$, $V_1 = [1, -1; 0, 0]$, $\epsilon_{1,t} \sim N(0, [\sigma^2, 0; 0, 10^{-6}])$, $A_2 = [1, 0; 1, 0]$, $V_2 = [0, 0; 0, 0]$, $\epsilon_{2,t} \sim N(0, [10^{-6}, 0; 0, 10^{-6}])$. $10^{-6}$ is some arbitrarily small number.

The initial condition of the $\boldsymbol{z}$ variable is set to be $\boldsymbol{z}_0 = [0, 0]$. If there is a bias $\beta$ in the decision process, then the initial state is $[\beta, 0]$. The initial condition of the $\boldsymbol{x}$ variable is $[1; 0]$, indicating that the starting regime is always evidence accumulation and not decision commitment.

The mapping from the variable $z$ to the firing rates $\boldsymbol{r}$ of a population of $N$ neurons is given by

$$\boldsymbol{r} = \text{softplus}\left(\boldsymbol{W}z + \boldsymbol{b}\right) \tag{65}$$

$\boldsymbol{W}$ is a $N \times 2$ matrix in which the first row represents the encoding weights in the evidence accumulation regime ($w_{EA}$) and the second row the encoding weights in decision commitment minus the encoding weights in the evidence accumulation regime ($w_{DC} - w_{EA}$). The $N \times 1$ vector $\boldsymbol{b}$ represents the decision-irrelevant input, including spike history.

### 4 Psychophysical kernel model

To test the predictions of MMDDM, we inferred the psychophysical kernel, which quantifies the time-varying weight of the stimulus input on the behavioral choice.

The traditional approach to infer the psychophysical kernel is to use the reverse correlation technique [7], a method highly similar to the spike-triggered average [13]. However, the traditional reverse correlation technique critically depends on the stimulus fluctuations being independent across time. The traditional approach can be biased by inputs that are correlated in time and also biased by stimulus-independent factors that influence the decision but cannot be incorporated in the analysis [10]. Moreover, the traditional technique typically assume a single temporal resolution for the weights of the stimulus fluctuations. When the assumed temporal resolution exceeds what can inferred from the data, the inferred kernel can be highly noisy and has limited interpretability.

We developed a new model, based on logistic regression, to infer the psychophysical kernel that mitigates the bias from sensory inputs with temporally-correlated stimulus fluctuations and the bias from omitted factors that impact the choice. The psychophysical kernel is parametrized using temporal basis functions, and therefore the temporal resolution of the kernel is quantified by the number of basis functions. The optimal number of basis functions for a given dataset can be identified using out-of-sample model comparison.

To mitigate the bias from temporally correlated stimulus fluctuations, we incorporate a method based on the logistic regression model presented in [9] that reparametrizes the inputs to remove the temporal correlations in the stimulus input. To mitigate the potential bias from omitted factors, a behavioral lapse parameter is incorporated. To identify the optimal temporal resolution of the psychophysical kernel that can be inferred using a given data set, the kernel is parametrized as a linear combination of smooth temporal basis functions. Cross-validated model comparison identifies the optimal temporal resolution.

#### 4.1 Model definition

The behavioral choice $c$ is modelled as a Bernoulli random variable. The probability of a right choice is given by

$$p(c = 1) = f(\beta_1)f(\beta_2) + [1 - f(\beta_1)]f(y) \tag{66}$$

where the function $f$ is the logistic function

$$f(x) = \frac{1}{1 + \exp(-x)} \tag{67}$$

which monotonically maps the input $x \in (-\infty, \infty)$ to the range $(0, 1)$. The value $f(\beta_1)$ quantifies the behavioral lapse rate, or the fraction of trials in which the animal is a behavioral state that ignores the stimulus and depends on only its bias $\beta_2$. In the remaining $1 - f(\beta_1)$ of the trials, the choice depends on not only the bias $\beta_2$ but also the sensory stimulus. The linear combination of these inputs is given by

$$y = \beta_2 + \beta_3 \lambda T \Delta t + \boldsymbol{e}^\top \Phi \boldsymbol{w} \tag{68}$$

The parameter $\beta_3$ is the weight of the expected stimulus input at each time step ($\Delta t = 0.01$), which is represented as $\lambda T \Delta t$. The independent variable $\lambda$ is the expected stimulus input per second given the random processes used to generate the stimuli on each trial. The hyperparameter $T$ indicates the number of time steps on each trial examined ($T = 40$ when aligned to the time of decision commitment inferred from MMDDM and $T = 75$ when aligned to stimulus onset). When $\lambda$ does not equal 0, the stimulus fluctuations are temporally correlated across time steps within a trial. Therefore, separate weights are learned for the expected stimulus input and the fluctuations from the expected input.

The expected input rate $\lambda$ is the difference between the values of right and left input rate expected given the random process used to generate the trial, and the typical expected input rates are $\lambda \in \{-38, -34, -25, -10, 10, 25, 34, 38\}$

Fluctuations from the expected input are quantified as the excess stimulus input $\boldsymbol{e}$. The $T$ dimensional vector $\boldsymbol{e}$ is the difference between the actual and the expected input at each time step $\tau$

$$e(t) = -\lambda \Delta t + \sum_{\tau \in [t-\Delta t, t)} \delta_R(\tau) - \delta_L(\tau) \tag{69}$$

Where $\delta_R(\tau)$ equals 1 when a right click occurred at time $\tau$ and is zero otherwise, and $\delta_L(\tau)$ is similarly defined for left clicks.

The weights of the excess stimulus input is given by the vector $\Phi\boldsymbol{w}$. The matrix $\Phi$ is $T \times D$. Each row of the matrix $\Phi$ corresponds to one of $T$ time step in the trial, and each column consists of the output of one $D$ basis functions (see section 5). The weight vector $\boldsymbol{w}$ has the dimensionality $D$ corresponding to the number of the temporal basis functions, and the values of the vector are learned from the data. When $D = 1$, $\Phi$ is constrained to be a vector whose elements are the identical, and the weights $\Phi\boldsymbol{w}$ is constrained to be a flat line.

#### 4.2 Inference

The free parameters of the model are $\{\beta_1, \beta_2, \beta_3, \boldsymbol{w} \in \mathbb{R}^D\}$. The hyperparameter $D$, which quantifies the number of the temporal basis functions and therefore represents the time resolution of the psychophysical kernel, is optimized by 10-fold cross-validated model comparison.

#### 4.3 Implementation details

##### 4.3.1 Figure 3m

The results shown in Figure 3m are inferred without any basis function. Only trials for which the auditory click trains preceded the moment of out-of-sample MMDDM-inferred decision commitment time by at least 0.2 s and also continued for another 0.2 s were included. In the shuffling procedure, the inferred time of decision commitment was randomly permuted across trials. Then, within this permutation sample, we selected trials for which the time of decision commitment preceded the end of the auditory click trains by at least 0.2 s to be used to compute the psychophysical kernel in the "shuffled" condition.

##### 4.3.2 Extended Data Figure 10b-d

The results shown in these panels were from inference using basis functions. Similar to the analysis for Figure 3m, we included only trials for which the auditory click trains both preceded and continued to play after the moment of out-of-sample MMDDM-inferred decision commitment time by at least 0.2 s.

##### 4.3.3 Extended Data Figure 10e-h

To estimate the psychophysical kernel aligned to trial onset (i.e., the start of the auditory click trains), trials with a stimulus duration of at least 0.75s were included from the set of recording sessions used in other analyses.

### 5 Basis functions

In either the multi-mode drift-diffusion model (MMDDM) and the psychophysical kernel model, nonlinear basis functions of the independent variables are used to reduce the number of parameters needed to be learned. For a given independent variable whose values vary across $T$ time steps on each trial, we did not learn the weight at each time step. Instead, we learn the weight of each of $D$ basis functions of the independent variable. Because $D << T$, the number of parameters can be greatly reduced.

For a given independent variable, such as the time-varying fluctuations in auditory click input in the psychophysical kernel model, its value on a given trial with $T$ time steps can be expressed as the $T$-dimensional input vector $\boldsymbol{x}$. The basis function output $\boldsymbol{y}$ of the independent variable is given by

$$\boldsymbol{y} = \Phi^\top \boldsymbol{x} \tag{70}$$

The matrix $\Phi$ has the dimensions $T \times D$: each column corresponds to an individual basis function, and each row to a time step. The basis function output $y$ is therefore a vector of length $D$.

The linear combination of $\boldsymbol{y}$ is an input in a pre-nonlinearity stage of either the MMDDM or the PKM. For example, in the MMDDM, the Poisson rate of a neuron $\boldsymbol{\lambda}$ is given by

$$\boldsymbol{\lambda} = h\left(\boldsymbol{w}_1^\top \boldsymbol{y}_1 + \boldsymbol{w}_2^\top \boldsymbol{y}_2 + ...\right) \tag{71}$$

where $h$ is the softplus function. The linear projection weights $\boldsymbol{w}_i$ are learned for each of the $i$-th independent variable.

The basis function matrix $\Phi$ is pre-specified and fixed for each independent variable and for all trials. Each column of $\Phi$ consists of a radial basis function evaluated on the index $\tau$ of each time step. The radial basis function we used has the form of a raised cosine [11]. The j-th basis function is given by

$$\phi_j(\tau) = \begin{cases} 1/2 - \cos\left\{\theta_j(\tau)\right\}/2, & \theta \in [0, 2\pi] \\ 0, & \text{otherwise} \end{cases} \tag{72}$$

where the transformation from time step index to radian is given by

$$\theta_j(\tau) = \frac{[f(\tau) - f(1) - (j-3)\Delta]\pi}{2\Delta} \tag{73}$$

The function $f$ is a monotonic time warping function described below. The value $\Delta$ is the distance between the peaks of two adjacent basis functions:

$$\Delta = \frac{f(T) - f(1)}{D - 1} \tag{74}$$

After evaluating $\Phi$, it is transformed into a unitary matrix using singular vector decomposition.

#### 5.1 Time warping

To parametrize the post-spike kernel in the MMDDM or the peri-commitment psychophysical kernel, we wish to parametrize the portion of the kernel closer to the event of interest with a higher density of basis functions than the portion of the kernel farther away from the event time. To implement a continuous change in the density of representation, the time step indices are monotonically transformed using the function $f$

$$f(\tau; \eta) = \sinh^{-1}\{\eta_{warp}(\tau - \tau_0)\} \tag{75}$$

The time index when the event occurred is indicated by $\tau_0$. The parameter $\eta$ indicates the degree of time warping. The larger is the value of $\eta$, the greater the warping: i.e., the portion of the kernel closer to $\tau_0$ is represented by more functions and can have a higher temporal resolution than the portion father away from $\tau_0$. For $1 >> \eta > 0$, the warping is essentially absent, and $f(\tau; \eta_{warp}) \approx \tau$. This can be seen using the first order Taylor approximation of $\text{asinh}^{-1}(\eta_{warp}\tau)$ around zero

$$f(\eta\tau) \approx f(0) + f'(0) \cdot \eta_{warp}\tau = \eta_{warp}\tau$$

### References

[1] C. M. Bishop. *Pattern Recognition and Machine Learning*. Springer Science & Business Media, 2006.

[2] A. D. Brown, G. C. Stecker, and J. T. Daniel. The precedence effect in sound localization. *Journal of the Association for Research in Otolaryngology*, 16, 2015.

[3] B. W. Brunton, M. M. Botvinick, and C. D. Brody. Rats and humans can optimally accumulate evidence for decision-making. *Science*, 340(6128):95–98, 2013.

[4] B. DePasquale, J. W. Pillow, and C. D. Brody. Neural population dynamics underlying evidence accumulation in multiple rat brain regions. *bioRxiv*, 2021.

[5] S. Escola, A. Fontanini, D. Katz, and L. Paninski. Hidden markov models for the stimulus-response relationships of multistate neural systems. *Neural Computation*, 23, 2011.

[6] S. W. Linderman, M. J. Johnson, A. C. Miller, R. P. Adams, D. M. Blei, and L. Paninski. Bayesian Learning and Inference in Recurrent Switching Linear Dynamical Systems. In *Proceedings of the 20th International Conference on Artificial Intelligence and Statistics*, volume 54, pages 914–922, 2017.

[7] P. Neri, A. J. Parker, and C. A. Blakemore. Probing the human stereoscopic system with reverse correlation. *Nature*, 1999.

[8] J. Nocedal and S. J. Wright. *Numerical Optimization*. Springer Science & Business Media, 2000.

[9] O. Odoemene, S. Pisupati, H. Nguyen, and A. K. Churchland. Visual evidence accumulation guides decision-making in unrestrained mice. *Journal of Neuroscience*, 2018.

[10] G. Okazawa, L. Sha, B. A. Purcell, and R. Kiani. Psychophysical reverse correlation reflects both sensory and decision-making processes. *Nature Communications*, 2018.

[11] I. M. Park, M. L. Mester, A. C. Huk, and J. W. Pillow. Encoding and decoding in parietal cortex during sensorimotor decision-making. *Nature Neuroscience*, 17, 2014.

[12] R. Salakhutdinov, S. Roweis, and Ghahramani Z. Optimization with em and expectation-conjugate-gradient. *Proceedings of the 20th International Conference on Machine Learning*, 2003.

[13] O. Schwartz, J. W. Pillow, N. C. Rust, and E. P. Simoncelli. Spike-triggered neural characterization. *Journal of Vision*, 2006.

[14] A. Wu, N. A. Roy, S. Keeley, and J. W. Pillow. Gaussian process based nonlinear latent structure discovery in multivariate spike train data. *Advances in Neural Information Processing Systems*, 30, 2017.

[15] J. D. Yao, P. Bremen, and J. C Middlebrooks. Rat primary auditory cortex is tuned exclusively to the contralateral hemifield. *Journal of Neurophysiolology*, 110, 2013.

[16] D. M. Zoltowski, J. W. Pillow, and S. W. Linderman. A general recurrent state space framework for modeling neural dynamics during decision-making. In *Proceedings of the 37th International Conference on Machine Learning*, volume 119, pages 11680–11691, 2020.
